# Plant Polymerase IV sensitizes chromatin through histone modifications to preclude spread of silencing into protein-coding domains

**DOI:** 10.1101/2021.08.25.457601

**Authors:** Vivek Hari Sundar G, Chenna Swetha, Debjani Basu, Kannan Pachamuthu, Steffi Raju, Tania Chakraborty, Rebecca A. Mosher, P. V. Shivaprasad

## Abstract

Across eukaryotes, gene regulation is manifested via chromatin states roughly distinguished as heterochromatin and euchromatin. The establishment, maintenance and modulation of the chromatin states is mediated using several factors including chromatin modifiers. However, factors that avoid the intrusion of silencing signals into protein coding genes are poorly understood. Here we show that a plant specific paralogue of RNA polymerase (pol) II, named pol IV, is involved in avoidance of facultative heterochromatic marks in protein coding genes, in addition to its well-established functions in silencing repeats and transposons. In its absence, H3K27 trimethylation (me3) mark intruded the protein coding genes, more profoundly in genes embedded with repeats. In a subset of genes, spurious transcriptional activity resulted in small(s) RNA production leading to post-transcriptional gene silencing. We show that such effects are significantly pronounced in rice, a plant with larger genome with distributed heterochromatin when compared to Arabidopsis. Our results indicate the surprising division of labour among plant-specific polymerases, not just in establishing effective silencing via sRNAs and DNA methylation, but also in influencing chromatin boundaries.

## Introduction

The genetic material DNA in eukaryotes is compacted as chromatin by means of association with histones and other proteins (Kornberg 1974; Luger et al. 1997). The accessibility of the DNA for various molecular processes such as transcription, repair and replication is dynamically modulated in distinct domains of the chromatin, largely categorized into fairly accessible gene-rich euchromatin and condensed heterochromatin (Feng and Michaels 2015; Brown 1966). The chromatin is decorated with modifications to DNA and histones constituting epigenetic modifications (Feng and Michaels 2015; Law and Jacobsen 2010). Proportion of genomic regions coding for proteins called euchromatin reduces with increase in genome size across organisms and such alterations and genome expansion is mainly due to the proliferation of repeats and transposons (Pellicer et al. 2018). Plant genomes are especially enriched with latent transposons poised for transcription located proximal to the protein coding genes in the euchromatic domains (Hirsch and Springer 2017).

The activity of RNA polymerase II (pol II) on the euchromatic domains is regulated locally by transcription factors, chaperones and chromatin remodelling enzymes that transduces the local epigenetic status (Schier and Taatjes 2020; Hahn 2004; Gibney and Nolan 2010). Evolution of plants with increased instances of gene-proximal repeats in the genome necessitates articulation of silencing states to the pol II with precision. This locally silenced state is called facultative heterochromatin and isenriched with Polycomb Repressive Complex (PRC) dependent H3K27me3 marks in contrast to the constitutive heterochromatin enriched with H3K9me2 modifications (Zhang et al. 2007). Studies in early land plant genomes like that of *Marchantia polymorpha* suggest that H3K27me3 evolved as a predominant silencing mark in plants (Montgomery et al. 2020) and H3K9me2 mark took over the constitutive silencing leading to the observed dichotomy between facultative and constitutive heterochromatin (Déléris et al. 2021). Mutants of MET1, a major player in constitutive heterochromatin establishment, exhibited compensation by H3K27me3 marks at repeats and transposons (Deleris et al. 2012; Soppe et al. 2002; Mathieu et al. 2005; Rougée et al. 2021), indicating that unknown players monitor heterochromatin states in specific domains to initiate compensatory marks in constitutive heterochromatic regions. Demarcation of facultative and constitutive heterochromatic boundaries is paramount in avoiding the intrusion of silencing states into neighbouring protein coding genes, warranting evolution of novel machineries curtailing silencing overshoot and simultaneously defending against genotoxic repeats. Although these mechanisms have been envisaged across eukaryotes, major upstream players involved in these beyond the obvious epigenetic readers, writers and erasers, are unknown.

Plants have evolved RNA silencing as an efficient mode of robust and targeted silencing of repeats and genes, both at transcriptional and post-transcriptional levels (Baulcombe 2004). Small RNAs (sRNAs), predominant effectors of plant RNA silencing, confer both specificity and amplification modality in silencing. Production of sRNAs associated with transcriptional silencing are primarily initiated by plant specific RNA polymerase II (pol II) paralog RNA polymerase IV (pol IV) and in peculiar cases by pol II itself (Nuthikattu et al. 2013; Cuerda-Gil and Slotkin 2016). The pol IV transcripts, majorly originating from the repeats and transposons are acted upon by RNA DEPENDENT RNA POLYMERASE2 (RDR2) while the pol II aberrant transcripts are acted upon by RNA DEPENDENT RNA POLYMERASE6 (RDR6), thereby converting them to double stranded duplexes that become substrates for several DICER-LIKE proteins (DCLs) (Nuthikattu et al. 2013). The resultant short duplex sRNAs are picked up by specific ARGONAUTE (AGO) effectors, specified by their length and one of the strands possessing the preferred 5’-nucleotide (nt) (Mi et al. 2008). For post transcriptional gene silencing (PTGS), majorly undertaken in plants by cleavage of target mRNA, 21-22nt size class sRNAs with 5’-Uracil (U) are loaded into the AGO1 leading to slicing of the target mRNA between positions 10 and 11 in the sRNA:mRNA duplex regions (Baumberger and Baulcombe 2005). On the contrary, for transcriptional silencing pol IV derived repeat associated 24nt sRNAs are loaded into AGO4 (Zilberman et al. 2003) with a 5’-Adenosine (A) preference and this ribonucleoprotein complex binds to the complementary region of the long scaffold transcript produced by RNA polymerase V (pol V) (Wierzbicki et al. 2008). This triggers recruitment of DNA methyl transferases like DOMAINS REARRANGED METHYLTRANSFERASE2 (DRM2) and majorly result in asymmetric DNA methylation at the target locus (Matzke and Mosher 2014). Several other pathways culminating in production of sRNAs, for example, viral transcription, transgene induced and endogenous aberrant transcription, the pol II-RDR6 aberrant activity feedback to the similar silencing pathway with identical end results (Elmayan and Vaucheret 1996; Herr et al. 2005; Voinnet et al. 1998; Nuthikattu et al. 2013). Numerous non-canonical modalities in the concerted activity of these RNA polymerase variants have also been reported (Cuerda-Gil and Slotkin 2016).

Even though RNA silencing via sRNAs is an effective process to accurately delimit silencing, the loci over which different polymerases operate are dependent on the chromatin states. For instance, the pol IV is recruited by the readers of H3K9me2 marks named SAWADEE HOMEODOMAIN HOMOLOG1 and 2 (SHH1 and 2) in coordination with CLASSY type chromatin remodellers (Law et al. 2013; Zhang et al. 2013; Zhou et al. 2018). The pol V recruitment involves DNA methylation readers like SUVH2 and SUVH9 (Liu et al. 2014), chromatin remodelling complex called as DDR complex (Law et al. 2010) and several other proteins that are modulated by chromatin states. The activity of these polymerases is tightly linked to the epigenetic status of the locus of interest and hence the RNA silencing. Few reports have probed the effects of loss of DNA methylation machineries on the chromatin states in Arabidopsis and observed excessive reorganisation of chromatin domains. For example, in *met1* mutants chromatin is decondensed with loss of clear heterochromatin and euchromatin boundaries, exemplifying the impact of the cross talk between these epigenetic states (Soppe et al. 2002; Mathieu et al. 2005; Zhong et al. 2021). Similarly, *pol iv* exhibits permanent loss of silencing at selected loci that are recalcitrant to rescue by complementation of pol IV and this phenomenon is due to associated loss of H3K9me2 marks (Li et al. 2020). Unrestricted to silencing, it is also evidenced that loss of pol IV leads to transcriptional mis-regulation as observed by the accumulation of atypical nascent transcripts in maize (Erhard et al. 2015) and by increased pol II activity at 3’-ends of pol II transcriptional units (McKinlay et al. 2018). The indirect facilitation of spurious transcription by virtue of loss of epigenetic players triggers cryptic transcription units by pol II both in genes and transposons (Le et al. 2020). Prolific presence of repeat fragments in the gene coding units like introns in rice have shown to trigger transcriptional silencing like features (Espinas et al. 2020). Taken together, epigenetic landscape modulates not only the silencing of repeats directly but also encumbers the cryptic transcriptional activity.

Such counter-balancing reinforcement loops between epigenetic states must contribute to molecular and morphological phenotypes upon perturbation, especially in monocots with higher proportion of transposons. It has been conclusively documented that reproductive structures in rice, including gametes, undergo massive reprogramming in terms of sRNA production and DNA methylation in a locus-specific manner (Chenxin et al. 2020). Indeed, unlike Arabidopsis, loss of silencing players display exacerbated phenotypes in the monocot model rice in the cases of *drm2*, *met1b, pol iv, pol v* and *dcl3* (Moritoh et al. 2012; Yamauchi et al. 2014; Zheng et al. 2021; Wei et al. 2014). In agreement with this, grass family (*Poaceae*) members have evolved specific neo-functionalized RNA polymerase paralog called RNA polymerase VI, functions of which are still unclear (Trujillo et al. 2018). These phenomena substantiate that complex gene arrangement interleaved by repeats mandates very robust and agile mechanism of epigenetic silencing in a very localised manner.

In this study, we resorted to examine the genome and phenome level functions of RNA pol IV in rice by generating precise knockdown (*kd*) mutants and validated them by genetic complementation. We present evidence for over-shoot of heterochromatic H3K27me3 marks into the protein coding regions in the *kd* and redistribution of the H3K9me2 marks. A subset of coding genes became hotspots of aberrant sRNA production. We show that these aberrant sRNAs from atypical loci in *kd* are present in loss of function mutants of Arabidopsis counterparts, selective only to pol IV complex assembly mutants. We provide evidence that these sRNAs induce PTGS of protein coding genes especially in complex genome like rice. Together, these results indicate the presence of several novel features of regulation of plant genomes, including intrusion of H3K27me3 in constitutive heterochromatin and role of pol IV in preventing heterochromatinization of protein-coding genes. This acts as a strong negative selection on pol IV-sRNA modules, absence of which can trigger PTGS of important genes in a species-specific manner.

## Results

### Knockdown of RNA polymerase IV induced pleiotropic phenotypic defects

The loss of function of the catalytically active component of the polymerase IV complex, NRPD1, causes a spectrum of effects in different plant species – from delayed growth transition in *Physcomitrella patens,* delayed flowering in *Arabidopsis thaliana,* reproductive defects in *Brassica rapa, Capsella rubella and Zea mays,* to increased tillering in *Japonica* rice (Coruh et al. 2015; Wang et al. 2020b; Kf et al. 2009; Grover et al. 2018). In order to understand the multifaceted roles of polymerase IV beyond RdDM in rice, we generated artificial microRNA (amiR)-mediated knockdown (*kd*) transgenic *indica* rice lines targeting the largest subunit of RNA pol IV (NRPD1). Since rice genome encodes two isoforms of NRPD1 – NRPD1a and NRPD1b, sharing 88.8% of sequence identity, amiR was designed to target a conserved sequence in the N-terminal region of both the transcripts, incorporating optimal design parameters as put forth earlier (Fig. 1A) (Narjala et al. 2020). Previous attempts targeted both the rice NRPD1 isoforms using RNA interference of the C-terminal region with sub-optimal targeting or the CRISPR-Cas9 mediated knockout (KO) that was reported as embryo lethal or showing drastic reduction in fertility (Xu et al. 2020; Debladis et al. 2020; Zheng et al. 2021; Zhang et al. 2020). In order to get a precise targeting and to avoid the lethality and sterility effects of KO, we resorted to the amiR technology. In order to unequivocally prove that amiR is targeting only NRPD1 leading to the effects, we super-transformed the *kd* with a construct expressing amiR-targeting resistant version of NRPD1b CDS driven by its cognate promoter (Fig. 1A). The expression of mature amiR was confirmed with northern hybridisation and the *kd* lines showed reduction in transcript abundance of both the isoforms with 70 percent restoration in the complementation (CO) (Figs. 1B, C and Supplemental Figs. S1A, B). Rice transformation was performed twice independently to score for consistent phenotypes and to eliminate T-DNA insertion effects, independent transgenic events were identified by junction fragment Southern analyses (Supplemental Figs. S2A, B).

**Figure 1.**
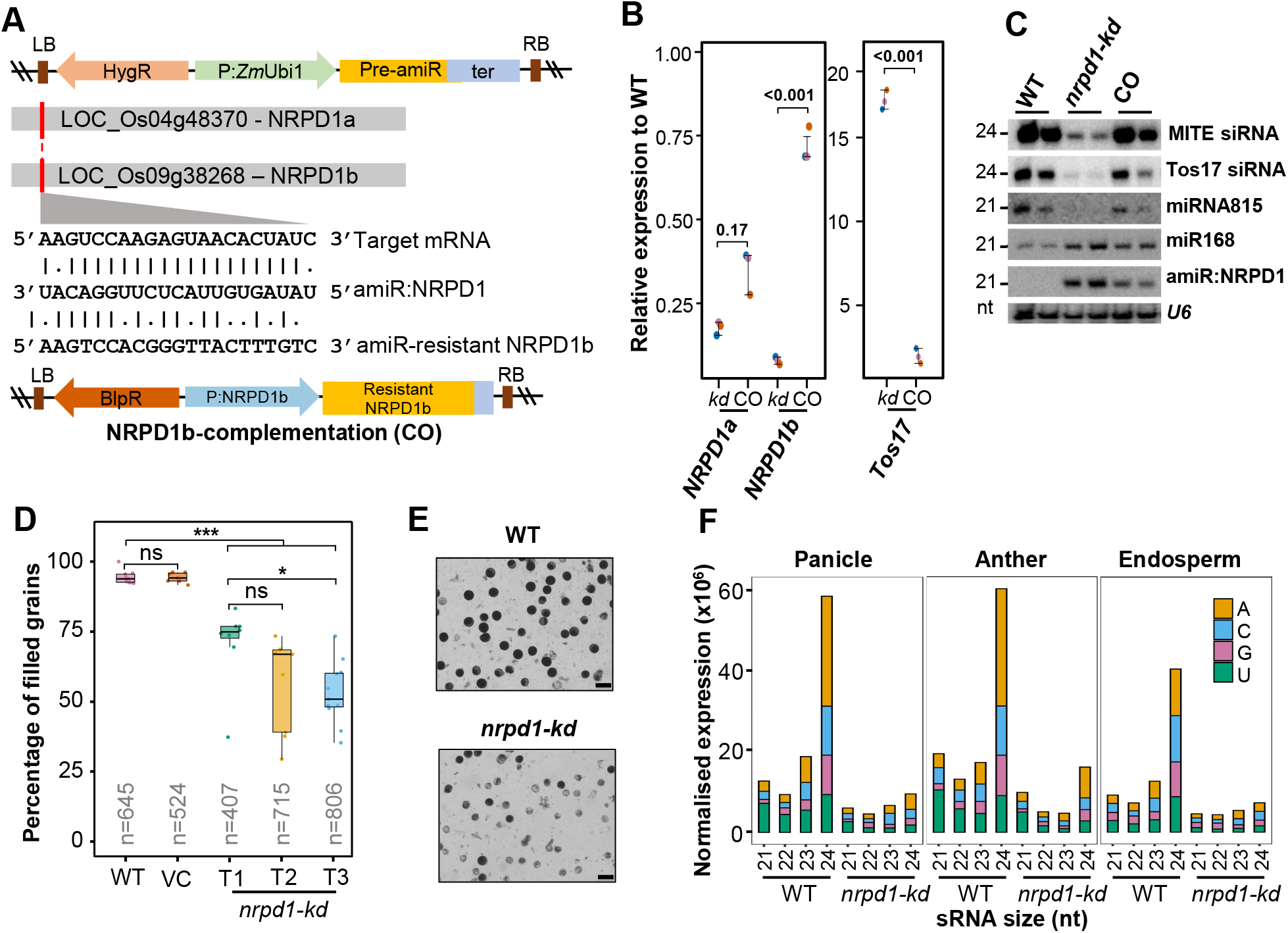
Knockdown of RNA pol IV in rice results in pleiotropic phenotypes. *(A)* T-DNA map of amiR coding binary construct with the super-transformed NRPD1b complementation (CO) construct. The alignment depicts the amiR binding region and the modifications in amiR-resistant CDS of NRPD1b that was driven by its cognate promoter (P:NRPD1b). BlpR – bialaphos resistance. The precursor-amiR (Pre:amiR) is driven by maize ubiquitin promoter (P:*Zm*Ubi1). ter – 35S-poly(A) signal; HygR – Hygromycin selection marker; RB and LB – right and left border. *(B)* Plots representing relative abundance of *NRPD1a, NRPD1b* and *Tos17* transcripts in *kd* and CO with respect to WT, measured by RT-qPCRs. Student’s t-test; p-values mentioned across comparisions. *(C)* sRNA northern blots showing the accumulation of 24nt siRNAs and miRNAs in WT, *kd* and CO. *U6* was used as loading control. *(D)* Boxplots showing the percentage of filled grains (n=number of grains). Dots represent average of panicles in each plant (Tukey’s test; ***p-value of 0.001, *p-value of 0.01, ns-non-significant). *(E)* Representative images of pollen grains of dehisced anther stained with iodine. Scale: 50 μm. *(F)* Stacked bar plots showing normalised abundance of sRNAs of different sizes with their 5’ nucleotides. The replicates were merged.

As observed in previous studies (Xu et al. 2020), the *kd* plants showed increased tiller number (Supplemental Fig. S2C). Strikingly, the percentage of viable filled grains were significantly low at 75% in T1 *kd* plants and diminished progressively over generations to reach 50% in T3 (Fig. 1C and Supplemental Fig. S1D). The unfilled florets showed structural deformities and they were consistent among the distinct transgenic events proving that the defects observed were not due to transgenesis (Supplemental Figs. S2C-E). Specific lineages of the *kd* lines had extreme reproductive defects without viable grains (Supplemental Fig. S3A). Southern and northern analyses showed these defects are not due to the zygosity of the T-DNA or the dosage of amiR in such a lineage indicating possible effect of induced epimutations (Supplemental Figs. S3B, C) (Johannes and Schmitz 2019). Since NRPD1 is reported to influence pollen development (Wang et al. 2020b), we further investigated the pollen quality by measuring size, iodine staining potential and ultrastructural characterisation using scanning electron microscopy that revealed pollen defects in *kd* (Figs. 1D, E and Supplemental Figs. S1E-G). Taken together, amiR mediated *kd* of RNA polymerase IV complex in rice led to severe reproductive defects consistent across generations in distinct transgenic lines similar to monocots such as maize (Kf et al. 2009), unlike mild abnormalities observed in *Arabidopsis*.

### Rice pol IV is required for biogenesis of repeat associated sRNAs aiding their silencing

In order to explore the molecular effects of loss of polymerase IV in rice and to discern the reasons for extreme reproductive defects upon NRPD1 *kd*, we performed total sRNA profiling in three reproductive tissues that exhibited drastic defects – pre-emerged panicle, anther and endosperm. As shown earlier in other species, *kd* plants showed evident reduction of 24nt sRNAs in all the tissues (Figs. 1C, F and Supplemental Fig. S4A) (Herr et al. 2005; Wang et al. 2020b; Coruh et al. 2015; Grover et al. 2018). As expected (Mi et al. 2008; Zhai et al. 2015), there was a loss of 5’-A containing sRNAs in all the three tissues (Fig. 1F). On the contrary, bulk of the pol II transcribed miRNAs and the sRNAs mapping to their precursors largely remained unchanged (Supplemental Figs. S4B, C). Further, as the pol IV derived sRNAs are known to be associated with transposons and repeats, we profiled the sRNAs from major annotated repeats in the rice genome. We observed substantial loss of repeat associated sRNAs and this was also validated by northern blotting in different tissues (Fig. 1C and Supplemental Fig. S4D-F). Notably, the repeat associated sRNAs were also restored to WT levels in the complementation lines as proven by northern blots (Fig. 1C). In conclusion, RNA polymerase IV is essential for production of repeat associated sRNAs in different tissues of rice and genetic complementation of one of the two isoforms of NRPD1 rescues the sRNA levels.

Given the severity of the defects found in the rice *kd* plants, we hypothesised that the loss of sRNA mediated regulation of transposons and the concomitant mis-regulation of genes might be attributable to the defects. Since the 24nt sRNAs are involved in establishing DNA methylation of the repeats, we profiled the DNA methylome at the annotated repeats in rice which revealed a substantial decrease of methylation in *kd* when compared to WT especially in CHG and CHH contexts (Supplemental Fig. S5A). In addition, several of the transposon classes showed increase in expression (Supplemental Figs. S6A-C). The upregulation of repeats and loss of DNA methylation over transposons prompted us to question if they have gained proliferative potential upon pol IV *kd*. This possibility was previously predicted to influence phenotypes and genomic integrity in a number of reports (Debladis et al. 2020; Cui et al. 2013; Wei et al. 2014). To verify such a possibility, we performed a PCR-based assay to detect the extrachromosomal circular DNA (ECC DNA) intermediates that are produced as by-products of transposition (Lanciano et al. 2017). We observed additional bands portraying increased proliferative potential of *PopRice and Tos17* transposons (Supplemental Fig. S6D). These bands were observed even after the removal of linear DNA using a specific exonuclease (Supplemental Fig. S6E and methods). We examined if specific transposons proliferated by performing transposon display Southern hybridisations. Among the *kd* lines tested, specific line showed discernible copy number increase of LINE1 element across generations (Supplemental Fig. S6F). This observation is in line with a previous report where silencing of NRPD1 caused an increase in transposon copy numbers (Debladis et al. 2020). This lineage specific effects on transposon copy number is shown by a previous study as well (Debladis et al. 2020), and this might be the reason for exacerbated reproductive defects in selected *kd* lines (Supplemental Fig. S3A). Notably, ribosomal RNA (rRNA) precursor expression and the expression of several genes were also mis-regulated in the *kd* lines when compared to the WT (Supplemental Figs. S5B, C).

In summary, the activity of RNA polymerase IV and ensuing sRNAs are essential for regulating expression of genes and repeats via DNA methylation and loss of these processes result in potentially genotoxic proliferation of transposons in the rice genome.

### Activity of RNA pol IV potentiates silencing by H3K27me3 and H3K9me2 marks and modulates their relative occupancy over genes and repeats

Several reports have suggested that the effective silencing is brought about by the cross talk between DNA methylation and histone modifications in a locus specific manner in plants (Mathieu et al. 2005; Deleris et al. 2012; Gent et al. 2014; Zhou et al. 2016). These reports have found that loss of DNA methylation led to perturbation of post translational modifications (PTMs) of histones at the respective loci, and together influencing status of transcriptional silencing (Jamge et al. 2022). Recent studies in Arabidopsis have suggested that loss of pol IV and associated sRNAs can influence the histone H3 PTMs levels at specific loci (Li et al. 2020; Parent et al. 2021). In particular, Rougee et al. (Rougée et al. 2021) have described H3K27me3 redistribution at a subset of *ddm1* hypomethylated loci in Arabidopsis without being able to repress the hypomethylated loci. These observations in Arabidopsis, along with lack of in-depth epigenetic analysis in Arabidopsis *nrpd1* mutants, prompted us to explore the multiple layers of epigenetic changes upon *kd* of pol IV in rice.

In order to examine the polycomb mediated H3K27me3 profiles, we performed nuclear immunostaining that revealed broader H3K27me3 occupancy in *kd* unlike the WT and the CO lines corroborating observations in Arabidopsis DNA methylation mutants (Fig. 2A). Further, to examine the H3K27me3 marks and other epigenetic marks in a locus specific manner, we performed ChIP-seq with H3K27me3, H3K4me3 and H3K9me2 specific antibodies in replicates that showed consistent enrichment profiles (Supplemental Fig. S7). We analysed the differential enrichment of the H3K27me3 mark genome-wide (Methods). This revealed 7686 loci possessing significantly higher H3K27me3 occupancy in *kd* when compared to WT, validating the immunostaining observations. Surprisingly, we found that the loci that showed enhanced occupancy of H3K27me3 marks also showed loss of H3K9me2 marks (Fig. 2B). This observation points out that pol IV activity is quintessential not only for maintaining the DNA methylation and sRNA profiles, but also to buffer the relative locations of gene-specific H3K27me3 marks and repeat-specific H3K9me2 marks. In the absence of pol IV, H3K27me3 marks intruded into new domains. On the other hand, H3K9me2 marks were reduced over the loci intruded by H3K27me3 marks, thus redistributing the silencing signatures (Fig. 2B).

**Figure 2.**
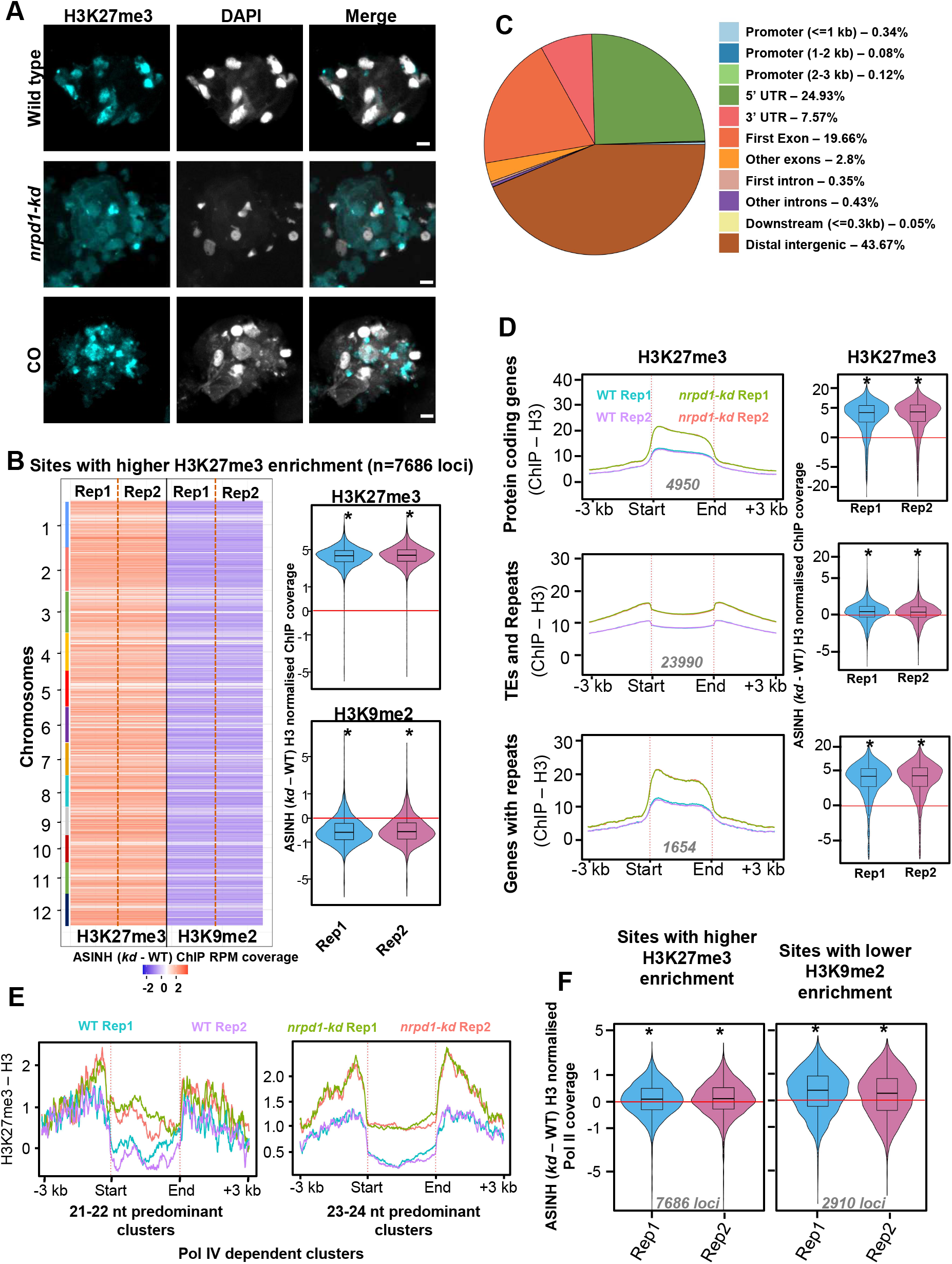
Pol IV maintains the genome-wide distribution of H3K27me3 mediated silencing states. *(A)* Immunostaining images of H3K27me3 marks in the nuclei extracted from WT, *kd* and CO. DAPI stained DNA. Scale bar: 5 µm. At least 40 nuclei from each genotype were examined. *(B)* Chromosome resolved heatmaps showing the levels of difference in H3K9me2 enrichment at sites with higher H3K27me3 enrichment in *kd* when compared to WT. Box-violin plots show the distribution of the differences in H3K9me2 and H3K27me3 levels. The Y-axis is scaled to inverse sine hyperbolic function of differences. *(C)* Pie-chart showing the feature annotation of the sites shown in *(B)*. All the sites that do not fall in the +/-3kb windows of genes were categorised as distal intergenic regions. *(D)* Metaplots showing the levels of H3K27me3 enrichment (normalised to H3) over the annotated features) overlapping with the identified H3K27me3 peaks. Numbers in grey depict the number of loci. Box-violin plots show the difference in enrichment in *kd* compared to WT. The Y-axis is scaled to inverse sine hyperbolic function of H3 ChIP normalised values. *(E)* Metaplots showing the abundance normalised enrichment of H3K27me3 marks w.r.t. H3 over the pol IV dependent sRNA clusters size categorised as 21-22nt and 23-24nt predominant clusters. *(F)* Box-violin plots showing the difference in Pol II coverage over the H3K27me3 higher enriched sites and H3k9me2 lower enriched sites in *kd* compared to the WT. Numbers in grey describe the number of loci taken for analyses. The Y-axis is scaled to inverse sine hyperbolic function of enrichment values. *(B, D* and *F)* Mann Whitney U test. * p-value < 0.0001.

Furthermore, annotation of the H3K27me3 occupied loci showed that almost 55% of them corresponded to coding genes with around 45% of the loci concentrated at the 5’-UTR (24.93%) and first exons (19.66%) of the protein coding genes (‘PCGs’) (Fig. 2C). All the non-PCGs accounted for 44% of the H3K27me3 over-enriched loci (Fig. 2C). Analysis of H3K27me3 peaks further revealed that it was globally over-enriched at PCGs, TEs and repeat loci, thus implicating the spread of this mark into both PCGs and non-PCGs in *kd* (Fig. 2D). Since monocot genomes are relatively enriched with repeats where they can pronouncedly influence the proximal genes to a greater extent than dicots (Hirsch and Springer 2017), we selected peaks containing PCGs that have at least 10% of their length overlapping with repeats (‘Genes with repeats’), and explored the redistribution of histone marks. As expected, this subset of PCGs recapitulated the hyper-occupancy of the H3K27me3 marks over the gene bodies to a greater degree than PCGs (Fig. 2D). Moreover, the hallmark RdDM sites enriched in pol IV dependent sRNAs gained H3K27me3 marks (Fig. 2E). This is in line with the observation that, even though the silencing by H3K9me2 and H3K27me3 are compartmentalized, they can, to a certain degree, substitute and overlap in cases of perturbation of other silencing signatures (Deleris et al. 2012; Déléris et al. 2021).

Surprisingly, in spite of the spreading of H3K27me3 silencing in *kd*, we observed enhanced transcription of transposons and repeats in *kd* lines, indicating that H3K27me3 overshoot and intrusion did not completely compensate for the loss of repressive signatures of H3K9me2, DNA methylation and sRNAs, supporting earlier investigations in Arabidopsis (Supplemental Figs. S4, 5 and 6) (Rougée et al. 2021). In order to test if occupancy of H3K27me3 marks can effectively prevent transcription at the loci that lost H3K9me2 marks, we performed ChIP-seq with RNA polymerase II antibody. This analysis revealed the over occupancy of pol II ChIP signals over the sites with increased H3K27me3 or reduced H3K9me2 occupancy (Fig. 2F) suggesting inefficient suppression of transcription by H3K27me3 as established in *ddm1* mutants in Arabidopsis (Rougée et al. 2021). Taken together, we identify a novel effect of loss of pol IV in buffering the occupancy of H3K9me2 and H3K27me3 marks over the PCGs and repeats. Failure to establish this balance led to a more permissive chromatin state, promoting mis-regulation of PCGs and repeats.

### Aberrant transcription in *kd* lines gives rise to atypical sRNAs that are suppressed by pol IV

Relaxed chromatin state and the reduction of silencing signatures in the *kd* lines might favour the enhanced transcription. Such observations were routinely reported in model dicot Arabidopsis and other organisms when mutants with altered chromatin state were analysed (McKinlay et al. 2018; Zheng et al. 2009; Ishihara et al. 2021; Jamge et al. 2022). In order to address this possibility, we performed pol II ChIP-seq using WT and *kd* panicles. We profiled the pol II status over PCGs and genes with repeats. Even though we did not observe significant difference of pol II occupancy over PCGs, we observed a significant hyper occupancy over the genes with repeats (Supplemental Figs. S8A, B). Since the antibody captures all the states of pol II, we postulated whether aberrant assembly, stalling or elongation of the pol II in *kd* might trigger the accumulation of mis-formed potentially aberrant transcripts. We explored if these transcripts over the subset of PCGs might trigger production of sRNAs from these regions. As anticipated, sRNA analysis showed that the numerous ShortStack identified sRNA clusters (‘clusters’) were upregulated in *kd* in all the three tissues tested in both the size classes (Supplemental Fig. S9A). The sRNAs that are upregulated in *kd* were denoted as pol IV suppressed sRNAs (‘suppressed sRNAs’) and those that are reduced in *kd* were named as pol IV dependent sRNAs (‘dependent sRNAs’). Surprisingly, suppressed sRNAs did not show any first nucleotide bias (Supplemental Fig. S9B). Further analysis revealed that these suppressed sRNA clusters had 21-22nt predominant, 23-24nt predominant and mixed clusters of both these size classes (Supplemental Fig. S10A; Methods). The suppressed and dependent clusters had uniform length distribution (Supplemental Fig. S10B).

In order to have an unbiased estimation of the relative distribution of these sRNAs, we performed a genome window (100bp) based analyses (Methods). While dependent bins were higher in number in all the three pol IV *kd* tissues, suppressed sRNA loci were fewer (Supplemental Fig. S11). We verified that these atypical sRNAs are not due to oversampling of the residual sRNAs in the *kd* tissues and associated normalisation artefacts by comparing the raw abundance of sRNAs in each library to the sum total of miRNAs in them (Supplemental Fig. S12). The suppressed bins were concentrated at fewer selective loci (Supplemental Figs. S13A, S14A and S15A). Further, cumulative sum plots describing the relation between the abundance of sRNAs and the number of uniformly sized bins also indicated that suppressed sRNAs were abundantly concentrated in fewer bins (Supplemental Figs. S13B, S14B and S15B).

Further to validate if suppressed sRNAs in *kd* are associated with other pol IV machineries, we analysed the published sRNA datasets derived from rice *nrpd1, rdr2* and *nrpe1* seedlings (Zheng et al. 2021; Wang et al. 2022). We indeed observed the accumulation of suppressed sRNAs in *nrpd1* and *rdr2*, but to a lesser degree in *nrpe1* seedlings (Fig. 3A). Considering how RDR2 is linked to pol IV transcription, this observation is not surprising. Further, we observed that the *nrp(d/e)2* sRNA datasets (Chakraborty et al. 2022) indeed retained the pol IV suppressed loci, suggesting only a minor role of pol V in altering the chromatin state and the accumulation of these sRNAs (Fig. 3A). We also detected the pol IV suppressed sRNAs by sRNA northern hybridisations (Fig. 3B), incontrovertibly proving the existence of the suppressed sRNAs in *kd* lines and they were not observed upon NRPD1 complementation (Fig. 3C). Furthermore, repeat features that are hallmarks of dependent sRNAs were completely devoid of suppressed sRNAs (Fig. 3D). Annotation of the differentially expressed sRNA bins from different tissues identified that nearly half of the pol IV suppressed sRNA bins mapped to coding regions while dependent counterparts mapped to the distal intergenic regions as expected (Supplemental Fig. S16A). In addition, the sRNAs from the suppressed bins overlapping with the PCGs mapped uniformly to both the sense and antisense strand of the gene, suggesting activity of one of the RDRs (Supplemental Fig. S16B). Unlike dependent sRNAs, suppressed sRNAs were within the PCGs (Supplemental Fig. S16C).

**Figure 3.**
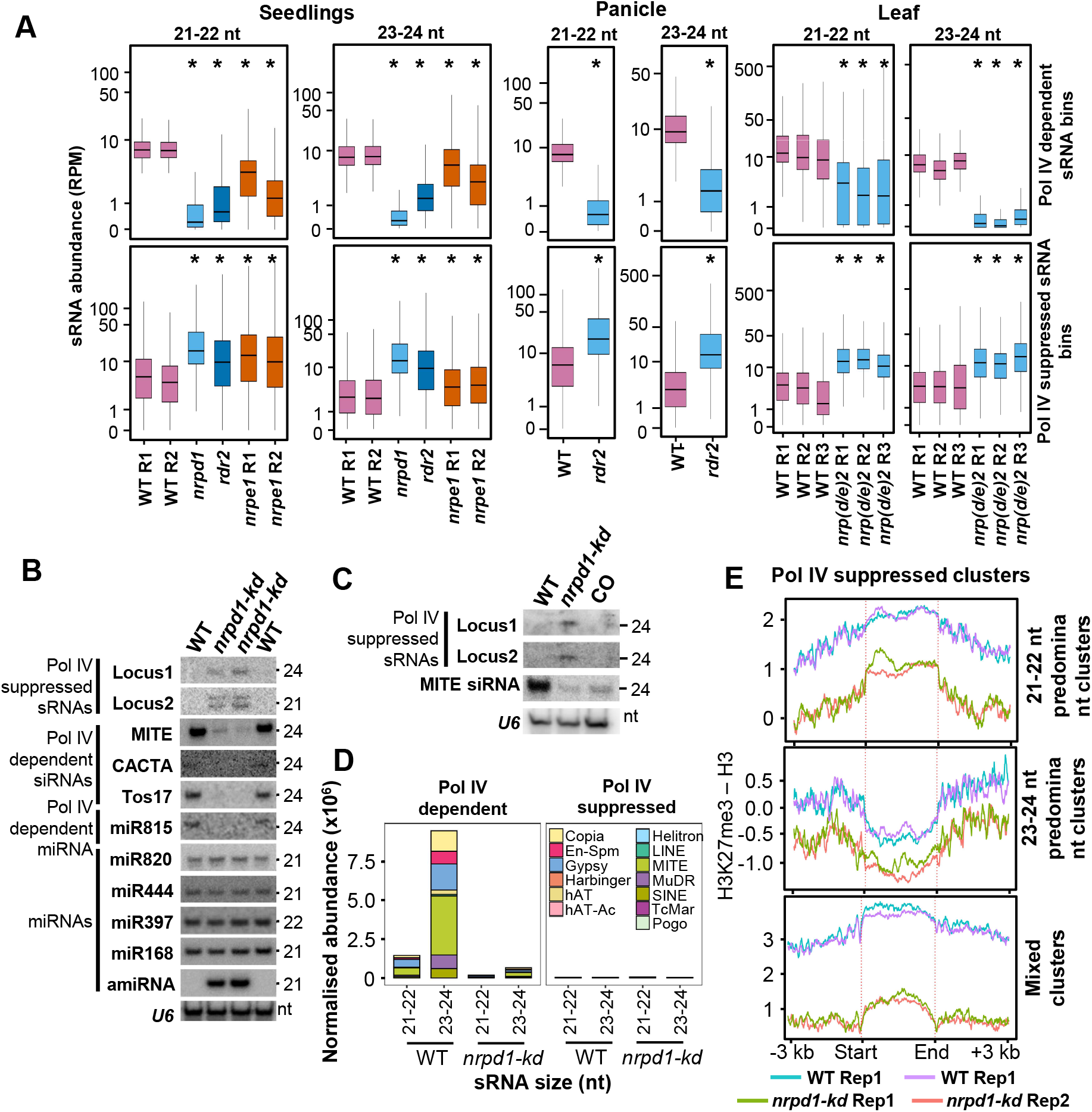
Pol IV complex suppresses sRNA production from several loci. *(A)* Boxplots showing the abundance of sRNAs from different mutants in rice. sRNAs were size-categorized into 21-22nt and 23-24nt and counted in 100bp non-overlapping windows. Plots depict the abundance of sRNAs in each size class over *nrp(d/e)2* (leaf)*, nrpd1* (seedlings) and *rdr2* (panicle) dependent and suppressed bins. The datasets are from GSE158709 (seedlings – *nrpd1, nrpe1*) and GSE130166 (seedlings and panicle – *rdr2*). The Y-axis is scaled to inverse sine hyperbolic function of RPM values. Mann Whitney U test. * p-value < 0.0001. *(B)* Small RNA northern blots validating the presence of pol IV suppressed sRNAs, dependent sRNAs and independent miRNAs. *U6* was used as loading control. *(C)* sRNA northern blots showing the abundance of sRNAs in WT, *kd* and NRPD1 complementation (CO) lines. *U6* was used as loading control. *(D)* Stacked bar plots showing abundance of Pol IV dependent and suppressed sRNAs from transposon categories. *(E)* Metaplots describing histone H3K27me3 occupancy normalised to total H3 signal over the pol IV suppressed sRNA clusters, categorized as 21-22 nt predominant clusters, 23-24 nt predominant clusters and the mixed sized clusters.

Furthermore, occupancy of H3K27me3 marks at the suppressed loci of all size classes were lower than WT in *kd* lines suggesting a chromatin mis-regulation associated with these loci (Fig. 3E). Moreover, genes overlapping the suppressed sRNA loci showed reduced H3K27me3 and H3K9me2 levels (Supplemental Fig. S17). In summary, we revealed a population of novel sRNAs that are suppressed by pol IV but not by pol V. These sRNAs are atypical in terms of their genomic origins and molecular signatures.

### Pol IV suppressed sRNAs are conserved in Arabidopsis and restricted to the mutants of pol IV arm of RdDM

We speculated that altered chromatin state due to loss of pol IV might be conserved across plants, and might be leading to the accumulation of suppressed sRNAs. In agreement with this, we observed that loss of Arabidopsis pol IV resulted in suppressed sRNAs, very similar to rice, but from fewer bins in inflorescence tissues (Zhou et al. 2018), (Supplemental Fig. S18A). Unlike pol IV dependent bins, suppressed sRNAs in Arabidopsis showed non-uniform distribution similar to rice tissues (Supplemental Fig. S18B). Availability of various genetic mutants in Arabidopsis prompted us to check the mechanistic origins of these sRNAs and possible regulators of chromatin state when pol IV is not functional. In this direction, we examined several siRNA biogenesis mutants, especially connected to RdDM, and found that the suppressed sRNAs were found mainly upon loss of pol IV itself or associated proteins using northern blots with various mutants (Fig. 4A). The suppressed sRNAs were unique to *nrpd1* and were not observed in sRNA processing mutants such as *dcl3, dcl234, ago4* or a mutant of chromatin remodeller *ddm1* (Fig. 4A). One of the pol IV suppressed loci accumulated aberrant transcripts of ∼200nt detected with specific probes designed to bind to suppressed sRNAs supporting the notion that these were indeed products of aberrant transcription (Fig. 4B). We also used sRNA datasets from the similar stage inflorescence tissues from additional set of mutants involved in RNA silencing (Zhai et al. 2015). This independent set of *nrpd1* datasets also confirmed the increase of suppressed sRNAs by at least 10-fold compared to WT mainly from non-repeat regions (Supplemental Figs. S19A-C). Interestingly, the suppressed sRNAs were also seen in other *nrpd1* associated mutants like *rdr2, nrp(d/e)2, nrpd1dcl3* and *rdr2dcl3* (Supplemental Fig. S19C). This indicates that the suppressed sRNAs are directly coupled to the absence of pol IV complex (NRPD1 or RDR2). In addition, most of the suppressed sRNAs were brought back to the WT levels upon NRPD1 complementation (Supplemental Fig. S19D).

**Figure 4.**
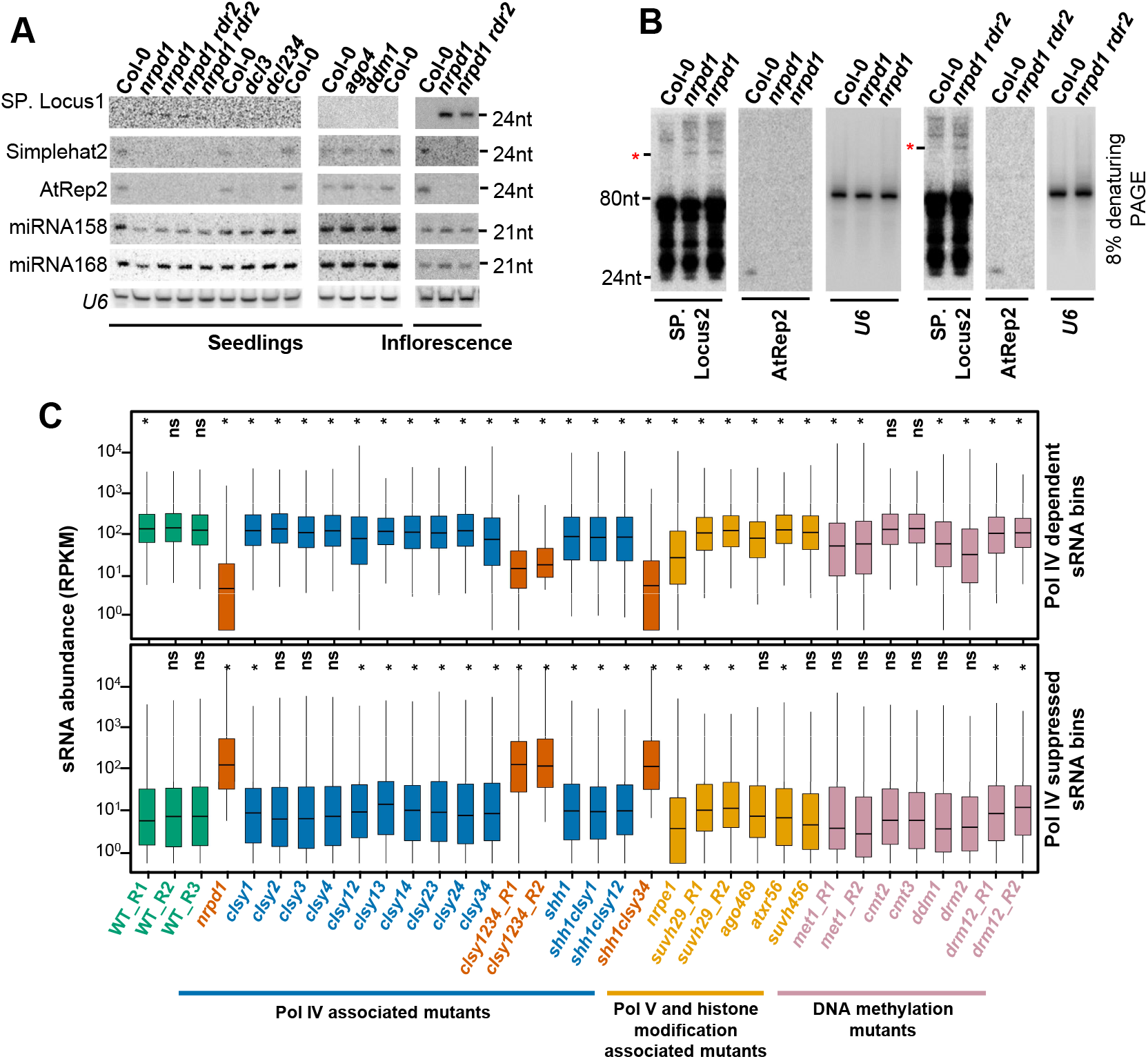
Polymerase IV suppressed sRNAs are conserved in Arabidopsis. *(A)* Small RNA northern blots showing the levels of pol IV suppressed sRNAs (SP. locus) and other sRNAs from Arabidopsis seedlings and inflorescence tissues. *(B)* Northern blots made with 8% denaturing PAGE gel depicting the resolved higher sized RNAs (*) from SP. locus. *(C)* Boxplots showing the sRNA abundance from different genotypes of Arabidopsis with counts from pol IV dependent and suppressed 23-24nt size class bins. The sRNA datasets were obtained from GSE165574, GSE45368 and GSE99694. Analyses depicts sRNAs from inflorescence tissues. The Y-axis is scaled to inverse sine hyperbolic function of RPKM values. Mann Whitney U test. * p-value < 0.0001.

In order to probe which part of the RdDM machinery is mainly involved in generation of pol IV suppressed sRNAs, we analysed published sRNA datasets from inflorescence tissues of 28 different genotypes of Arabidopsis mutants, pertaining to 3 categories – pol IV associated mutants, pol V and histone modification associated mutants and DNA methylation mutants (Supplemental Table S2). For the set of loci identified as suppressed and dependent loci, we evaluated the abundance of sRNAs across the mutants. We observed evident and drastic accumulation of suppressed sRNAs comparable to *nrpd1* only in *clsy1234* quadruple mutants and in *shh1clsy34* triple mutant (Fig. 4C). This interesting observation points out that the suppressed sRNAs are the effect of ablation of pol IV enzyme complex along with the machinery responsible for its assembly. On the contrary, the pol IV suppressed sRNAs are not initiated upon removal of either the pol V arm of RdDM or the DNA methyltransferases.

Probing the contributing molecular signatures associated with the suppressed sRNAs, we also tested the status of the DNA methylation in reproductive tissues over these clusters and found that methylation over CG, CHG or CHH contexts in both rice and Arabidopsis remain largely unchanged (Supplemental Fig. S20 and S21). Pol II occupancy over the suppressed and dependent loci also remained unchanged, likely due to the transient interaction by pol II in generating these transcripts (Supplemental Fig. S22). However, it is likely that aberrant RNAs generated by pol II were quickly cleaved into the dependant and suppressed sRNAs that we have observed. Taken together, these results indicate that loss of pol IV changes the chromatin globally and the latent H3K27me3 occupancy partially prevents aberrant transcription at pol IV dependent loci (Supplemental Figs. S23 and S24). On the other hand, reduction of H3K27me3 over specific genomic regions upon loss of pol IV might have triggered production of suppressed sRNAs, that are conserved across monocots and dicots.

### Pol IV suppressed sRNAs are loaded into AGO1 to induce PTGS at protein coding loci

Given that these abundant atypical sRNAs are from gene coding regions and they did not change the DNA methylation signature of the loci, we explored if they can target genes post transcriptionally. To test if AGO1-loaded suppressed sRNAs targeted genes post-transcriptionally, we performed degradome sequencing in WT and *kd* panicles. Among the genes that overlapped with the suppressed sRNAs (389 genes), target prediction tools identified 154 genes as potential targets of suppressed sRNAs (Additional File 6). The degradome tag density at the predicted target loci was substantially increased in *kd* lines (Fig. 5A). In further agreement with the targeting process, the AGO1 IP was enriched with suppressed sRNAs at the degradome predicted targeting position (Fig. 5B) (Wu et al. 2009). These target RNAs had significantly reduced expression, suggesting precise slicing at these sites mediated by suppressed sRNAs (Fig. 5C). We also observed that many of the genes that underwent PTGS by suppressed sRNA-mediated targeting in rice were previously implicated in reproductive growth and development (Additional File 6). For instance, OsMADS18 (APETALA1 homolog in rice), a member involved in floral architecture establishment (Wang et al. 2020a), a close homolog of fertility restorer (RF) (Os10t0497366), glycine rich interaction partner of RF5 (Os12t0632000) (Hu et al. 2012) and pollen specific desiccation associated protein (Os11t0167800) were targeted for degradation in *kd* lines (Additional File 6). In addition, these genes had reduced mRNA expression in *kd* lines. For example, RF5 interaction protein (Os12t0632000), mutant of which shows pollen non-viability due to loss of cytoplasmic male sterility (Hu et al. 2012) was not only effectively targeted in *kd* lines, but also had reduced mRNA expression in these lines (Additional File 6).

**Figure 5.**
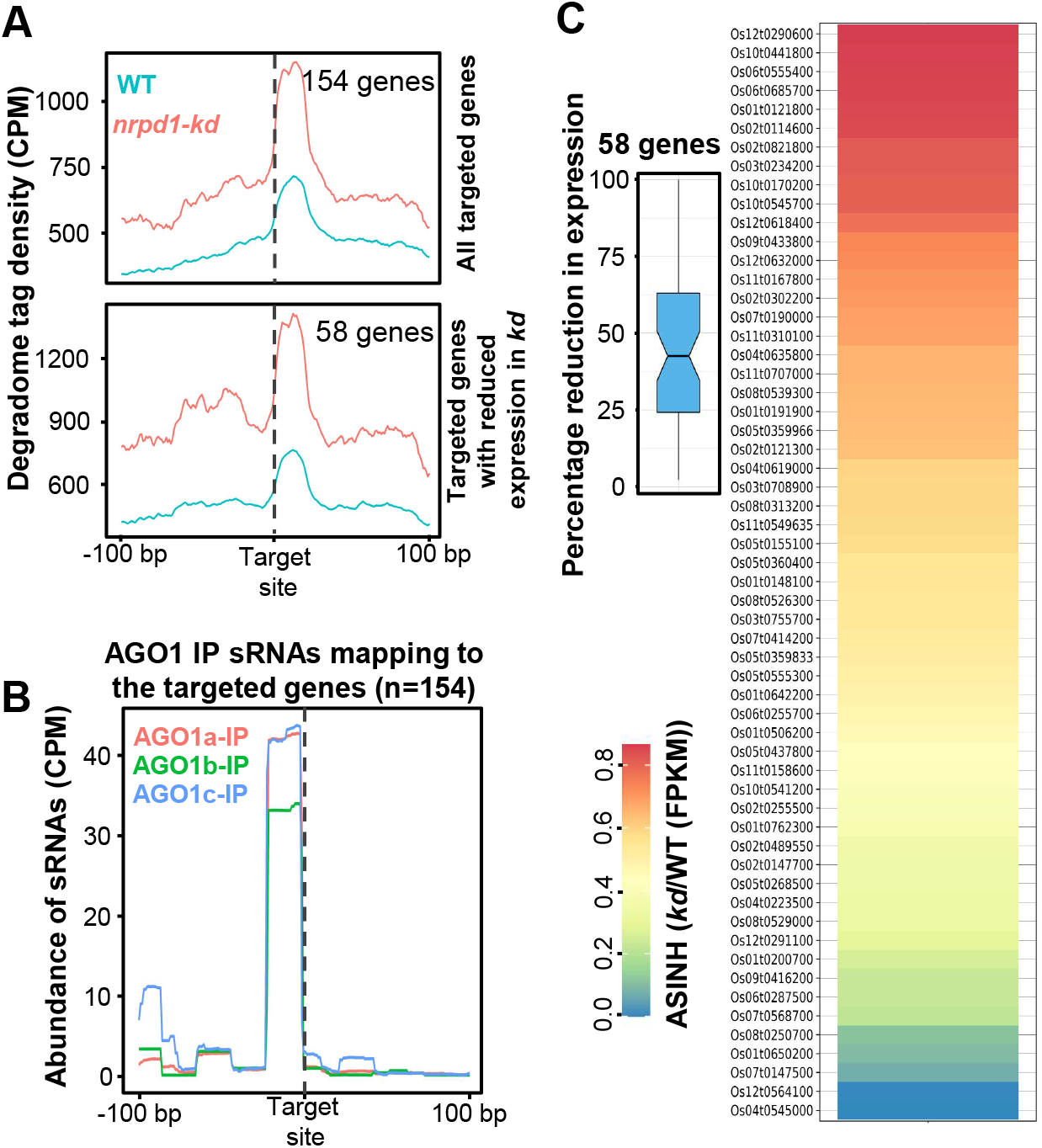
Pol IV suppressed sRNAs get loaded into AGO1 to mediate PTGS. *(A)* Metaplots showing the degradome tag density over the genes identified as targets in *nrpd1-kd* panicle degradome (top panel) and the same for the subset of targeted genes that show reduced expression in *kd* compared to WT. Targeted locations are centred and 100bp on either side is displayed. *(B)* Metaplots showing the abundance of AGO1 IP enriched sRNAs over the identified targeted genes centred at the targeting site. *(C)* Heatmap showing the fold changes (scaled to inverse sine hyperbolic function) in expression (FPKM) of the 58 genes that showed reduction in expression when compared to WT. The inset boxplot shows the distribution of reduction in expression (FPKM) observed.

On the other hand, the suppressed sRNAs in Arabidopsis did not accumulate in the AGO1 IP datasets in both Col-0 and *nrpd1* plants while the dependent sRNAs were abundantly found in AGO4 IP datasets (Panda et al. 2020; Zhai et al. 2015) (Supplemental Fig. S25A). While such targeting is not possible in Arabidopsis where suppressed sRNAs are neither abundant nor get loaded into AGO1 (Supplemental Fig. S25B), it is tempting to speculate that abundant sRNAs target same or similar key genes in other plant species where perturbation of NRPD1 showed strong phenotypes (Wang et al. 2020b; Grover et al. 2018). Such a targeting mediated by suppressed sRNAs might act as strong deterrent for the loss or reduction of pol IV activity, and might contribute towards evolution of additional copies of this polymerase. In fact, monocots that have large genomes, also code for pol VI that is well expressed and appears to have neo-functionalized to partly carry out pol IV activity (Trujillo et al. 2018).

## Discussion

RNA polymerases play a crucial role in the central dogma of life by transcribing the genetic material into functional RNA entities of different types. Activity of these polymerases are under tight control of several layers – epigenetic aspects of the substrate chromatin as well as the availability of necessary cofactors like transcription factors, enhancers and other chaperones. Two major conserved contributors for epigenetic silencing of this nature are DNA methylation and histone modifications that relay the regulatory outcome. These marks mainly prevent the spurious transcription by RNA polymerases at unintended locations. The precise and dynamic regulation of these modifications in a coordinated manner hallmarks the centre-stage of regulation. A mediator of such a synergistic response modulating both of these aspects is paramount. Such a mediator should not only obstruct illegit transcription but also prevent unwarranted occlusion of transcription of necessary genes and loci. Our investigation uncovers a plant specific RNA polymerase, pol IV, to be endowed with such a potential.

Evolutionary analyses suggest that plant-specific RNA polymerases IV, V, and in specific cases VI are novel machineries evolved in conjunction with the complexity of plant genomes and the degrees of importance of these polymerases varies across plants. For instance, in early land plants like *Physcomitrella patens* the pathway is completely redundant with other sRNA-independent DNA methyltransferases introducing *de novo* DNA methylation (Yaari et al. 2019) as opposed to the widespread defects observed in other plants with severity in rice (Fig. 1 and Supplemental Figs. S1, S2 and S3) (Zheng et al. 2021). The functions of these polymerases are pronounced in reproductive tissues as they are involved in faithful transmission of epigenetic information, antagonizing genome dosage aberrations and hybridisation (Satyaki and Gehring 2019; Erdmann et al. 2017; Martinez et al. 2018; Zhang et al. 2016). It is apparent that the green-lineage specific RNA polymerases have neo-functionalized to perform additional roles with increasing genome complexity. The comparisons we present with respect to redistribution of epigenetic marks between rice and Arabidopsis serves as strong evidence to these predictions.

Several investigations in the model plant Arabidopsis have established that pol IV initiates biogenesis of sRNAs from repeats and transposons establishing *de novo* DNA methylation aiding against their genotoxic proliferation. Our studies in rice corroborate this function of pol IV in multiple reproductive tissues (Fig. 1 and Supplemental Fig. S4). Several reports have mentioned the synergistic effects and cross-dependence of histone modifications and global CG DNA methylation in establishing epigenetic modalities in Arabidopsis (Soppe et al. 2002; Mathieu et al. 2005; Deleris et al. 2012; Li et al. 2018; Zhong et al. 2021). These studies had employed *met1* and *ddm1* mediated DNA methylation loss occurring at the CG contexts globally resulting in loss of compaction at the pericentromeres and constitutive heterochromatin domains. On the other hand, the RdDM is not restricted to heterochromatin and its purview extends into the gene rich regions as well. Whether modulation of the sRNAs can directly perturb the chromatin states attributable to the gene expression was unclear. Especially, in monocots like rice, the interspersion of repeat fragments within genes mandates gene regulation not at extended length scales but in localised compartments even within the euchromatin (Espinas et al. 2020). Probing the effect of loss of RdDM on the chromatin states over the genes in our study uncovered a distinct redistribution of H3K9me2 and H3K27me3 marks (Fig. 2). Predominantly, H3K9me2 marks, attributed with well-suppressed constitutively silenced loci and H3K27me3 associated with silenced gene coding units by action of Polycomb Represive Complex2 showed over-occupancy in *kd* over the transposons and genes. Strikingly, the authentic pol IV transcribing loci that result in sRNAs showed significant H3K27me3 occupancy, likely an ectopic compensatory mode of silencing (Fig. 2E). This feature is evolutionarily conserved between plants and animals and loss of transposon methylation can contribute to the redistribution of these marks into transposon territories (Deleris et al. 2012; Déléris et al. 2021). Close proximity of the protein coding genes and transposons might be the trigger causing the intrusion of facultative silencing marks on the genes in species with larger genomes such as rice. This is well-supported by the fact that upon *kd* of the pol IV, genes that are dispersed with the repeat fragments showed increased degree of H3K27me3 intrusion (Fig. 2E). PRC2 mediated H3K27me3 on genes impacts reproductive success in plants by controlling several imprinted genes that scale the genome dosage (Köhler et al. 2003; Roszak and Köhler 2011; D. et al. 2013; Jiang et al. 2017). Such indirect effects on the protein coding genes by unwarranted H3K27me3 marks, to a certain extent, might explain the defects in *kd*.

Loss of silencing signals and redistribution of H3K27me3 marks over the protein coding genes upon loss of RdDM in rice should result in differential accessibility of these loci to RNA polymerases. Ungauged transcription in the absence of pol IV might feed to the small RNA pools via activity of DICER-LIKE proteins that are devoid of pol IV precursor load. Exploration of sRNA pools in the *nrpd1* mutants of rice and Arabidopsis indeed showed resultant aberrant sRNAs, likely triggered by spurious transcripts. Interestingly, such spuriously transcribed loci showed reduced in H3K27me3 occupancy (Fig. 3D). This commonality of spurious sRNA transcripts in RdDM mutants is observed in *nrpe1* very recently (Zheng et al. 2021) where a similar notion of chromatin relaxation triggering pol IV and pol II transcription is promulgated. Similarly, studies in maize and Arabidopsis have suggested atypical transcripts in the *nrpd1* mutation (Erhard et al. 2015; McKinlay et al. 2018). Our studies mechanistically delineate the causative chromatin features in the *nrpd1* mutant. Not limiting to that, we find that the resultant pol IV suppressed sRNAs are capable of targeting genes post transcriptionally by loading into specific AGOs.

These analyses reveal that the atypical pol IV suppressed sRNAs from diverse plants such as rice and Arabidopsis, were a result of mis-regulated chromatin states due to the perturbations to pol IV, and at least in rice, these are capable of targeting protein-coding genes post transcriptionally. Such atypical sRNAs are limited to selective loci as the poised loci for aberrant transcription from repeats are protected by compensatory H3K27me3 silencing. We catalogue reciprocity of the silencing between transcriptional and post-transcriptional modes, repurposing an existing machinery. Arabidopsis *nrpd1* related mutants also exhibited production of these sRNAs, nevertheless, they did not get loaded into AGO1 that potentiates them for active targeting. Absence of efficient PTGS in Arabidopsis *nrpd1* mutants might have alleviated the extent of defects unlike in rice. These results suggest a strong and multifaceted impact of pol IV in the expression of protein coding genes, especially in rice while encouraging evolution of additional genomic complexities, architecture and heterogeneity.

## Materials and Methods

### Plant material

*Indica* variety rice (*Oryza sativa indica sp.*) Pusa Basmati 1 (PB1) plants were grown in growth chamber at 24°C/70% RH/16h-8h light-dark cycle for hardening before transferring to greenhouse maintained at 28°C with natural day-night cycle. Arabidopsis plants were grown in controlled growth chamber maintained at 22°C/70% RH/16h-8h light-dark cycle. Different mutants used were reported in Supplemental Table S4.

### Binary vector construction and *Agrobacterium* mobilisation

For amiR precursor binary construct, the artificial miRNA was designed using WMD3 tool (Ossowski et al. 2008). The amiR was chosen so that it targets both the NRPD1 isoforms (NCBI Ids: NRPD1a – XM_015781553.2 and NRPD1b – XM_015756207.1) and optimal miRNA parameters were satisfied (Narjala et al. 2020). The mature amiR sequence (5’-UAUAGUGUUACUCUUGGACAU-3’) was embedded in the *Osa*miR528 precursor in the pNW55 plasmid using WMD3 recommended primers (Warthmann et al. 2008; Ossowski et al. 2008) (Supplemental Table S3). The precursor region was sub-cloned into a binary vector (pCAMBIA1300 backbone) between *Zm*Ubiquitin1 promoter and 35S poly(A) signal and mobilised into *Agrobacterium* cells using electroporation method.

For the complementation construct, NRPD1b promoter and 5’-UTR were amplified from the genomic DNA (chr09:22015503-22019281) and fused to the amiR-resistant CDS of OsNRPD1b (obtained by site directed mutagenesis) and this was cloned into pCAMBIA3300 vector (with bialaphos resistance marker, BlpR). The construct was super-transformed to the HygR *kd* calli using the same method described.

### Southern hybridisation

Southern hybridisation was performed as described previously (Ramanathan and Veluthambi 1995; G and Shivaprasad 2022). Total DNA (around 5 µg) was isolated from plant tissues using CTAB method and digested using 30 units of appropriate restriction enzyme overnight. The digested DNA was electrophoresed on a 0.8% agarose gel in 1x TBE buffer and capillary transferred to Zeta probe nylon membrane (Biorad). The transferred membrane was UV crosslinked and taken for hybridisation. The probes (PCR amplified from gDNA or binary plasmid using the oligos in Supplemental Table S3) were internally labelled using [α-P^32^] dCTP (BRIT India) using Rediprime labelling kit (GE healthcare) and hybridisation and washes were performed as described. The blots were exposed to phosphor imaging screen and scanned using Typhoon scanner (GE healthcare).

### sRNA northern hybridisation

sRNA northern blots were performed as described earlier (Tirumalai et al. 2020; Shivaprasad et al. 2012). Around 15 µg of TRIzol extracted total RNA from different tissues was electrophoresed on a denaturing 15% acrylamide gel. The gel was electroblotted onto Hybond N+ membrane (GE healthcare) and UV crosslinked. The membrane was hybridised with the T4 PNK end labelled oligonucleotides (with [γ-P32]-ATP) in hybridisation buffer (Ultrahyb buffer – Invitrogen). The blots were washed, exposed to phosphor imaging screen and scanned using Typhoon scanner (GE healthcare). Post-scanning, blots were stripped at 80°C in stripping buffer and proceeded with repeat hybridisations with subsequent probes.

### Chromatin Immunoprecipitation – sequencing (ChIP-seq) and analyses

ChIP was performed as described by earlier methods (Song et al. 2016; Saleh et al. 2008). Around 1.2 grams of pre-emerged panicle tissues were taken and crosslinked with 1% formaldehyde. The crosslinked tissues were pulverized in liquid nitrogen and nuclei were isolated. Equal number of nuclei were lysed and sheared using ultrasonication (Covaris) until fragments substantially accumulate in the 150-350bp size range. The sheared chromatin was incubated overnight with 50ul of protein G dynabeads (Thermo Fisher) at 4 degrees that were pre-bound to the appropriate antibodies. The beads were pre-bound with 4-5 micrograms of antibodies (H3K4me3 – Merck 07-473; H3K9me2 Abcam ab1220; H3K27me3 – Acive Motif 39155; H3 – Merck 07-10254; PolII – Abcam ab817) before incubation with the sheared chromatin. Washes, elution, decrosslinking and purification were performed as described. The purified IP products were taken for library preparation using NEBNext® Ultra™ II DNA Library Prep with Sample Purification Beads (NEB, E7103L) as per manufacturer’s protocol. The libraries (with replicates) were sequenced on Illumina HiSeq 2500 platform in single end mode (50bp).

The obtained datasets were adapter trimmed using cutadapt (Martin 2011) and aligned to the IRGSP 1.0 genome using Bowtie2 tool (Langmead and Salzberg 2012) with the following parameters: -v 1 -k 1 -y -a --best –strata. PCR duplicates were removed before further analyses. The alignment files were converted to coverage files and compared (difference) to total H3 signal (for histone H3 PTMs) using bamcompare utility of Deeptools (Ramírez et al. 2014). The average signals over the desired regions/ annotations were estimated using computematrix utility of deeptools and the coverage signal metaplots were plotted using plotprofile utility that were modified using custom R script. The peak calling was performed using MACS2 (Zhang et al. 2008) with broad peak calling for all ChIP datasets except for H3K4me3. The peaks with enrichment above 3-fold compared to H3 ChIP were taken as valid peaks and the peaksets were merged using bedtools across genotypes and replicates. The composite peak sets obtained were intersected with the annotated PCGs and repeats to get the peak overlapping features used for signal counting using Deeptools. The replicate concordance was measured across the samples using Deeptools plot correlation feature for the signals obtained over the composite peak sets.

### sRNA sequencing and analyses

Small RNA sequencing was performed from pre-emerged panicle, anther and endosperm of equally developed WT and *kd* plants. The size fractionation and library preparation was done as described in (Tirumalai et al. 2019). Reads obtained were quality checked, adapter removed and size selected using UEA small RNA workbench (Stocks et al. 2018). The reads obtained are aligned after categorisation into 21-22nt or 23-24nt sizes to IRGSP1.0 genome using Bowtie (Langmead and Salzberg 2012) with the following parameters: -v 1 -m 100 -y -a --best –strata. Step-wise analyses for genomic annotations and 5’-nucleotide abundance was performed as described earlier (Swetha et al. 2018). Only the mapped reads were used for further analyses. MicroRNA (miRNA) abundance was calculated using miRProf tool (Stocks et al. 2018). Arabidopsis datasets were handled the same way except that the reads were aligned to TAIR10 genome.

### Availability of data

Data generated or analysed during this study are included in this article and its supplementary files. Deep sequencing datasets have been deposited in GEO (https://www.ncbi.nlm.nih.gov/geo/) under the super-series GSE180457 (Refer Supplemental Table S1).

#### Reviewer TOKEN

To review GEO accession GSE180457:

Go to https://www.ncbi.nlm.nih.gov/geo/query/acc.cgi?acc=GSE180457

Enter token klynguiytngnfyd into the box

## Acknowledgements

We thank Professor K. Veluthambi for *Agrobacterium* strains, PB1 seeds and binary plasmids. We are grateful to Prof. David Baulcombe for the Arabidopsis mutants. We thank genomics, electron microscopy, IT, radiation, greenhouse and lab-kitchen facilities at the NCBS campus. We acknowledge the guidance of Anushree Narjala in bioinformatics analysis. We thank Dr. Dimple Notani for sharing reagents. We thank Rahul Raj Singh, M. Rajagopalan and Sumvit Goyal for amiR-NRPD1 binary plasmid cloning. We acknowledge the R based meta-plotting script from the Github page of Jeffrey Grover (https://github.com/groverj3). We are grateful to Nitish Dua and Mohammad Shariq for the help with immunostaining. We thank all the lab members for discussions and comments.

## Author contributions

PVS and VHS designed all experiments and discussed results and wrote the manuscript. VHS performed most of the experiments and bioinformatics analyses. SC performed bioinformatics analysis. DB generated transgenic lines. KP performed electron microscopy and micro-CT scanning. SR performed confocal imaging. TC generated and shared the *nrp(d/e)2* sRNA datasets. RAM helped with the analysis and discussion. All authors have read and approved the manuscript.

