## Supplementary Figures, Methods and Tables S1-4 for "Plant Polymerase IV sensitizes chromatin through histone modifications to preclude spread of silencing into protein-coding domains"

**Supplemental Figures S1-25**

Supplemental Figure S1. Knockdown of RNA pol IV in rice results in pleiotropic phenotypes.

Supplemental Figure S2. Distinct sets of Agrobacterium mediated transformation events yielded knockdown lines with similar phenotypic defects.

Supplemental Figure S3. Specific lineage of Pol IV knockdown plants showed severe defects that are not due to transgenesis or knockdown dosage suggesting possible epimutation.

Supplemental Figure S4. Loss of pol IV causes specific loss of repeats and transposon associated sRNAs.

Supplemental Figure S5. RNA pol IV is responsible for maintaining DNA methylation, repeat silencing and contributes to gene regulation.

Supplemental Figure S6. Pol IV silences transposons and repeats.

Supplemental Figure S7. ChIP – Sequencing exhibits concordant signals across replicates.

Supplemental Figure S8. Loss of pol IV impacts occupancy of pol II.

Supplemental Figure S9. Pol IV complex suppresses peculiar sRNA production from several loci.

Supplemental Figure S10. Pol IV dependent and pol IV suppressed sRNA clusters are of similar length.

Supplemental Figure S11. Pol IV dependent and pol IV suppressed sRNA bins are distinct in producing sRNAs with specific sizes.

Supplemental Figure S12. Pol IV dependent and pol IV suppressed sRNA bins are not due to oversampling and library normalisation.

Supplemental Figure S13. Pol IV suppressed sRNAs from panicle are non-uniformly distributed across the genome.

Supplemental Figure S14. Pol IV suppressed sRNAs from anther are non-uniformly distributed across the genome.

Supplemental Figure S15. Pol IV suppressed sRNAs from endosperm are non-uniformly distributed across the genome.

Supplemental Figure S16. Pol IV suppressed sRNAs have significant overlap with coding regions.

Supplemental Figure S17. Genes overlapping with suppressed sRNAs exhibit reduced silencing compensation by H3K27me3.

Supplemental Figure S18. Pol IV suppressed sRNAs from Arabidopsis inflorescence are non-uniformly distributed across the genome similar to rice tissues.

Supplemental Figure S19. Pol IV suppressed sRNAs from Arabidopsis are characteristically similar to that of rice.

Supplemental Figure S20. Canonical RdDM does not potentiate pol IV suppressed sRNAs in rice.

Supplemental Figure S21. Canonical RdDM does not potentiate pol IV suppressed sRNAs in Arabidopsis.

Supplemental Figure S22. Net Pol II occupancy over pol IV suppressed sRNA clusters does not change.

Supplemental Figure S23. Representative screenshots of epigenetic features at representative suppressed and dependent loci.

Supplemental Figure S24. Distinct H3K27me3 modifications at pol IV dependent and suppressed sRNA clusters.

Supplemental Figure S25.  Arabidopsis pol IV suppressed sRNAs are not effectively loaded into AGO1 and unlikely to target genes.

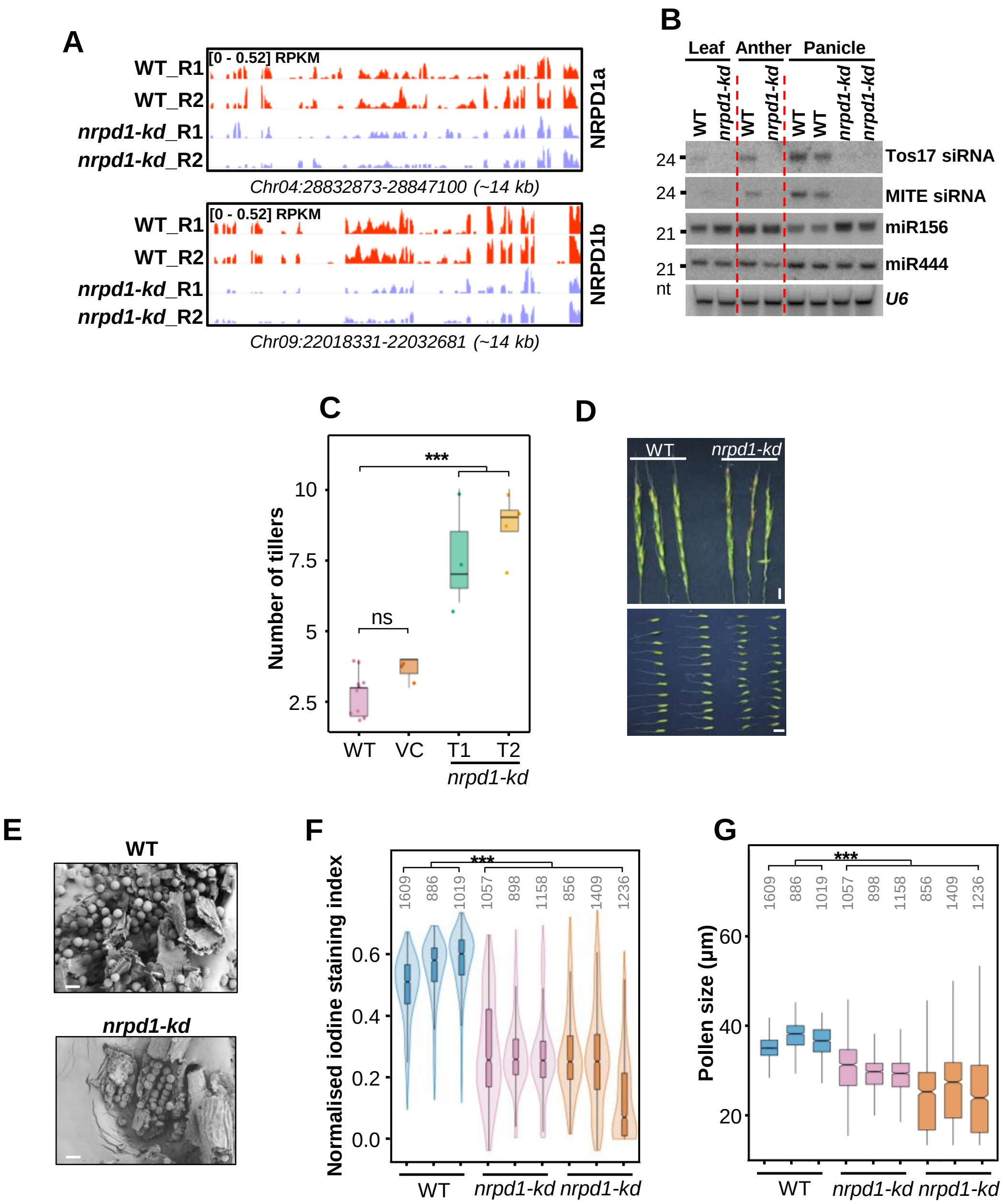


**Supplemental Figure S1.** Knockdown of RNA pol IV in rice results in pleiotropic phenotypes. *(A)* IGV screenshots of panicle RNA-seq coverage over the NRPD1a and NRPD1b loci in WT and *kd. (B)* sRNA northern blots indicating loss of 24nt siRNA in different tissues. miR444 and miR156 were examples of miRNAs. *U6* was used as loading control. *(C)* Boxplots of number of tillers observed in WT, vector control (VC), T1 and T2 generations of *nrpd1-kd* plants. Dots represent result of each plant (Tukey's test; ***p-value of 0.001,: *p-value of 0.01, ns-non-significant). *(D)* Images showing the panicle (top) and the individual florets (bottom) of WT and *nrpd1-kd* plants. Scale bar:1 cm. *(E)* Representative SEM images of pollen grains from dehisced anther. Scale: 40 μm. *(F)* Violin-box plots showing the distribution of normalised iodine staining index of pollen grains. Numbers in grey represent the number of pollen grains examined. (Wilcoxon test; *** p-value < $2.2x10^{-16}$). *(G)* Distribution of pollen sizes in WT and *nrpd1-kd* plants. The stained light microscopy images are quantified using custom ImageJ scripts to detect the sizes and degree of staining of pollen grains. (Wilcoxon test; *** p-value < $2.2x10^{-16}$).

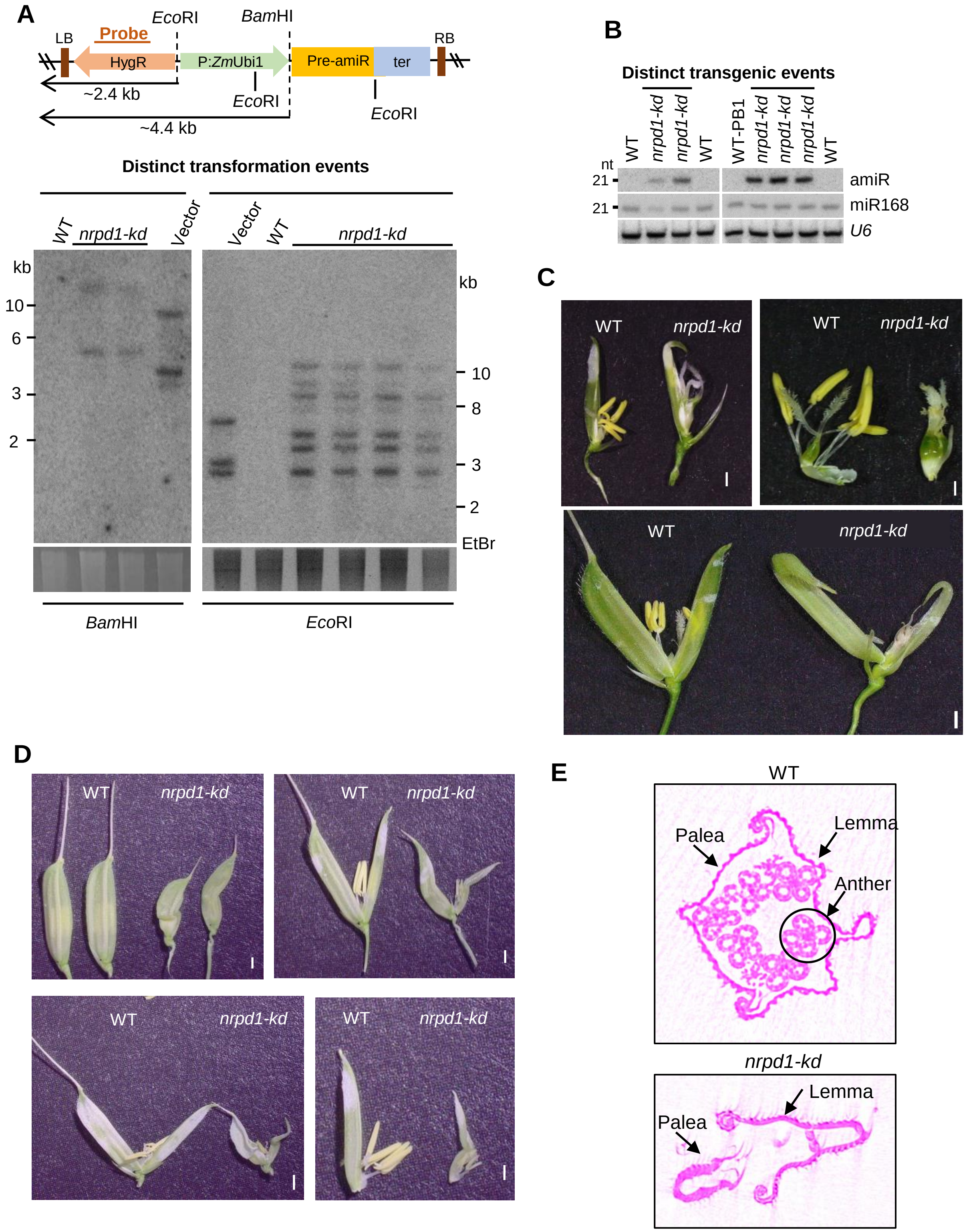


**Supplemental Figure S2.** Distinct sets of Agrobacterium mediated transformation events yielded knockdown lines with similar phenotypic defects. *(A)* DNA blots showing the T-DNA junction fragments corresponding to digestion of DNA with mentioned restriction sites. Minimum lengths of junction fragment sizes are mentioned by arrows on the T-DNA map above. Probe (HygR) region is mentioned in the map. *(B)* sRNA northern blots indicating the accumulation of amiRNA in transgenic knockdown (*nrpd1-kd*) plants obtained from independent transgenic events. miRNA168 is used as miRNA control and *U6* as loading control. *(C – D)* Defects in the reproductive structures obtained in the plants from two independent transgenic events as seen by distinct T-DNA integration pattern in *(A)*. Scale bars correspond to 1 mm. *(E)* Micro-computed tomographs (micro-CT) of equally developed florets in WT and *kd* lines.

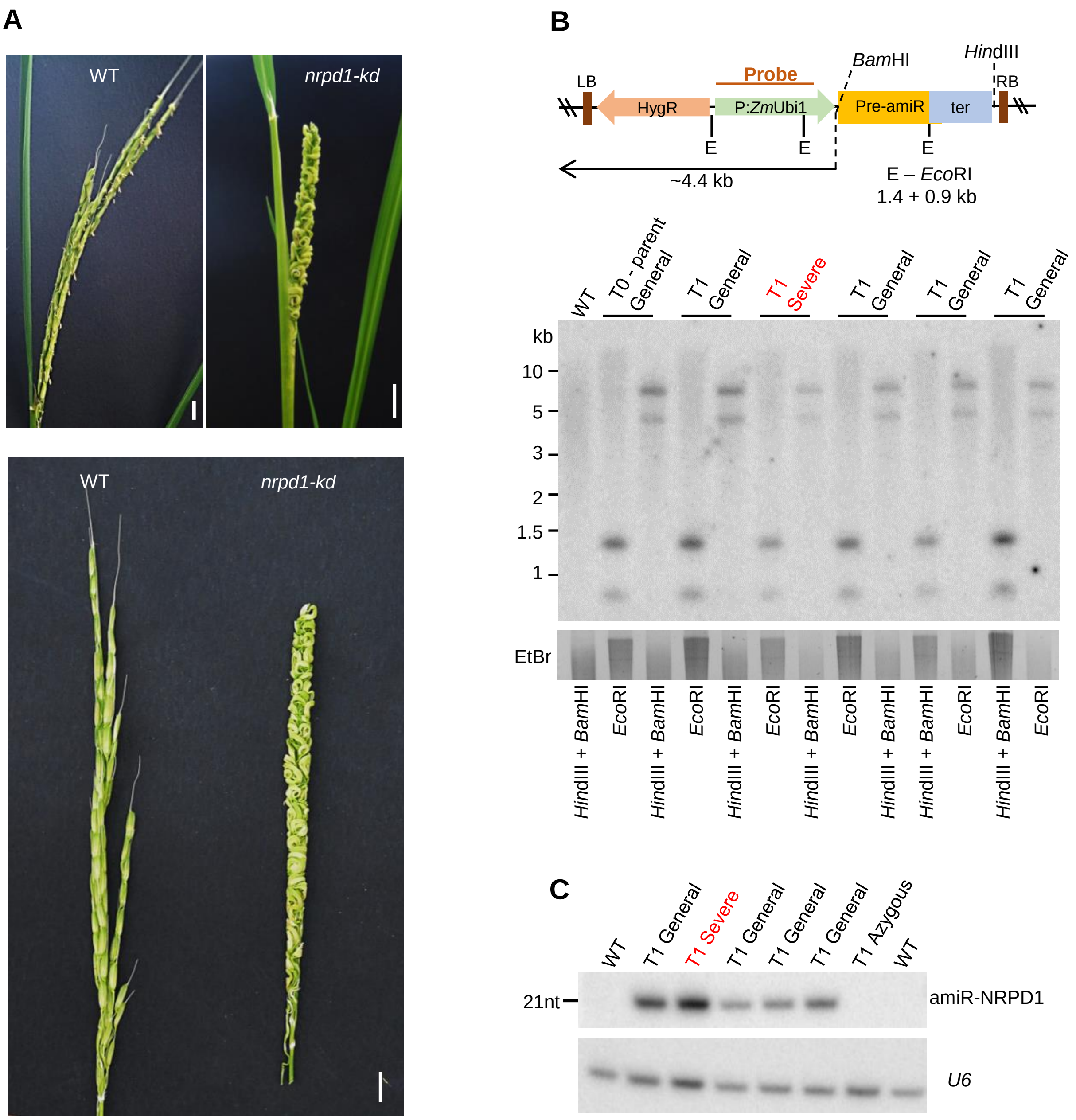


**Supplemental Figure S3.** Specific lineage of Pol IV knockdown plants showed severe defects that are not due to transgenesis or knockdown dosage suggesting possible epimutation. *(A)* Phenotype of panicle displaying the severe defect in T1 generation (T1 severe) that is more drastic than the other offsprings of the same parent. Scale bar: 1 cm. *(B)* DNA blots showing the T-DNA profiles of the T1 severe line compared to that of the parent and other offsprings of the same parent that showed characteristic (general) *kd* like phenotypes. T-DNA map shows the location of the restriction sites used and the expected band sizes upon enzymatic digestion. *(C)* sRNA northern blots showing the levels of artificial miRNA (amiR-NRPD1) levels in the severe as well as the general phenotype displaying lines.

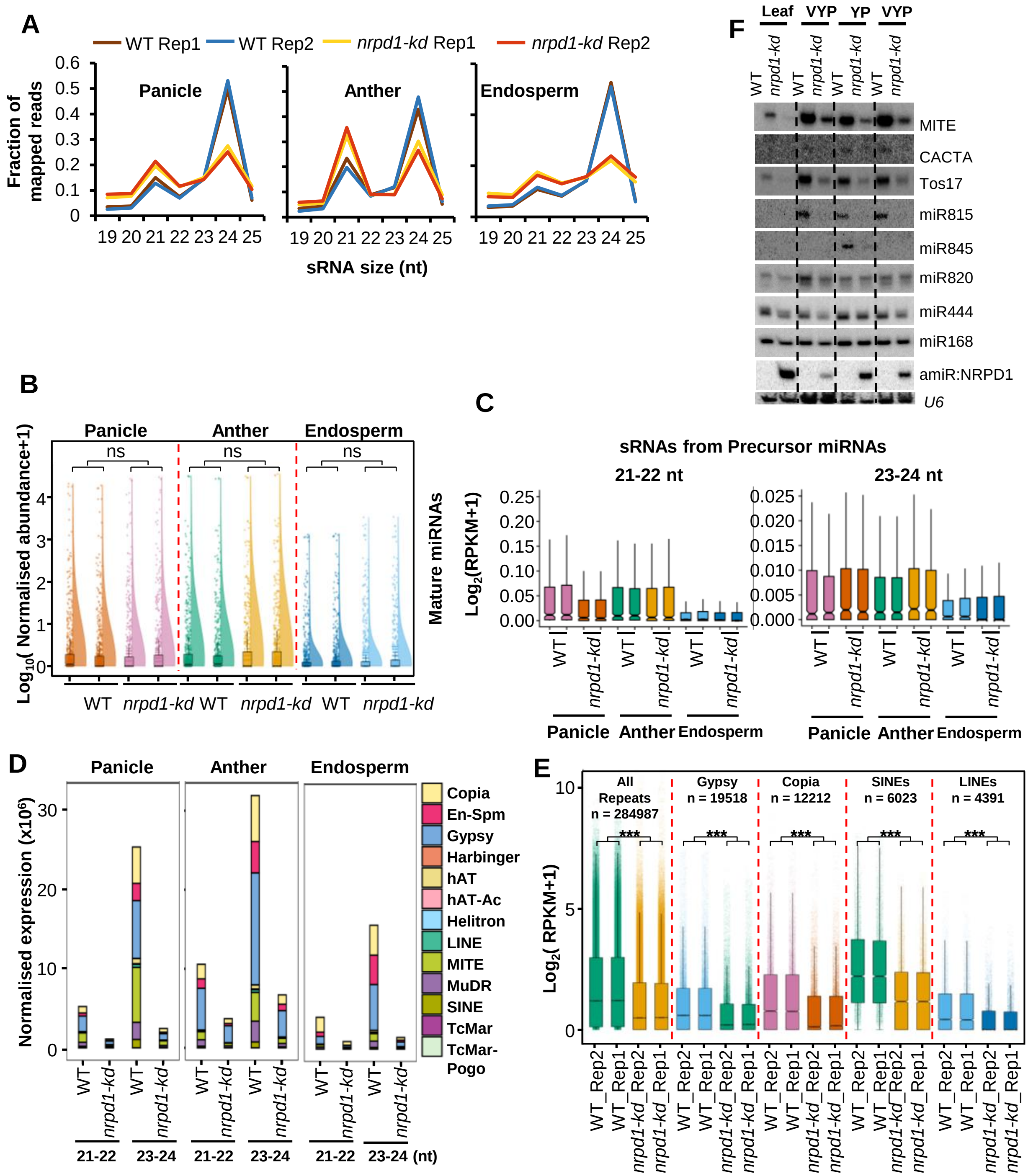


**Supplemental Figure S4.** Loss of pol IV causes specific loss of repeats and transposon associated sRNAs. *(A)* Comparison of relative accumulation of sRNAs of different sizes profiled by small RNA sequencing performed in replicates from different tissues. *(B)* Normalised levels of mature miRNAs in the different tissues, in replicates, of WT and *kd* plants as estimated by miRProf. Rain cloud plots represent the median and the distribution of the datapoints (n = 633). (Wilcoxon test; ns:p-value > 0.05). *(C)* Boxplots showing the 21-22 nt and 23-24 nt sRNA abundance counted over the miRbase annotated miRNA precursor loci from rice (n = 586). *(D)* Abundance of small RNAs from different transposon categories. 21-22 nt and 23-24 nt sizes of sRNAs from corresponding replicates were merged. *(E)* Abundance of sRNAs from different TE categories in replicates. (Wilcoxon test; *** p-value < $2.2 \times 10^{-16}$). *(F)* sRNA northern blots showing the abundance of sRNAs of different categories (miRNAs, siRNAs and TE derived miRNA845) in WT and *kd* tissues – leaves, very young panicle (VYP) and pre-emerged young panicle (YP).

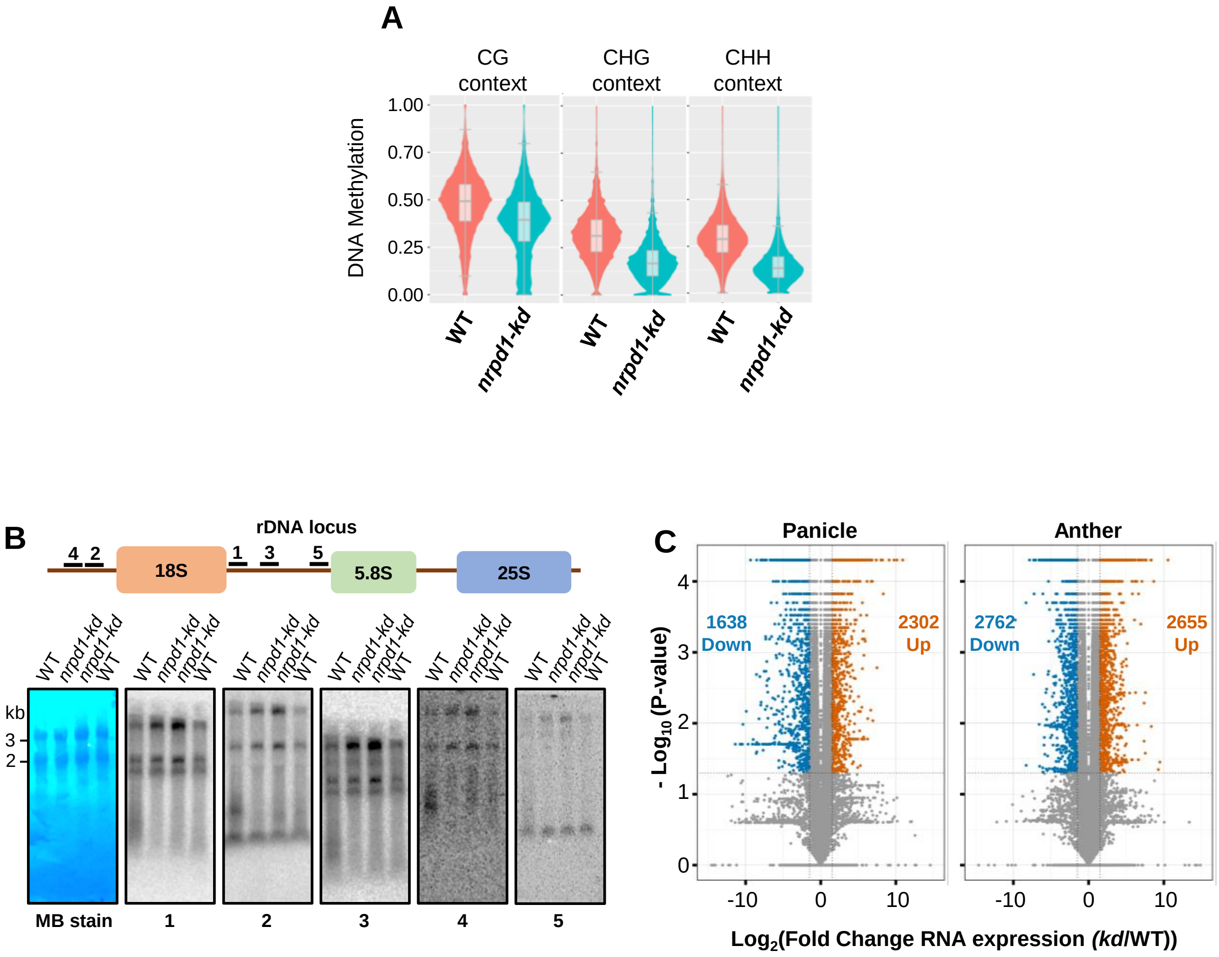


**Supplemental Figure S5**. RNA pol IV is responsible for maintaining DNA methylation, repeat silencing and contributes to gene regulation. *(A)* Box-violin plots showing DNA methylation levels at transposons and repeat features (285215 loci) in WT and kd lines. *(B)* Northern blots showing the abundance of rRNA precursors from WT and *kd* panicle. Methylene blue (MB) stained membrane is shown as loading control. *(C)* Volcano plots showing the number of significantly upregulated ($Log_2FC > 2$) and downregulated ($Log_2FC < 2$) genes in *kd* compared to WT in panicle and anther.

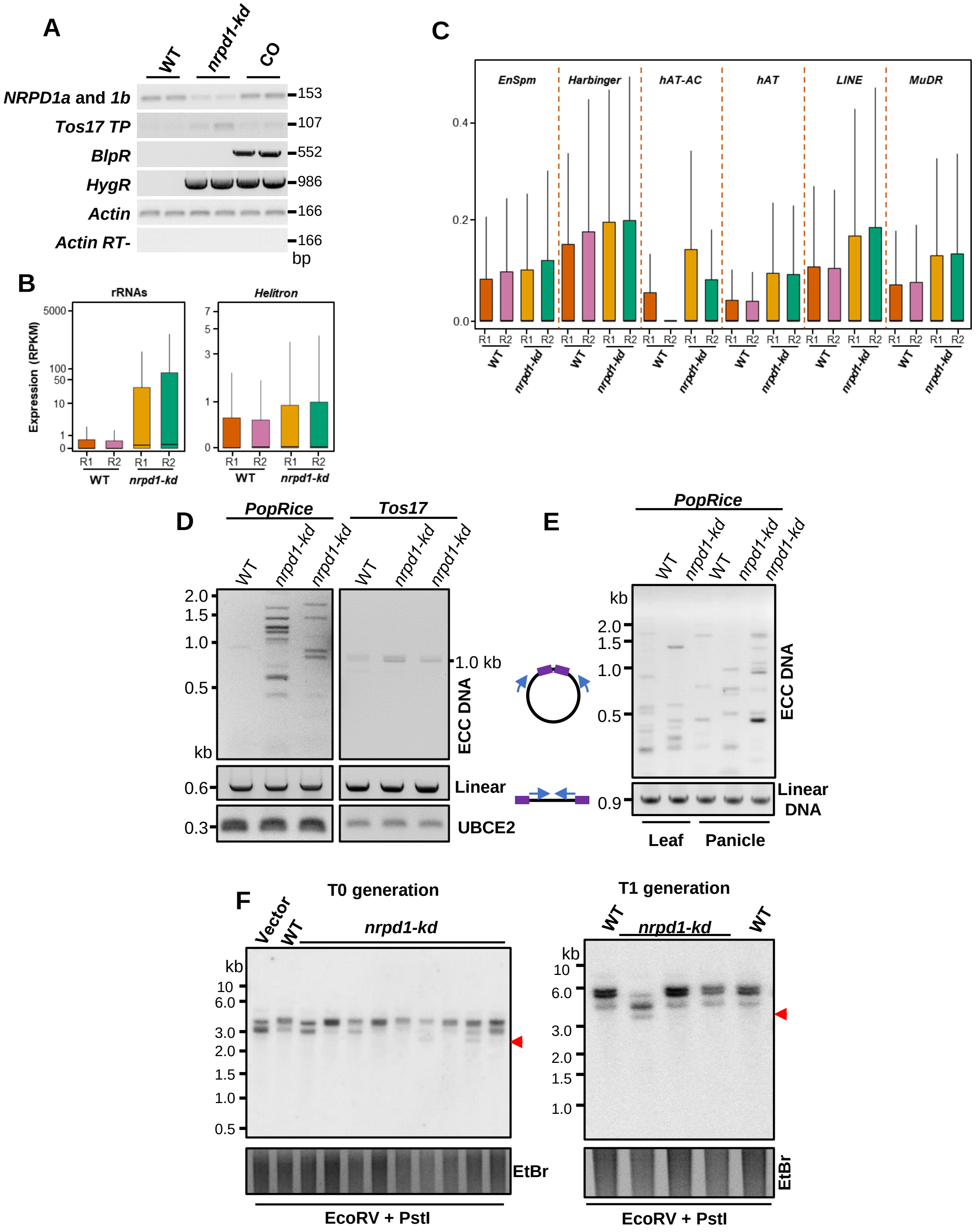


**Supplemental Figure S6.** Pol IV silences transposons and repeats. *(A)* Semi-quantitative RT-PCRs showing levels of the selection markers (HPTII-*HygR* and Bialaphos resistance-*BlpR*), *NRPD1a* and *NRPD1b* (primers binding a conserved region) and *Tos17 transposase (Tos7 TP)* in WT, *kd* and NRPD1b complementation (CO). *(B-C)* Boxplots showing the normalised abundance of transcripts from annotated repeat loci in the replicates (R1 and R2) of WT and *kd* plants. The Y-axis is scaled to inverse sine hyperbolic function of RPKM values. *(D)* PCRs showing accumulation of extrachromosomal circular (ECC) DNA intermediates of TEs *PopRice* and *Tos17* with intact DNA from panicle. UBCE2 fragment is used as linear DNA loading control. *(E)* PCRs showing accumulation of extrachromosomal circular (ECC) DNA intermediates of *PopRice* DNA from panicle pre-digested with Plasmid-safe$^{TM}$ DNase. *(F)* Transposon copy number Southern blot showing copy number variation (arrows) of LINE1 transposon in both T0 and T1 generations in *kd*. Ethidium bromide (EtBr) staining served as control.

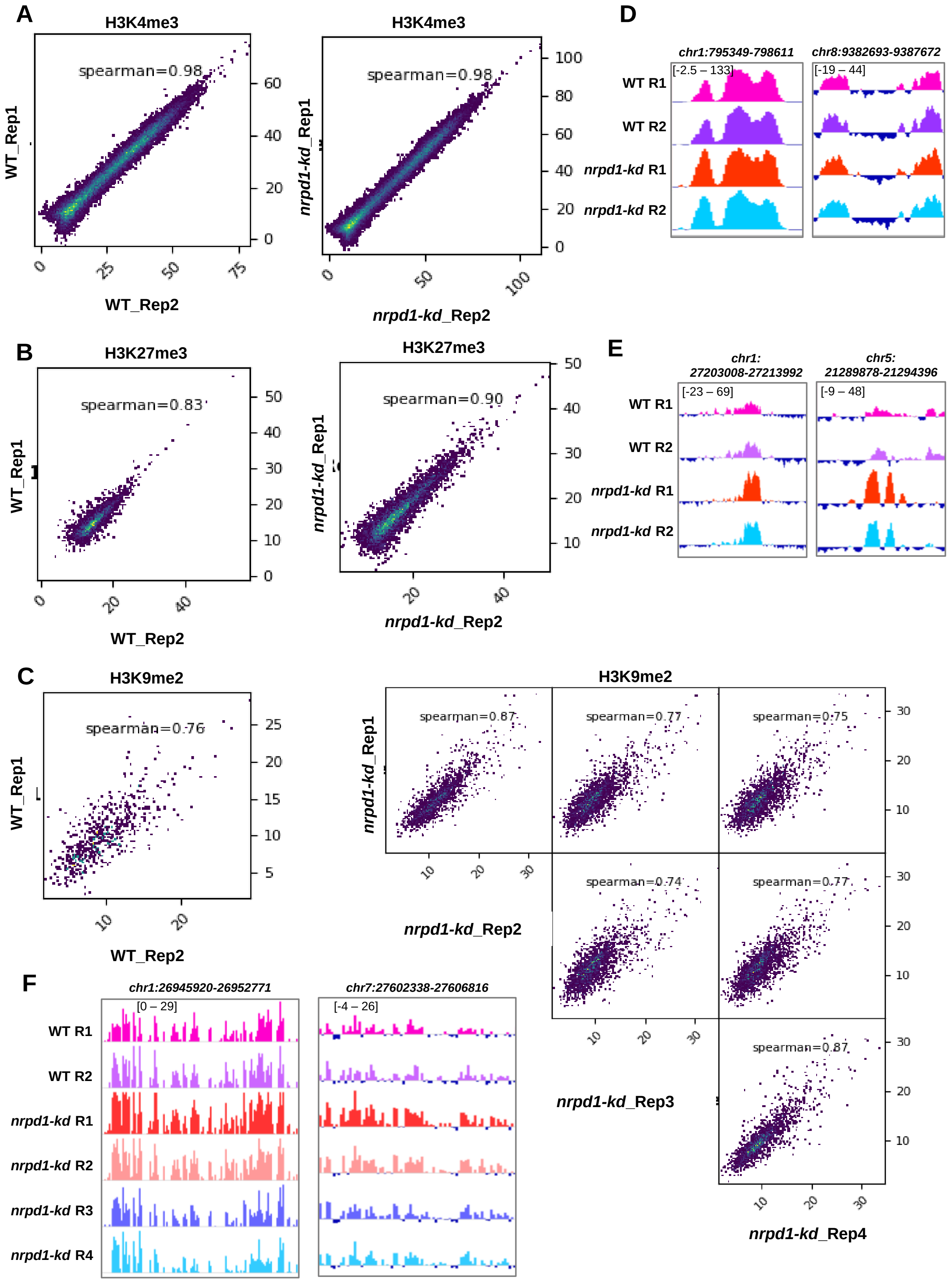


**Supplemental Figure S7.** ChIP – Sequencing exhibits concordant signals across replicates. *(A-C)* Spearman correlation scatter plots showing the ChIP signals normalised to H3 (ChIP-H3) over the combined sets of peaks identified by MACS2 in WT and *nrpd1-kd* for H3K4me3 *(A)*, H3K27me3 *(B)* and H3K9me2 *(C)*. Spearman correlation co-eficients calculated are inside the plots for pairwise comparisons. *(D-F)* Genome browser screen shots of H3K4me3 *(D)*, H3K27me3 *(E)* and H3K9me2 *(F)* occupancy with normalised signals (ChIP-H3) depicted.

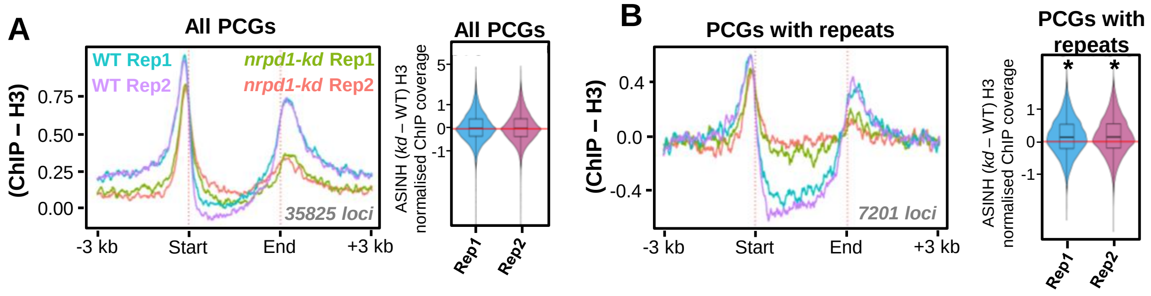


**Supplemental Figure S8.** Loss of pol IV impacts occupancy of pol II. *(A)* Metaplot depicting pol II occupancy over rice protein coding genes normalised to the H3 ChIP. Numbers in grey describe the number of loci taken for analyses. Box-violin plots shows the difference in enrichment in *kd* compared to WT over the described sets of loci. The Y-axis is scaled to inverse sine hyperbolic function of enrichment values. *(B)* Metaplot depicting pol II occupancy over genes having at least 10% of their length with annotated repeats normalised to the H3 ChIP. Numbers in grey describe the number of loci taken for analyses. Box-violin plots shows the difference in enrichment in *kd* compared to WT over the described sets of loci. The Y-axis is scaled to inverse sine hyperbolic function of enrichment values. Mann-Whitney U test, * p-value < 0.0001.

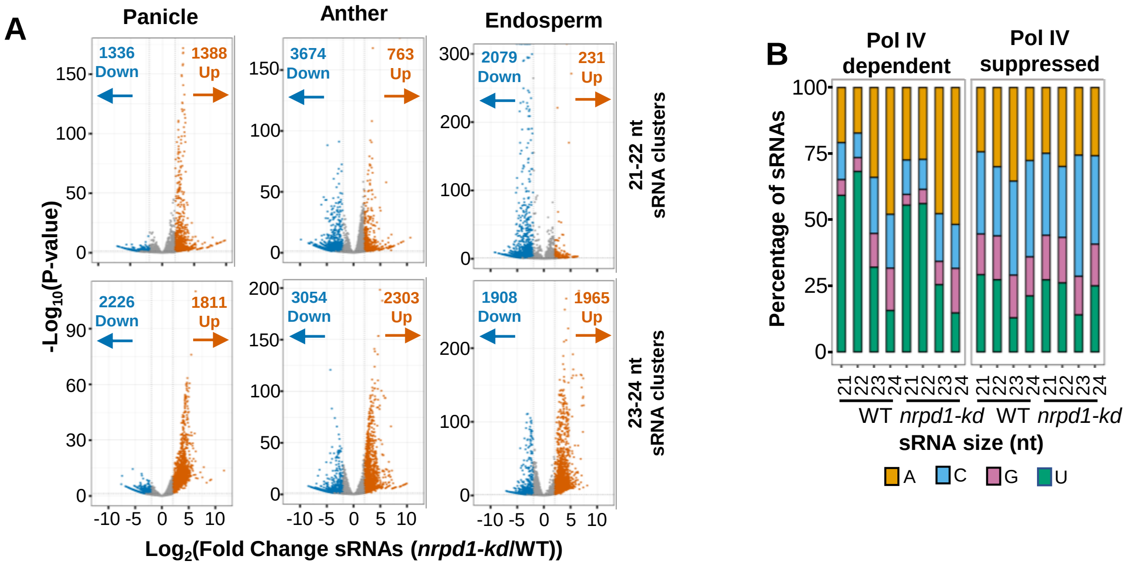


**Supplemental Figure S9.** Pol IV complex suppresses peculiar sRNA production from several loci. *(A)* Volcano plots showing levels of deregulation of sRNAs over clusters identified by shortstack from panicle, anther and endosperm sRNA datasets. The clusters were identified after subcategorizing into 21-22 nt and 23-24 nt size classes. Clusters showing statistically significant difference of 4-folds relative to WT are highlighted. The number of up- and downregulated clusters are mentioned with arrows. *(B)* Stacked bar plots showing normalised abundance of rice sRNAs of different sizes from pol IV suppressed and dependent bins displaying the 5' nucleotide bias.

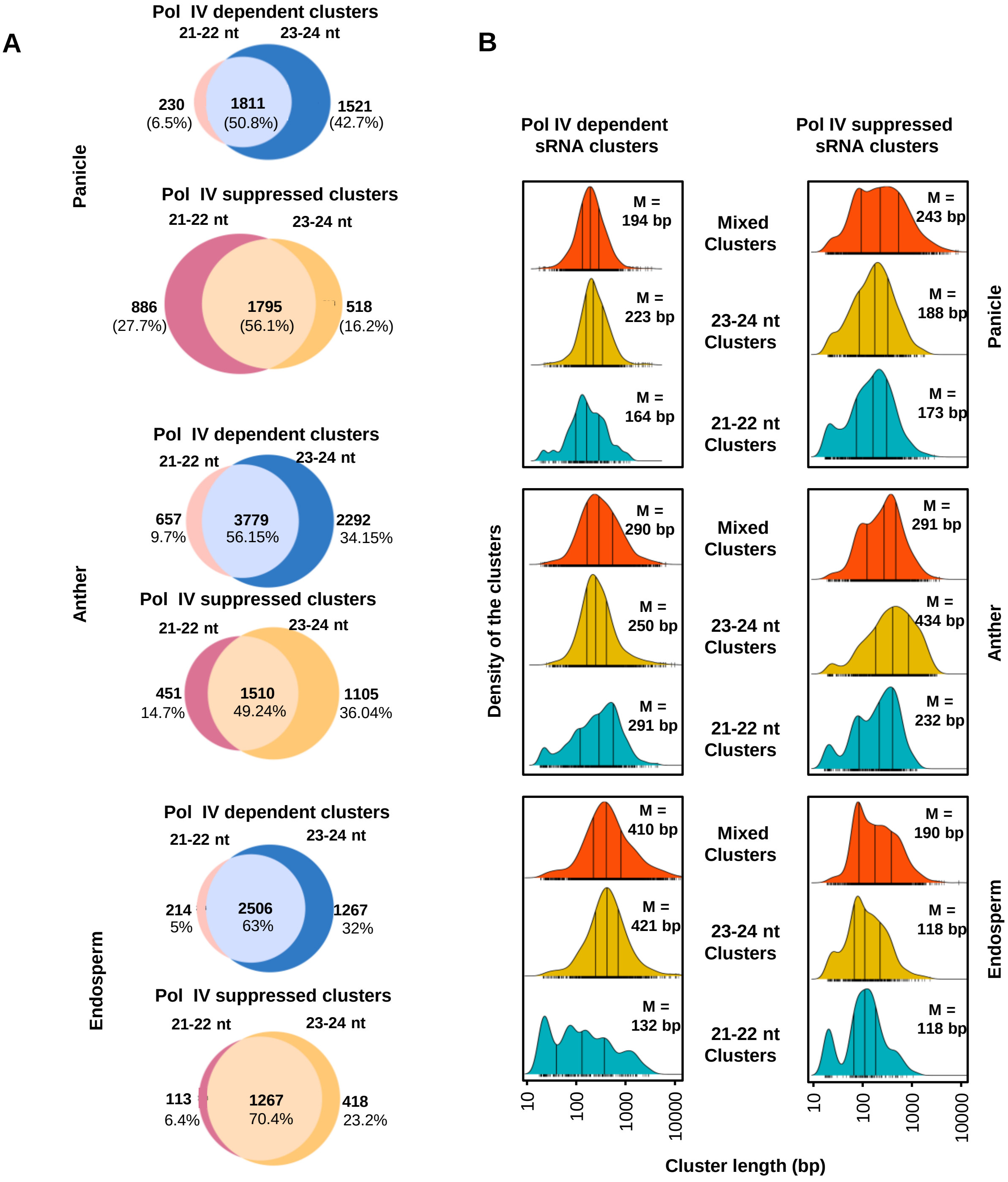


**Supplemental Figure S10.** Pol IV dependent and pol IV suppressed sRNA clusters are of similar length. *(A)* Venn plots showing sRNA size enrichment of pol IV dependent and suppressed clusters. Clusters are classified as 21-22nt (clusters with more than 65% sRNAs in 21-22nt size class), 23-24nt (clusters with more than 65% sRNAs in 23-24nt size class) and mixed clusters (all the other clusters with mixed size profile of sRNAs). *(B)* Density distribution plots of the sizes of shortstack identified sRNA clusters sub-categorized into pol IV suppressed and dependent clusters. Central line within the peak denotes the median length of the clusters and other two lines represent the interquartile range. Median length is denoted in the plots as M.

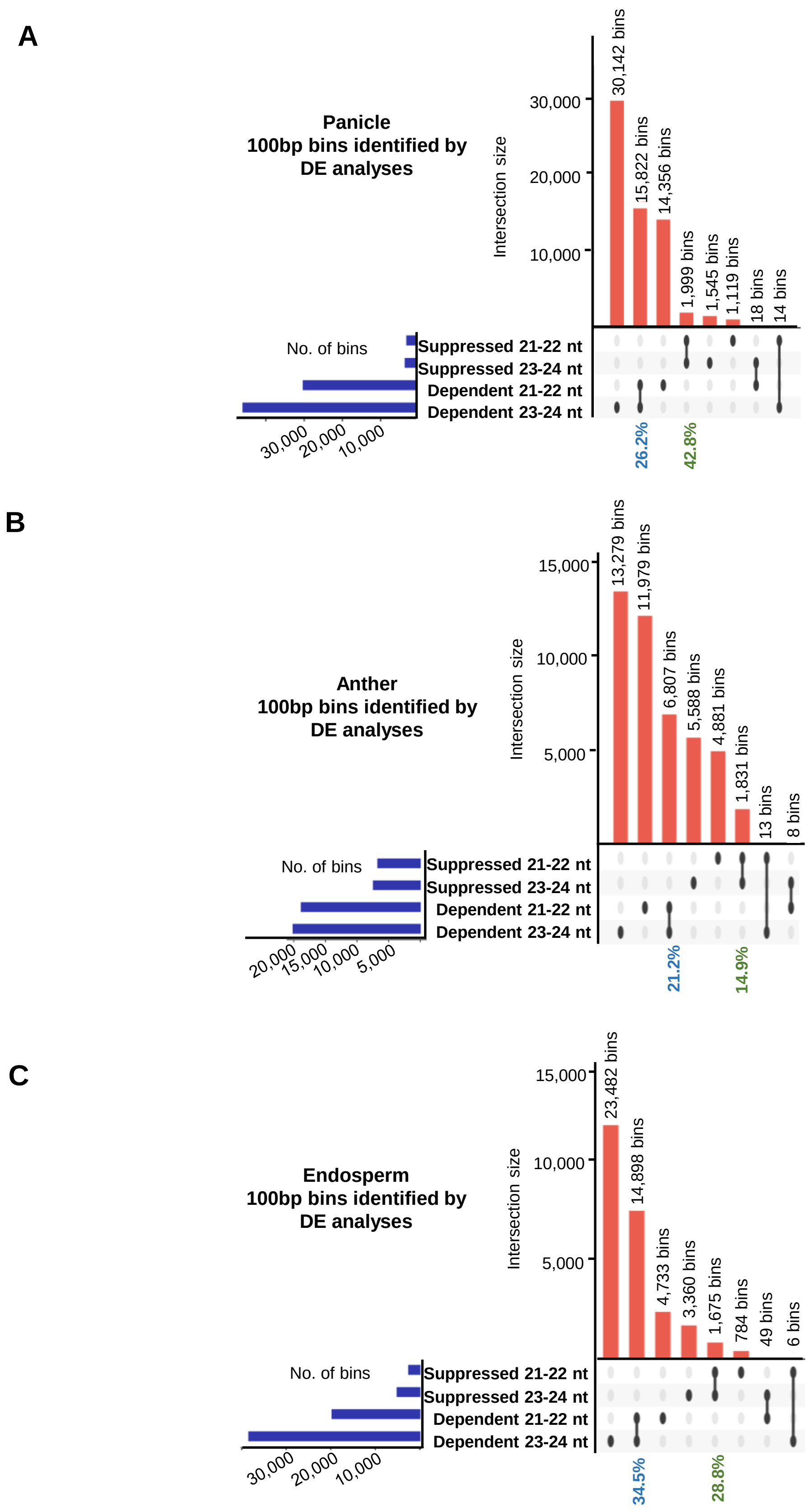


**Supplemental Figure S11.** Pol IV dependent and pol IV suppressed sRNA bins are distinct in producing sRNAs with specific sizes. *(A-C)* Upset plots describing the numbers and overlap of differentially expressed sRNA bins categorized as pol IV suppressed and dependent bins in panicle *(A)*, anther *(B)* and endosperm *(C)*.

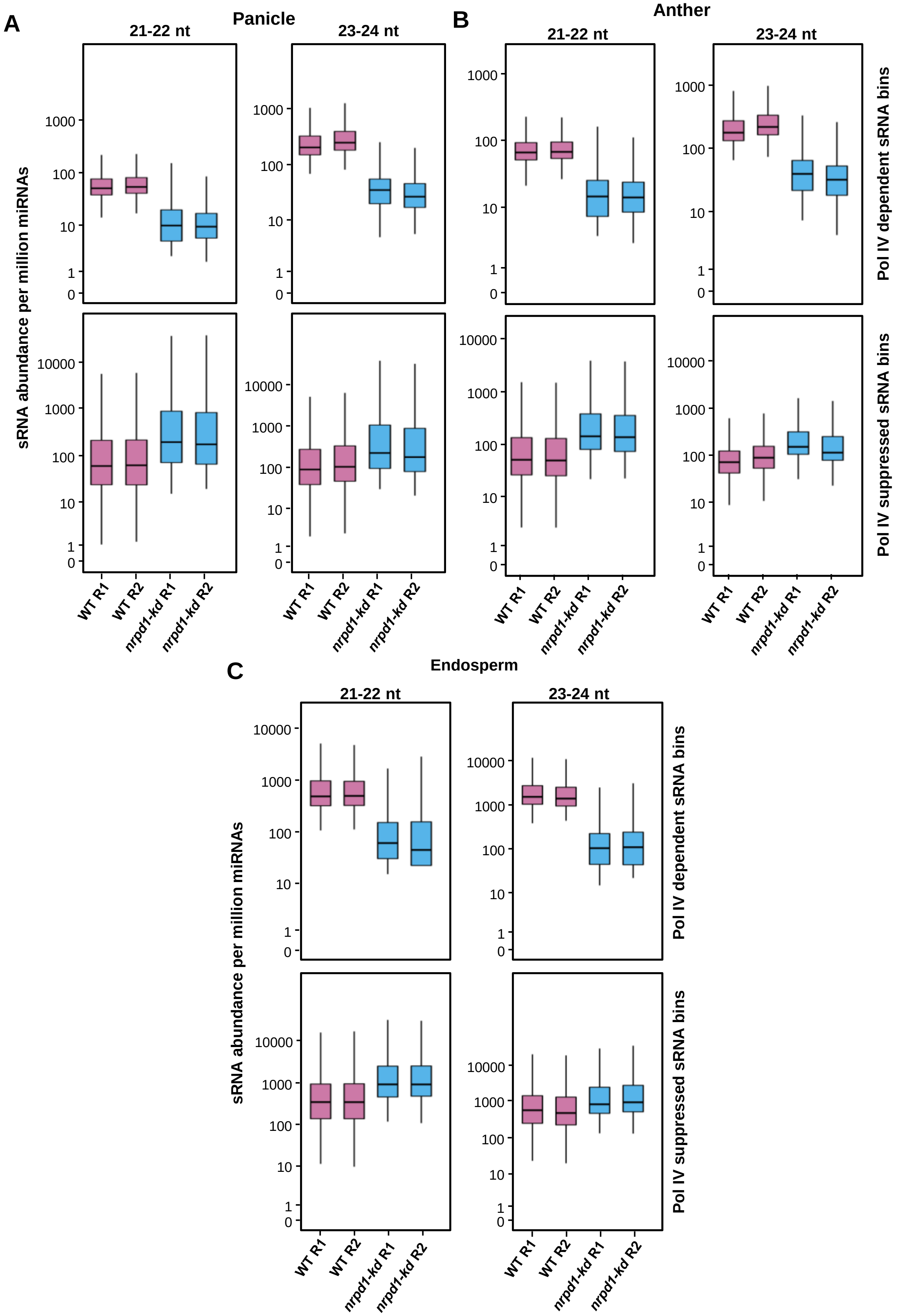


**Supplemental Figure S12.** Pol IV dependent and pol IV suppressed sRNA bins are not due to oversampling and library normalisation. *(A-C)* Boxplots describing the relative abundance of sRNAs normalised to net raw abundance of miRProf identified miRNAs in pol IV suppressed and dependent bins from panicle *(A)*, anther *(B)* and endosperm *(C)*. The Y-axis is scaled to inverse sine hyperbolic function of values.

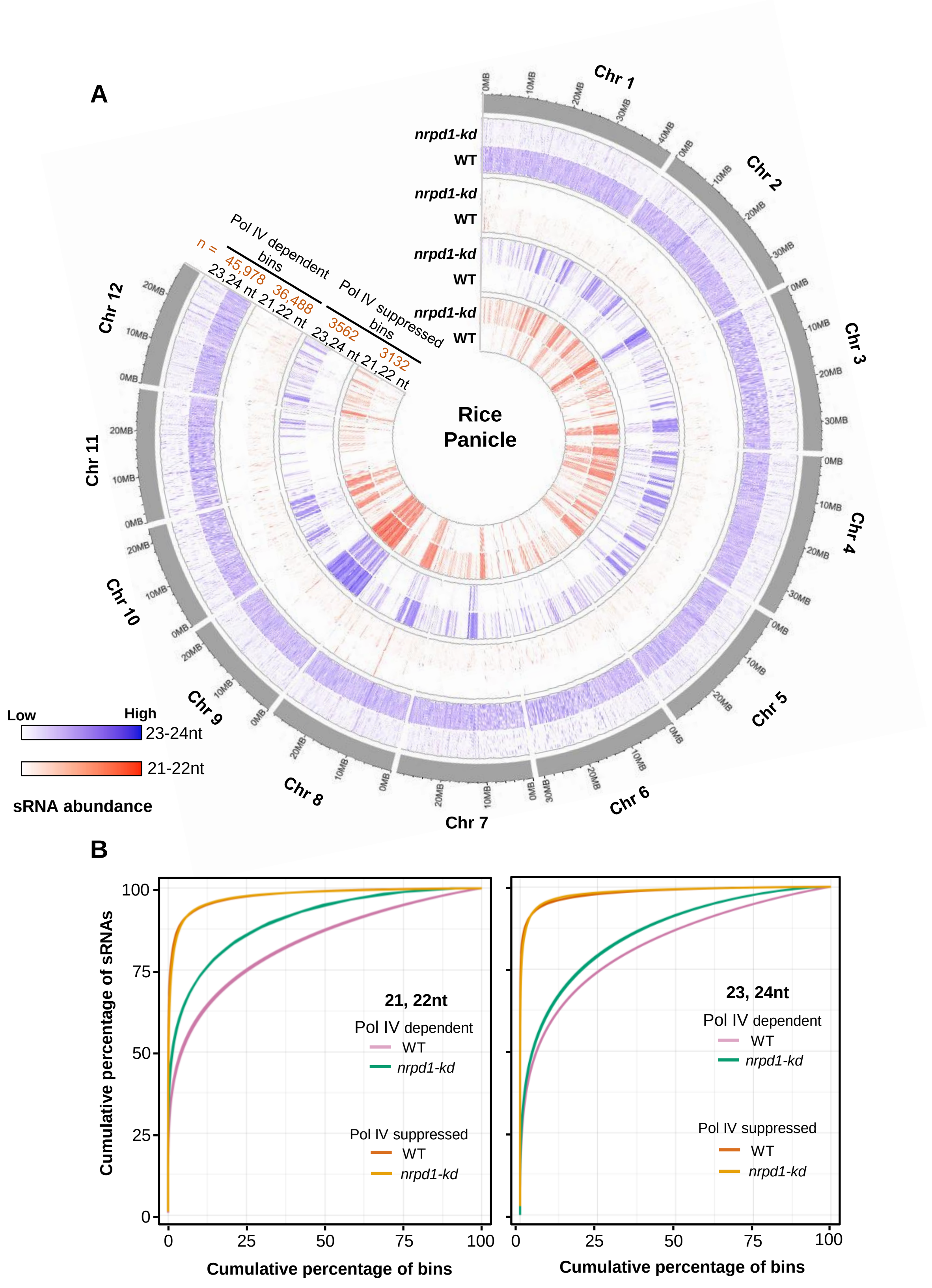


**Supplemental Figure S13.** Pol IV suppressed sRNAs from panicle are non-uniformly distributed across the genome. *(A)* Circos plot showing the normalised abundance and distribution of sRNAs in 100 bp windows categorised as pol IV dependent and pol IV suppressed bins across 12 chromosomes. The abundance values of replicates are merged and displayed as 21,22nt and 23,24nt tracks. The number of bins identified for each category is labelled as n. *(B)* Percentage cumulative sum plots for 21,22nt and 23,24nt size classes of sRNAs from panicle subclassified into pol IV suppressed and dependent sRNA bins. The replicate lines are identically coloured.

A

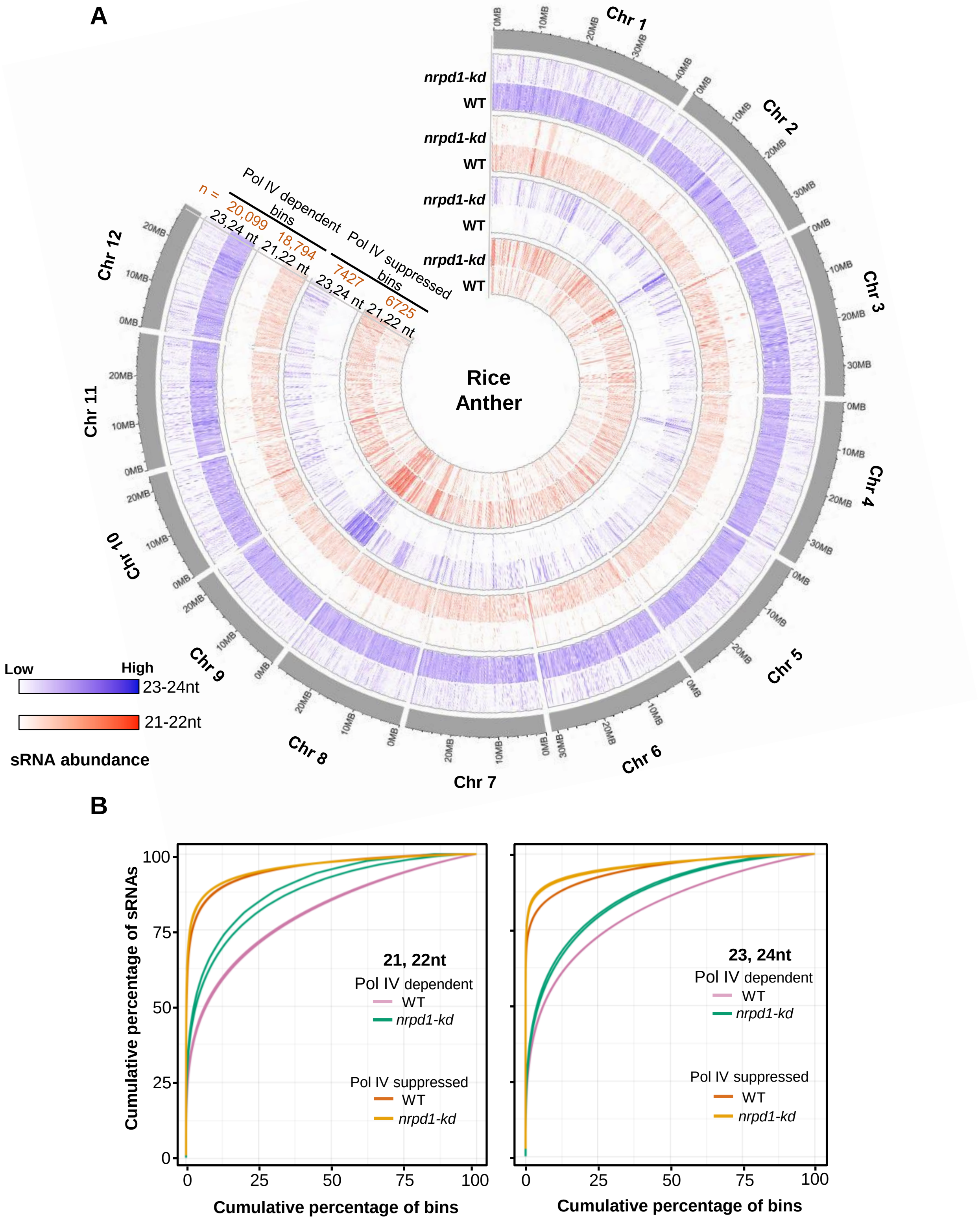


B



**Supplemental Figure S14.** Pol IV suppressed sRNAs from anther are non-uniformly distributed across the genome. *(A)* Circos plot showing the normalised abundance and distribution of sRNAs in 100 bp windows categorised as pol IV dependent and pol IV suppressed bins across 12 chromosomes. The abundance values of replicates are merged and displayed as 21,22nt and 23,24nt tracks. The number of bins identified for each category is labelled as n. *(B)* Percentage cumulative sum plots for 21,22nt and 23,24nt size classes of sRNAs from anther subclassified into pol IV suppressed and dependent sRNA bins. The replicate lines are identically coloured.

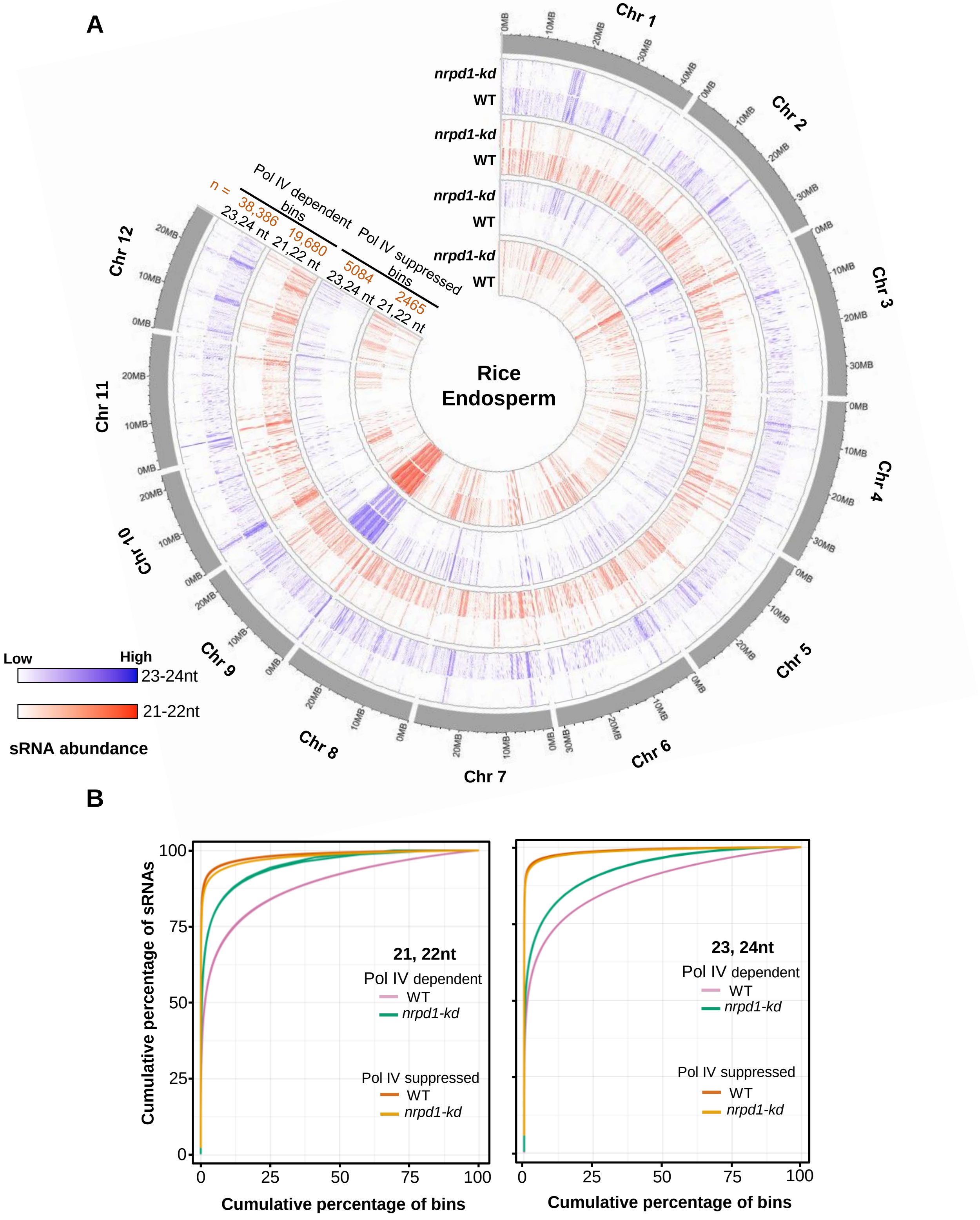


**Supplemental Figure S15.** Pol IV suppressed sRNAs from endosperm are non-uniformly distributed across the genome. *(A)* Circos plot showing the normalised abundance and distribution of sRNAs in 100 bp windows categorised as pol IV dependent and suppressed bins across 12 chromosomes. The abundance values of replicates are merged and displayed as 21,22nt and 23,24nt tracks. The number of bins identified for each category is labelled as n. *(B)* Percentage cumulative sum plots for 21,22nt and 23,24nt size classes of sRNAs from endosperm subclassified into pol IV suppressed and dependent sRNA bins. The replicate lines are identically coloured.

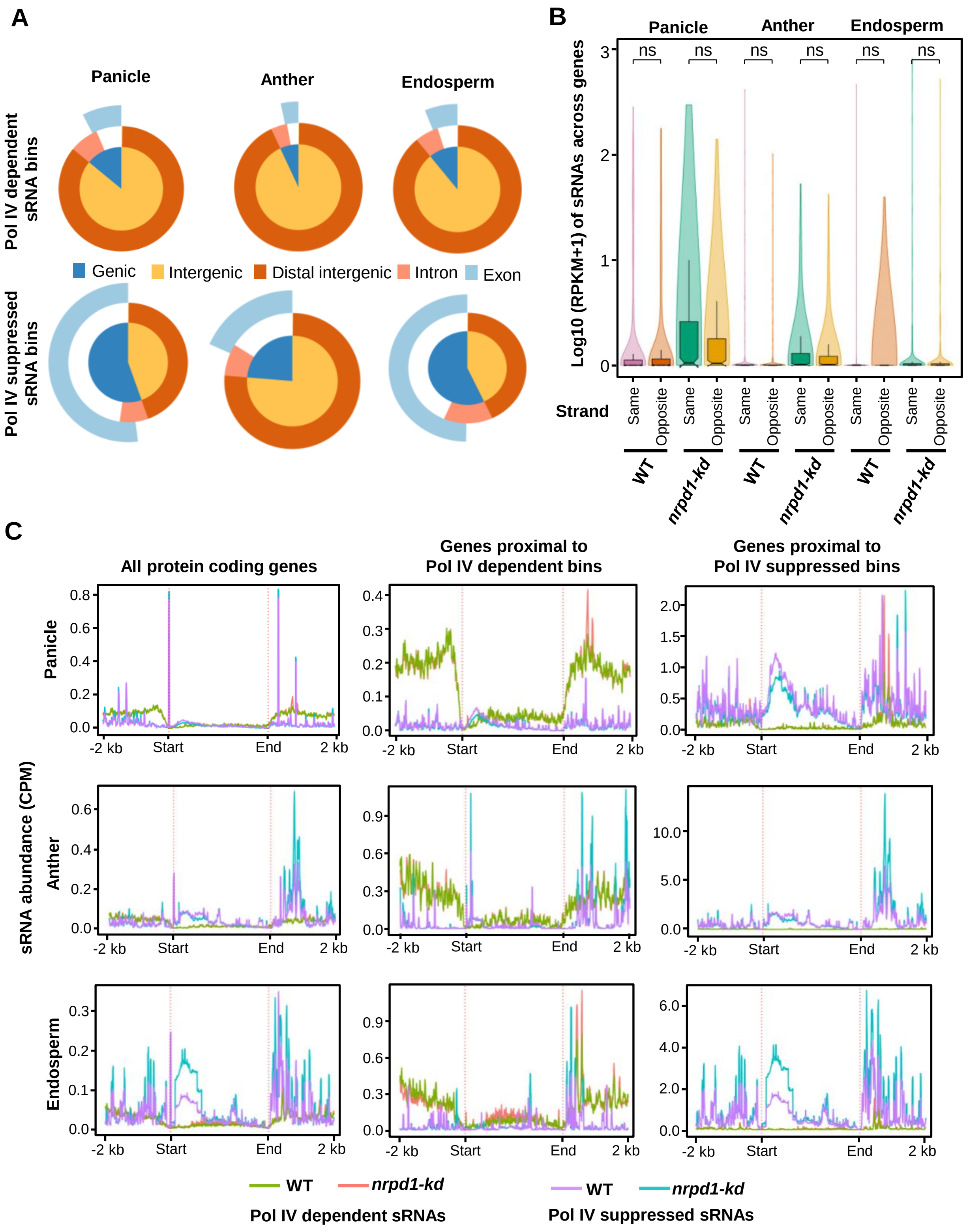


**Supplemental Figure S16.** Pol IV suppressed sRNAs have significant overlap with coding regions. *(A)* Vennpie plots showing the overlap of the pol IV suppressed and dependent sRNA bins in major genomic features as identified by ChIPseekeR annotation. *(B)* Boxplots showing the distribution of pol IV suppressed sRNAs across genes overlapping with the pol IV suppressed bins. sRNAs are counted with respect to gene coding strand. Wilcoxon signed rank test was used for statistical testing. ns - not-significant (p-value > 0.05). The box shows the interquartile range and violin represent the distribution of datapoints. *(C)* Metaplots representing the coverage of sRNAs (pol IV dependent and suppressed sRNAs) across all the protein coding genes, genes proximal to dependent bins and genes proximal to suppressed bins identified in panicle, anther and endosperm.

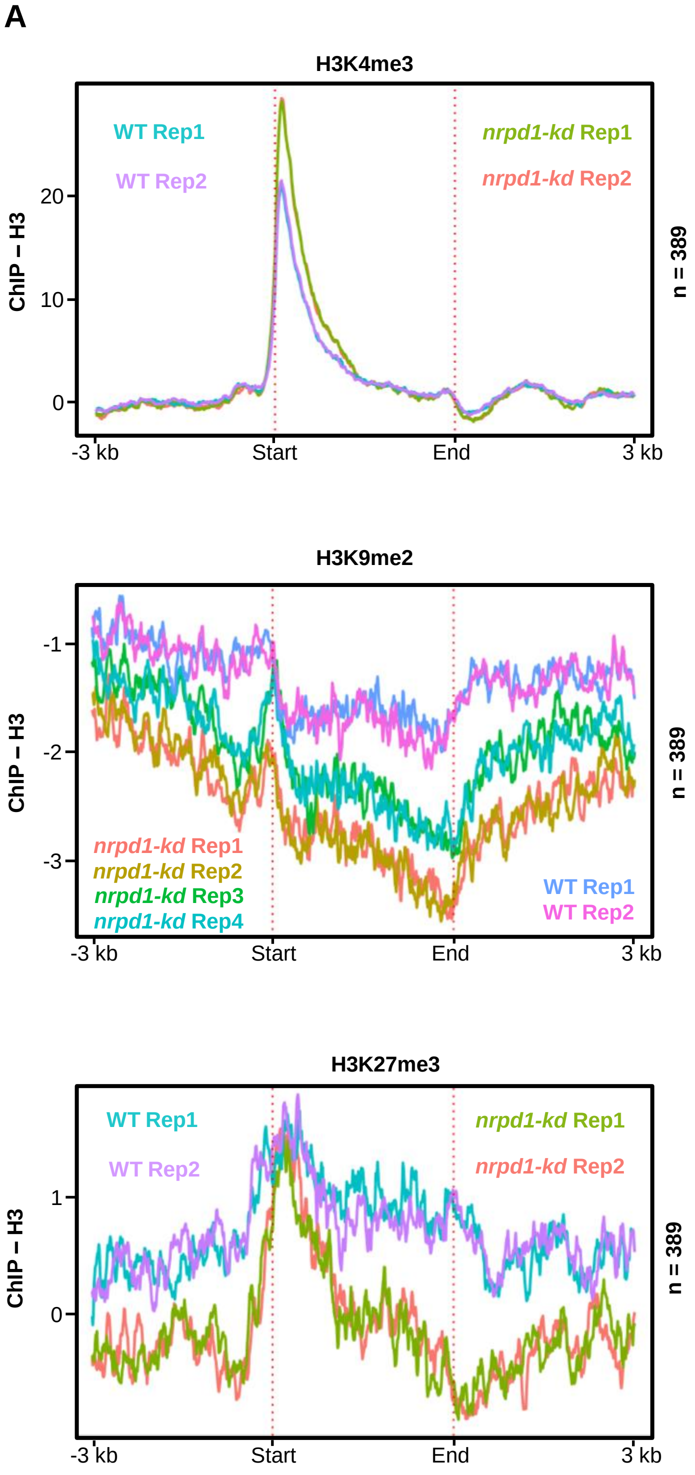


**Supplemental Figure S17.** Genes overlapping with suppressed sRNAs exhibit reduced silencing compensation by H3K27me3. *(A)* Metaplots describing the occupancy of histone H3 modifications H3K4me3, H3K9me2 and H3K27me3 normalised to total histone H3 occupancy plotted over the proteins overlapping with the suppressed sRNA bins (n = 389) from panicle.

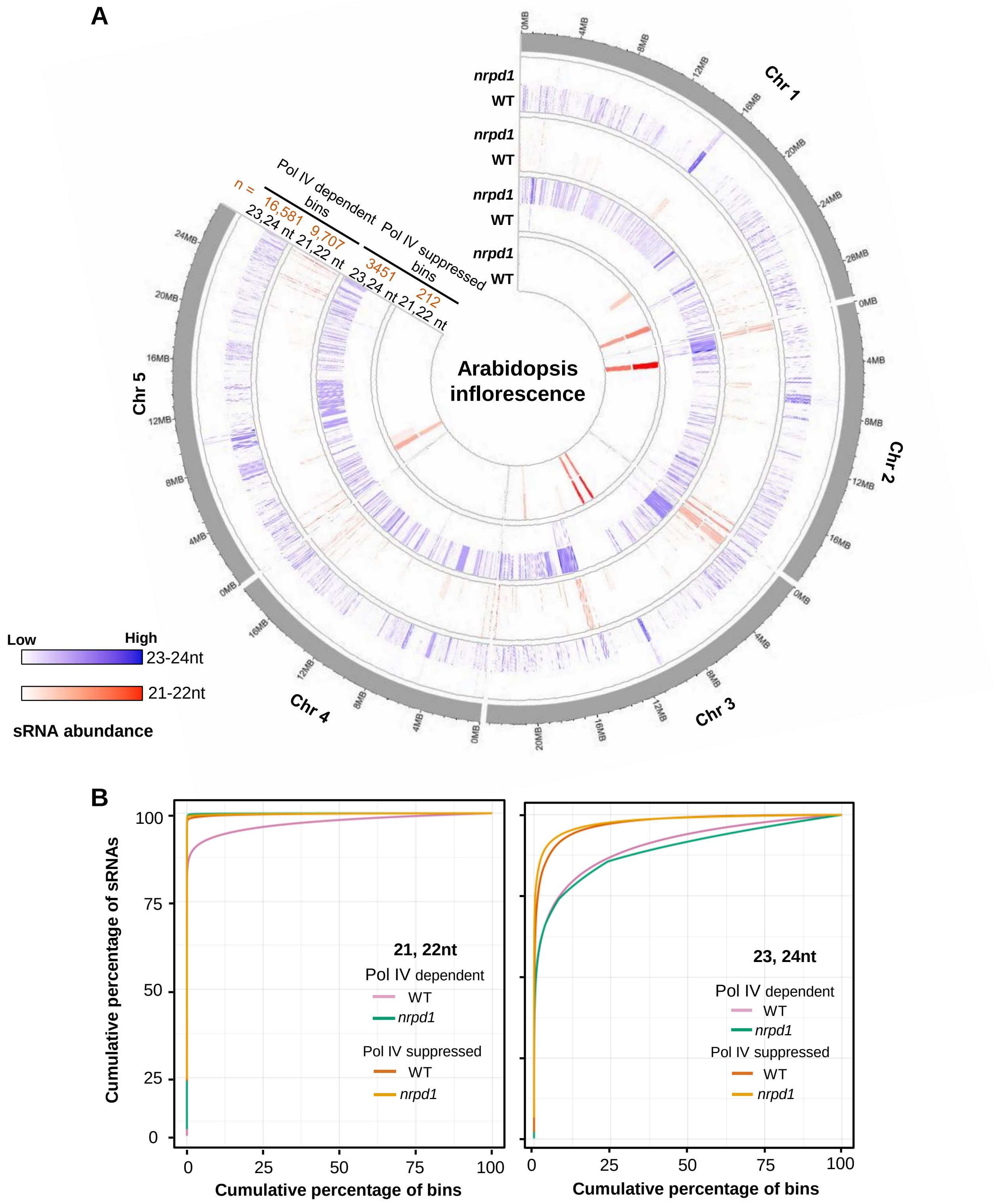


**Figure 18.** Pol IV suppressed sRNAs from Arabidopsis inflorescence are non-uniformly distributed across the genome similar to rice tissues. *(A)* Circos plot showing the normalised abundance and distribution of sRNAs in 100 bp windows categorised as pol IV dependent and suppressed bins across 5 chromosomes. The abundance values are displayed as 21,22nt and 23,24nt tracks. The number of bins identified for each category is labelled as n. The datasets are taken from from GSE61439. *(B)* Percentage cumulative sum plots for 21,22nt and 23, 24nt size classes of sRNAs from arabidopsis inflorescence subclassified into pol IV suppressed and dependent sRNA bins.

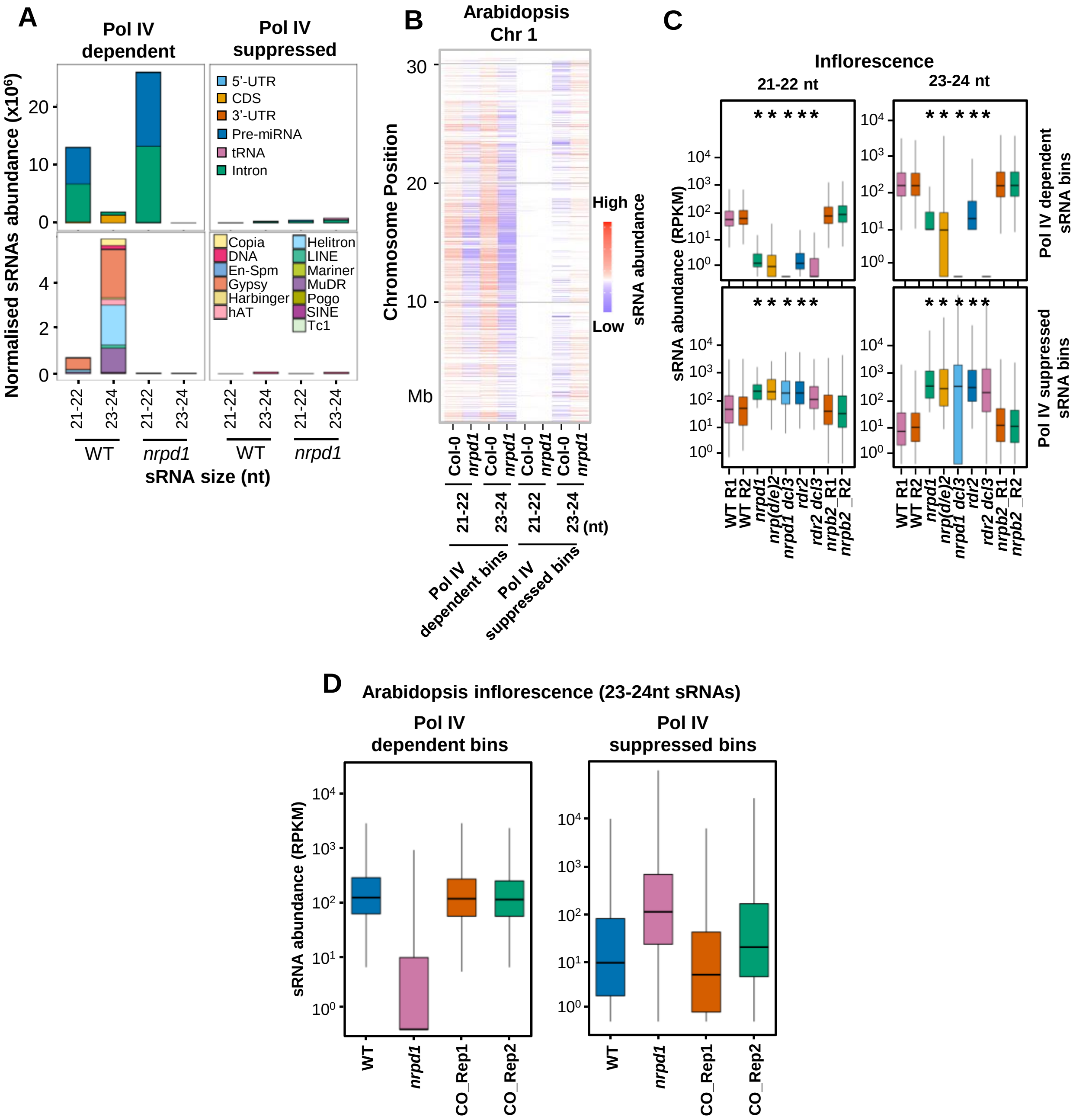


**Supplemental Figure S19.** Pol IV suppressed sRNAs from Arabidopsis are characteristically similar to that of rice. *(A)* Stacked bar plots showing abundance of sRNAs from different features (coding features in the top and transposon categories in the bottom). *(B)* Heatmap showing the normalised abundance of sRNAs from pol IV dependent and suppressed bins identified in the same way as in rice over the Arabidopsis chromosome 1. The datasets from GSE61439 (inflorescence tissue) was analysed and sRNAs from differentially expressed bins were represented. *(C)* Boxplots showing the normalised sRNA abundance of pol IV suppressed and dependent bins from Arabidopsis seedlings tissue taken from GSE72993 and GSE98285. The Y-axis is scaled to inverse sine hyperbolic function of RPKM values. Mann Whitney U test. * p-value < 0.0001. *(D)* Boxplots showing the abundance of 23-24nt sRNAs in WT, *nrpd1* and NRPD1 complemented lines (CO) in the pol IV dependent and suppressed bins identified. Datasets taken from GSE140566. The Y-axis is scaled to inverse sine hyperbolic function of RPKM values.

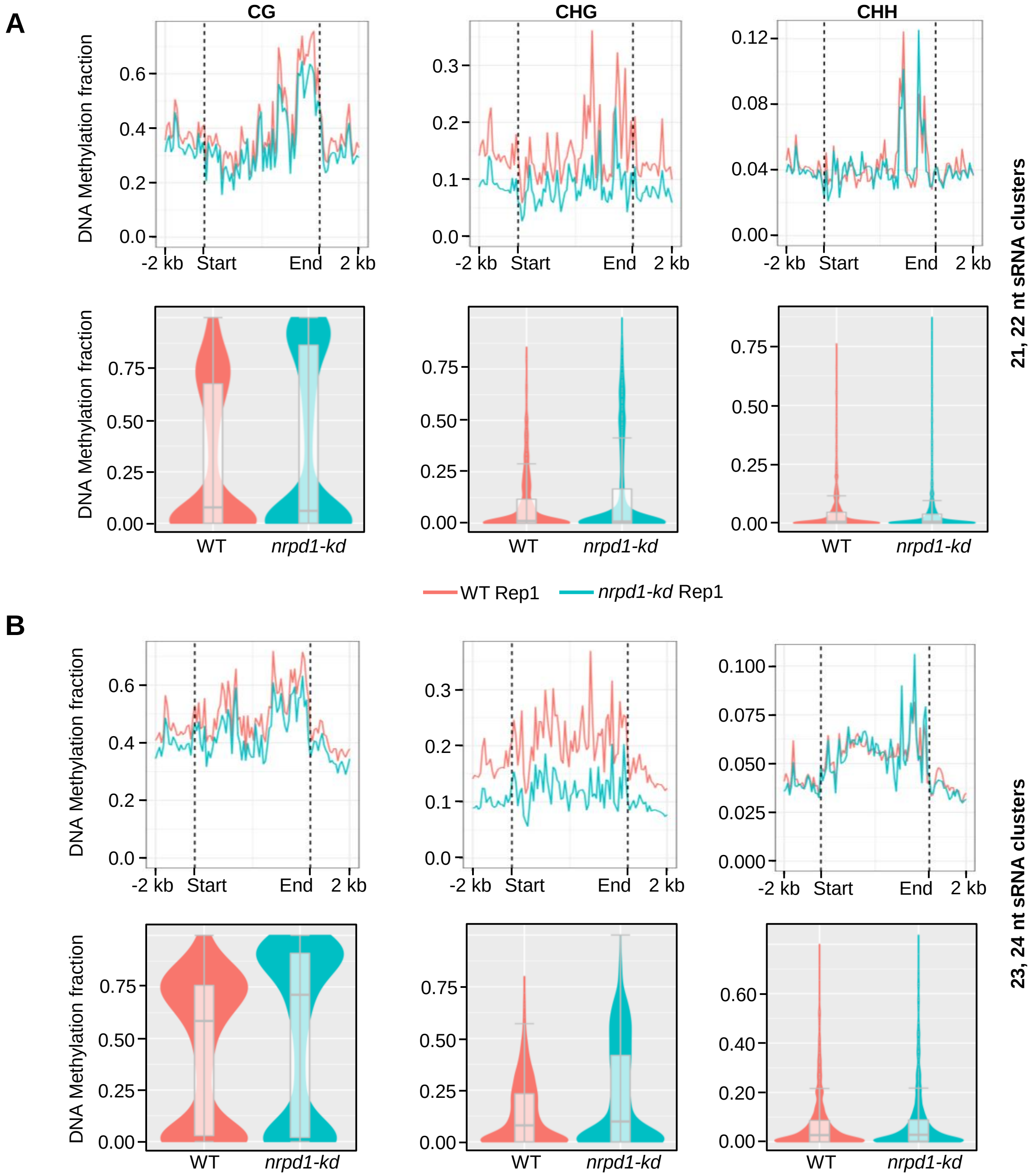


**Supplemental Figure S20.** Canonical RdDM does not potentiate pol IV suppressed sRNAs in rice. *(A-B)* Metaplots and box-violin plots depicting the level of DNA methylation in the panicle tissue over the 21,22nt *(A)* and 23,24nt *(B)* pol IV suppressed panicle sRNA clusters in CG, CHG and CHH contexts.

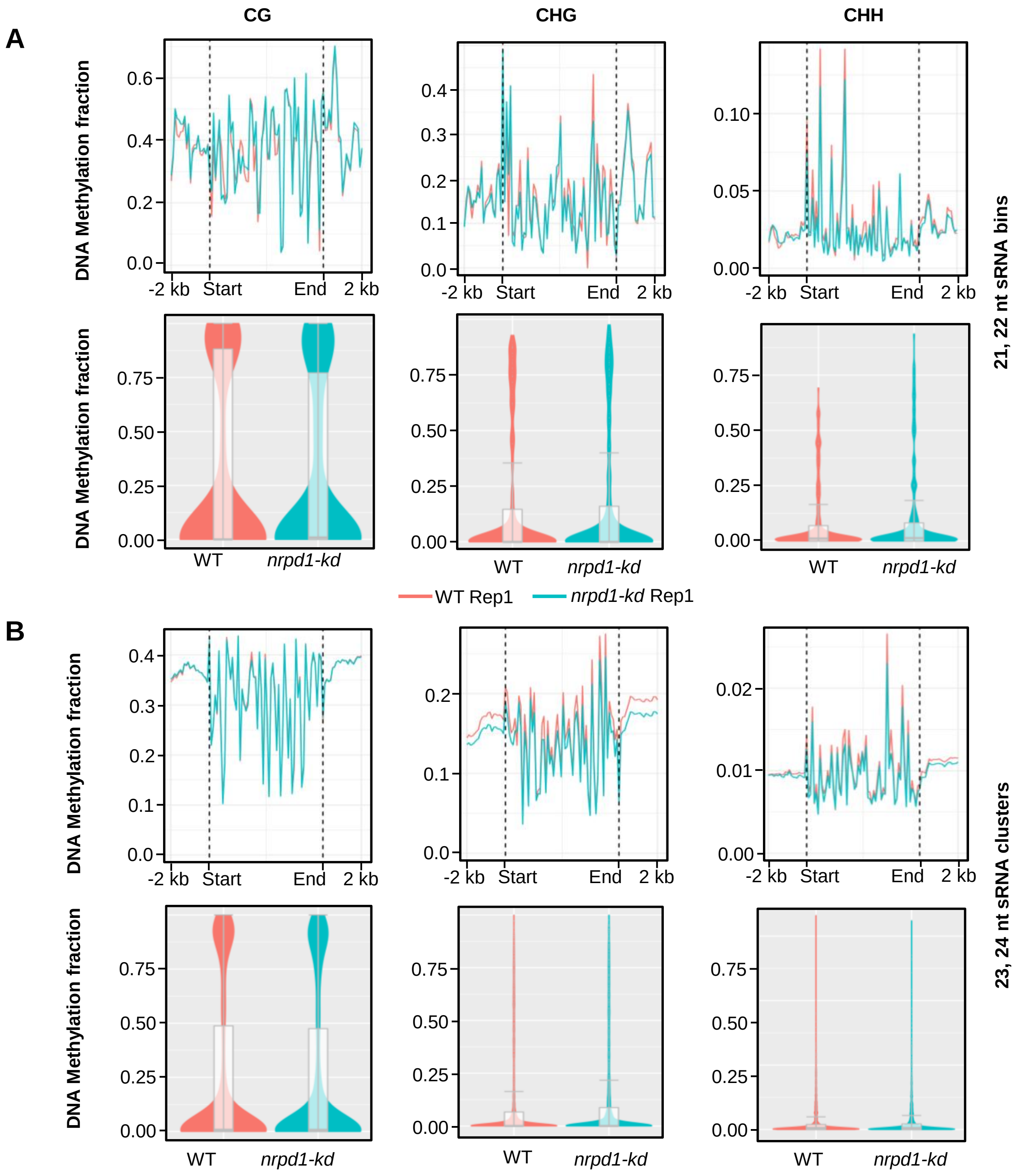


**Supplemental Figure S21.** Canonical RdDM does not potentiate pol IV suppressed sRNAs in Arabidopsis. *(A-B)* Metaplots and box-violin plots depicting the level of DNA methylation over the 21,22nt *(A)* and 23,24nt *(B)* pol IV suppressed panicle sRNA bins in CG, CHG and CHH contexts. The datasets were taken from GSE99689.

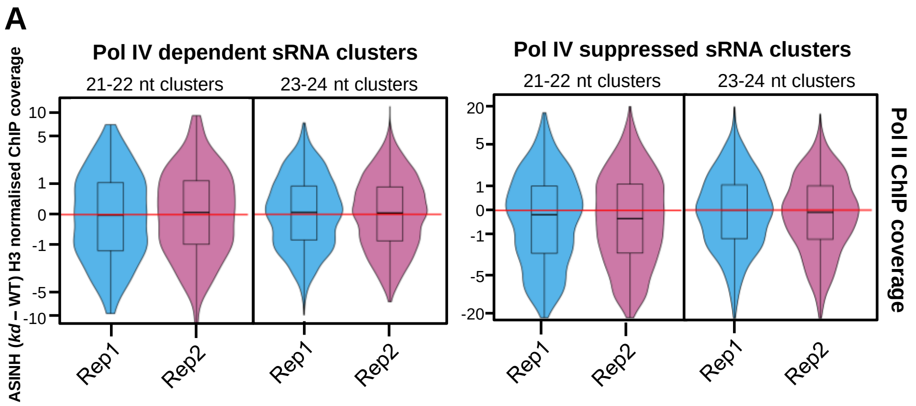


**Supplemental Figure S22**. Net Pol II occupancy over pol IV suppressed sRNA clusters does not change. *(A)* Box-violin plots showing the difference in Pol II coverage over the pol IV dependent and pol IV suppressed sRNA clusters size classified into 21-22 nt and 23-24 nt sRNAs. The Y-axis is scaled to inverse sine hyperbolic function of difference values normalised to H3.

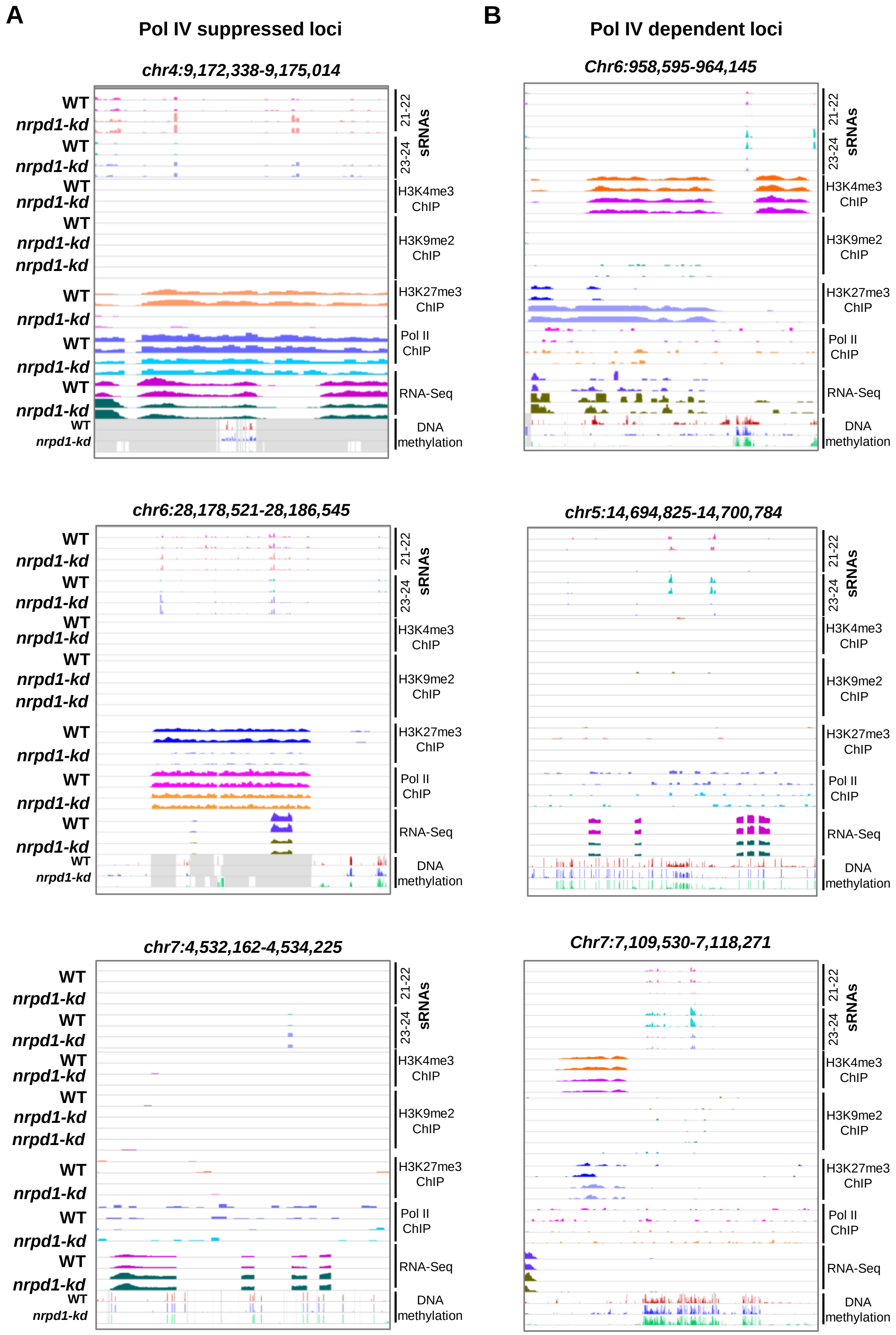


**Supplemental Figure S23.** Representative screenshots of epigenetic features at representative suppressed and dependent loci. *(A-B)* Genome browser screenshots showing the status of the different epigenetic marks in WT and *kd* lines of the representative pol IV suppressed *(A)* and dependent *(B)* loci.

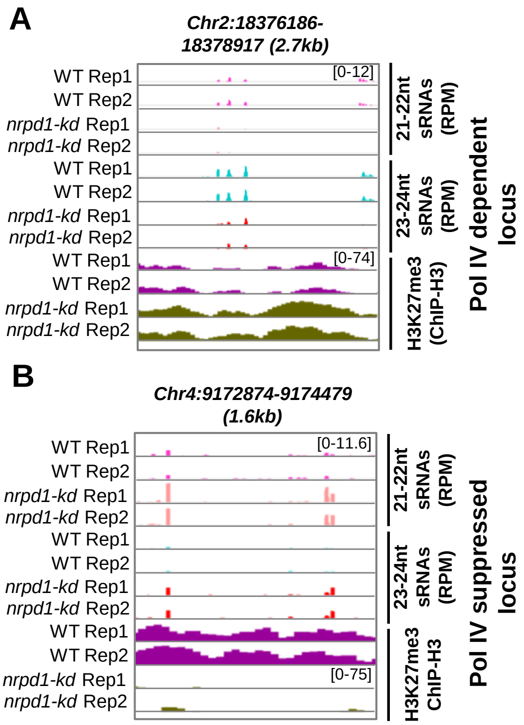


**Supplemental Figure S24**. Distinct H3K27me3 modifications at pol IV dependent and suppressed sRNA clusters. *(A-B)* Genome screenshots of representative pol IV dependent sRNA locus *(A)* and suppressed sRNA locus *(B)*.

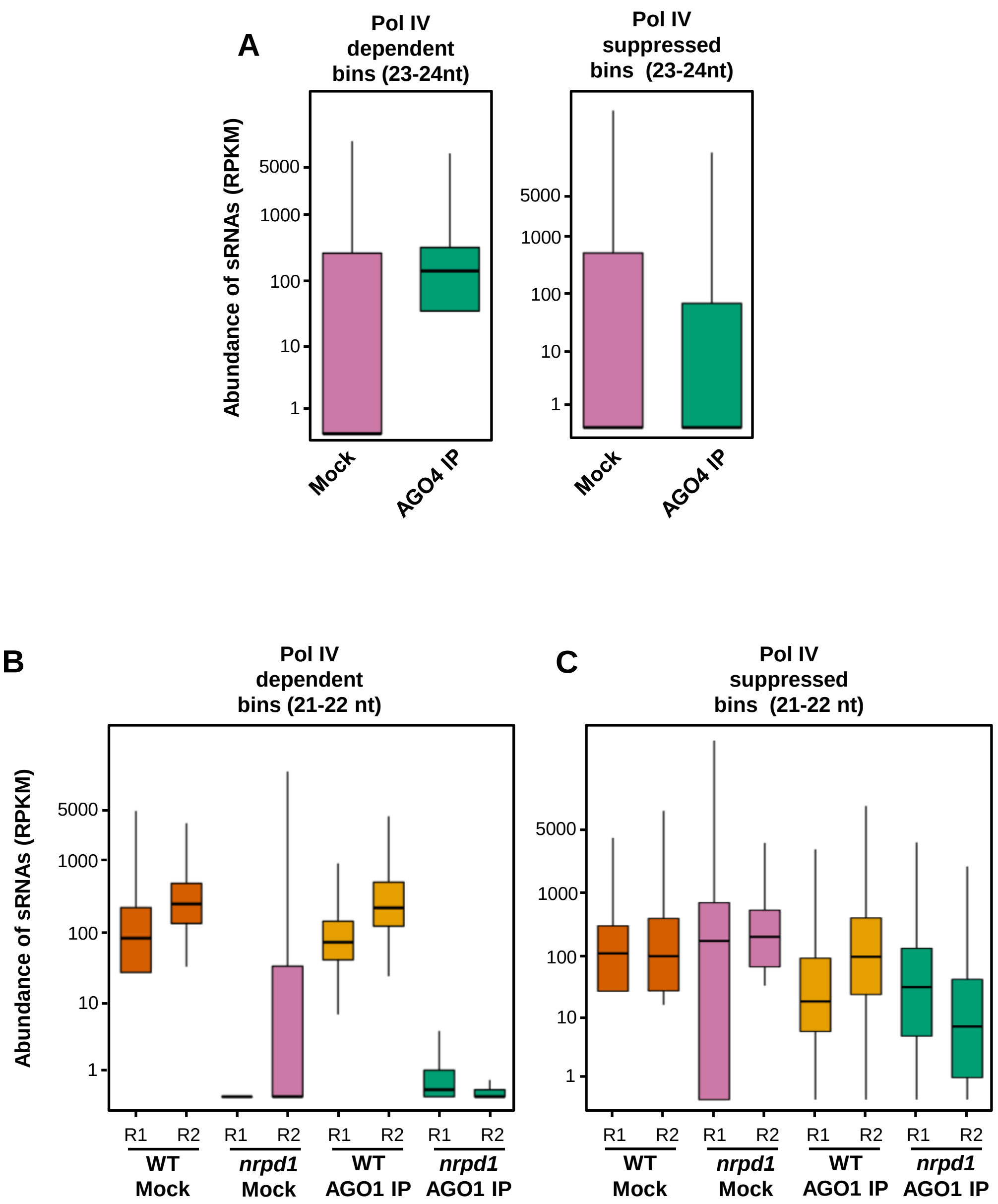


**Supplemental Figure S25.** Arabidopsis pol IV suppressed sRNAs are not effectively loaded into AGO1 and unlikely to target genes. *(A)* Stacked bar plots showing normalised abundance of small RNAs of different sizes from pol IV suppressed and dependent bins displaying the 5' nucleotide abundance in both the categories. *(B)* Box plots showing the accumulation of 21-22nt sRNAs categorized into pol IV dependent and suppressed sRNA bins from WT inflorescence in the AGO1 immuno-precipitate fraction. Datasets were taken from GSE61439. The Y-axis is scaled to inverse sine hyperbolic function of RPKM values. *(C)* Box plots showing the accumulation of 21-22nt sRNAs categorized into pol IV dependent and suppressed sRNAs from WT and *nrpd1* inflorescence in the AGO1 immuno-precipitate fraction. Datasets were taken from GSE133618. The Y-axis is scaled to inverse sine hyperbolic function of RPKM values.

**Transgenic plant generation and segregation analysis.** *Agrobacterium tumefaciens* strain LBA4404 with a helper plasmid pSB1 (containing extra copies of virulence genes) harbouring the binary vector of interest was used for infecting embryogenic calli derived from scutellum. Transformation of PB1 calli was performed as per previous methods (Hiei et al. 1994; Sridevi et al. 2003). About 21 days (d) old scutellar calli maintained on the callus induction media (CIM) (Murashige-Skoog (MS) medium, 2.5 mg/L 2,4-D, 300 mg/L casein hydrolysate, 500 mg/L L-proline, pH 5.8) was incubated for 4 d after cutting and infected with *Agrobacterium* culture of $OD_{600nm}$ around 1.0. The calli were co-cultivated with *Agrobacterium* in dark and washed with liquid CIM and then incubated in selection medium (CIM with 50 mg/L hygromycin or 5 mg/L phosphinothricin and 250 mg/L cefotaxime) for two rounds of 21 days each. The selected calli were moved to regeneration medium (MS medium, 3 mg/L kinetin, 2 mg/L NAA) in light. The regenerated shoots were moved to half-strength MS medium for rooting before hardening in growth chamber and then transferred to a transgene-compliant green house.

For segregation to subsequent generations, de-husked seeds were surface sterilised using ethanol, bleach and 0.1% mercuric chloride and placed on half-strength MS medium with 50mg/L hygromycin.

**Plant phenotyping.** WT and *kd* plants of different generations were grown uniformly after genotyping for the transgene. Tillers were counted on a per plant basis and the grain filling rate was calculated as number of filled grains per tiller.

Pollen staining and imaging was done as described in (Pedersen et al. 2004). Equally developed, pre dehisced anthers were collected and stained as described. Three biological replicates were chosen per transgenic line (2 transgenic lines - 6 plants) and multiple images were taken post staining under light microscope. The obtained images were analysed using custom ImageJ scripts for detecting the pollen staining and size of the pollen. Staining index was calculated as percentage of negative stain normalised to the background.

**Reverse Transcriptase – quantitative Polymerase Chain Reaction (RT-qPCR).** Total RNA isolated from the respective plant tissues using TRIzol method was converted to cDNA using reverse transcriptase kit (Thermo RevertAid RT kit) following manufacturer's protocol after treating with DNaseA. The cDNA template was used for the qPCRs using SYBR green qPCR master mix (Solis Biodyne - 5x HOT Firepol Evagreen qPCR Master Mix). Rice *Actin* (LOC_Os03g50885.1) was used as an internal control. The primers used for quantification are listed in Supplemental Table S3.

**mRNA northern hybridisation.** Total mRNA northern hybridisation was performed as described previously (Shivaprasad et al. 2006). Around 20ug of total RNA was ethanol precipitated and resuspended in denaturing loading buffer. RNA was electrophoresed in 1.5% denaturing agarose gel with 1% formaldehyde in MOPS buffer. The gel was blotted onto Hybond N+ membrane (GE healthcare) by capillary elution and UV crosslinked. The crosslinked membrane was hybridised using rRNA oligo probes described previously (Hang et al. 2018) end labelled with [γ-P32]-ATP in ultrahyb-buffer (Invitrogen). The blots are hybridised at 42$^{o}$C, washed, exposed to phosphor screen (GE healthcare) and scanned using Typhoon scanner. After scanning stripping and re-probing was performed as done for sRNA northern blots. The membrane was stained with methylene blue solution and shown as loading control.

**Extrachromosomal Circular DNA (ECC DNA) PCR.** Profiling of extrachromosomal fragments were performed as described previously (Lanciano et al. 2017). The primers were designed to amplify either only the circular DNA products or linear products. UBCE2 gene was used as loading control. To further validate the results, the total DNA (100ng) was digested with PlasmidSafe DNase (Lucigen) to eliminate the linear DNA and the same PCRs were repeated again. The primers used are listed in Supplemental Table S3.

**Scanning electron microscopy.** SEM imaging was performed as described previously (Das et al. 2020). Equally grown WT and *kd* spikelets were collected before anthesis and fixed in 0.1 M sodium phosphate buffer (pH 7.0) with 3% paraformaldehyde and 0.25%

glutaraldehyde. After incubation at 4°C for 24 h,spikelets were rinsed with 0.1 M sodium phosphate buffer and serially dehydrated with ethanol (30% to 100%). Samples were further dehydrated using critical point drying (LEICA EM CPD300), gold coated and the images were acquired using a Carl ZEISS scanning electron microscope at an accelerating voltage of 2 kV.

**Micro computed tomography (CT) scan.** Micro CT scanning of developing florets was performed as described earlier (Kannan et al. 2021). Florets from panicles that were equally developed before anthesis was scanned on Bruker Skyscan-1272 (Kontich, Belgium) at 40 kV, 250 µA, 4 µm image pixel size, without filters. The obtained image series was merged into a montage after colour adjustments using ImageJ.

**ChIP differential enrichment analyses.** ChIP differential enrichment analyses for the histone marks was performed using the tool Diffreps (Shen et al. 2013). The differentially enriched loci were identified using default parameters and the p-value cut-off of 0.001. The obtained loci were converted to bedfiles and the coverage of the ChIP signals normalised to H3 ChIP signal was obtained using bedtools multicov followed by library normalisation. The results were plotted using custom R scripts. Chromosome resolved heatmaps are generated using shinyChromosome (Yu et al. 2019).

**Nuclei immunostaining.** Immunofluorescence labelling and microscopy was performed as described earlier (Yelagandula et al. 2014) with the following modifications. The nuclei were isolated exactly as described for ChIP with fixation using 4% formaldehyde. The isolated nuclei are fixed again using 4% formaldehyde in PBS and taken for immunostaining performed in a tube. The post-fixed, washed nuclei are blocked with blocking solution (1xPBS with 0.1% Triton-X 100, 1%BSA, 10% normal goat serum and 100mM glycine) for 1 hour at room temperature. Primary antibody (H3K27me3 - Acive Motif 39155 [1:300]) incubations were done at 4 degrees for 16 hours in antibody binding solution (1xPBS with 0.1%BSA, 1% normal goat serum, 0.1% Triton-X 100 and 100mM glycine). The nuclei are washed with 1xPBS with 0.1% Triton X 100 and incubated with appropriate secondary antibodies (Invitrogen; goat raised Anti-mouse 488 and Anti-Rabbit 555) in antibody binding solution at 1:1000 dilution in dark at RT. Post washing, the nuclei are DAPI stained, mounted with fluoromount anti-fade reagent and imaged on a Confocal microscope (Zeiss LSM780). The images are analysed using ImageJ and JACoP (Bolte & F)was used to perform colocalization analysis.

**DNA methylome analyses.** Total DNA was isolated using CTAB method from the tissues mentioned. The library was constructed using the NEBNext® Enzymatic Methyl-seq Kit (Catalog no-E7120) using manufacturer's instructions and protocols described previously (Feng et al. 2020). The libraries were sequenced in a paired end mode (100bp) on a Hiseq2500 platform.

The obtained reads were quality checked and trimmed using cutadapt (Martin 2011) followed by alignment to IRGSP1.0 genome using Bismark aligner tool with default parameters (Krueger and Andrews 2011). DNA methylation status was extracted and coverage reports were generated using the Bismark tools. The obtained results are analysed using methylation package ViewBS (Huang et al. 2018). The chloroplast and mitochondrial genomes' cytosine conversion rate was used as a quality control. ViewBS tools were used for estimating DNA methylation in different contexts over the defined regions.

For Arabidopsis datasets (Supplemental Table S2), same analyses pipeline was used except that the reads are aligned to the TAIR10 genome.

**sRNA differential expression analyses.** The filtered sRNAs, size classified into 21-22nt and 23-24nt, were aligned and sRNA loci identified using Shortstack (Axtell 2013) with following parameters: --nohp --mmap f --mismatches 1 --mincov 0.2 rpmm. The raw counts obtained in each cluster was used for differential expression analyses using DESeq (Love et al. 2014). Only the clusters with p-value less than 0.05 and absolute $\log_2$ abundance fold change more than 1 were called as differential sRNA clusters (Supplemental Table S5). Volcano plots generated using custom R scripts. The clusters identified were further subcategorized into predominant 21-22nt clusters (if the 21-22nt population in a particular cluster exceeds 65% of total small RNAs), predominant 23-24nt (if 23-24nt sRNAs exceed 65%) and mixed clusters (all the other DE clusters).

**Genome window-based sRNA analyses and plotting.** The genome was split into 100bp non-overlapping windows using bedtools makewindows (Quinlan and Hall 2010) and the sRNAs (after dividing into 21-22nt and 23-24nt) were counted in each of the bins in the respective sized alignment files using bedtools multicov . The raw counts were normalised to RPM by considering the total reads mapped in each size class. All the bins that accumulate non-zero RPM values in both genotypes and at least minimum of 2 RPM when summed were taken for differential analysis. 2-fold change in RPM and absolute difference in RPM as minimum 5 were considered as the cut-off for differential calling for a bin (upregulated or downregulated in *kd* w.r.t. WT) (Supplemental Table S6). The bins identified from different tissues were taken for further analyses of mapping to genomic feature (overlap with the annotation), 5'-nucleotide bias of the sRNAs mapping to them, circos plots and the cumulative sum plots.

ChIPseekeR (Yu et al. 2015) was used for plotting the Vennpie diagrams of annotations of the suppressed and dependent sRNA bins after merging the different size category bins. Distal intergenic regions were the ones beyond 3 kb of any annotated PCGs and intergenic were the ones that are within this window. Circos plots displaying the heatmaps of the sRNA abundance of size-categorized bins were plotted using shinyCircos (Yu et al. 2018). The Upset plots were created in R using Intervene (Khan and Mathelier 2017).

Arabidopsis sRNA analysis was also performed the same way with the same thresholds for differential bin calling and associated analyses as handled for rice bins. The differential expressed sRNA bins were calculated with the same cut-off as used for rice when comparing WT and *nrpd1* sRNAs in different tissues (Supplemental Table S7). The chromosome-wide heatmaps of the sRNA bins were plotted using shinyChromosome (Yu et al. 2019). All the Arabidopsis datasets were processed the same way as rice and the bedtools multicov was used to estimate the sRNA abundance in each of the upregulated and downregulated sRNA bins. The data was plotted as a boxplot using ggplot in R. Statistical analyses are performed using R.

**Degradome analysis.** The obtained reads were processed for adapter removal and size filtering from 18 to 21 nt using UEA small RNA workbench (Stocks et al. 2018). CleaveLand version 4 (Addo-Quaye and Axtell 2008) with default parameters was used to identify the degradome validated target genes. Three fasta files were given as input to CleaveLand, sRNA from the pol IV suppressed loci, genes overlapping with pol IV suppressed loci and degradome reads. The cut-offs of Allen score 8 and mfe ratio of 0.65 were used to obtain the valid gene targeting list. The list of targeted genes is in Supplemental Table S8. The target sites are mapped onto the target transcripts and deeptools was used to generate the metaplots as described earlier centering at the slicing site.

**AGO-IP data analyses.** AGO IP datasets of various AGOs from rice and Arabidopsis were processed in the same way as the sRNA datasets, mapping to their corresponding genomes. The IP sRNAs mapping to the upregulated and downregulated bins/clusters were counted using bedtools multicov and box-plots were plotted using custom R scripts.

**RNA-seq and analyses.** RNA seq was performed in pre-dehisced anthers and pre-emerged panicle tissues. The total RNA was extracted using TRIzol method and was poly(A) enriched before library preparation. Library preparation was done with NEBNext® Ultra™ II Directional RNA Library Prep kit (E7765L) as per manufacturer's instructions. The obtained libraries were sequenced in paired end mode (100bp) on a Illumina Hiseq2500 platform.

The obtained reads were adapter trimmed using Trimmomatic (Bolger et al. 2014) and rRNA depletion was done using SortmeRNA (Kopylova et al. 2012). The reads are mapped to IRGSP1.0 genome using HISAT2 (Kim et al. 2015) with default parameters. Cufflinks (Trapnell et al. 2012) was used to perform differential gene expression analyses and statistical testing. The volcano plots were generated for DEGs using custom R scripts with the p-value cut-off of less than 0.05 and absolute $\log_2$ (fold change) expression cut-off of more than 2. For quantifying the expression of genes and transposons, bedtools multicov (Quinlan and Hall 2010) was used to obtain raw abundance and then normalised to RPKM values. These values are plotted as box-plots using custom R scripts.

**Supplemental Table S1 Details of high-throughput genomics data generated in this study**

| Sl. No. | Dataset type | Genotype | Replicate | Source tissue | SRA number | GSE number | Total number of reads obtained | Sequencing mode |
|---|---|---|---|---|---|---|---|---|
| 1 | Small RNA-seq | WT | Rep1 | Panicle | SRX11504325 | GSE180456 | 23329364 | Single end-50bp |
| 2 | Small RNA-seq | WT | Rep2 | | SRX11504326 | GSE180456 | 21698490 | Single end-50bp |
| 3 | Small RNA-seq | *nrpd1-kd* | Rep1 | | SRX11504327 | GSE180456 | 23575976 | Single end-50bp |
| 4 | Small RNA-seq | *nrpd1-kd* | Rep2 | | SRX11504328 | GSE180456 | 22447409 | Single end-50bp |
| 5 | Small RNA-seq | WT | Rep1 | Anther | SRX11504317 | GSE180456 | 29905295 | Single end-50bp |
| 6 | Small RNA-seq | WT | Rep2 | | SRX11504318 | GSE180456 | 27694239 | Single end-50bp |
| 7 | Small RNA-seq | *nrpd1-kd* | Rep1 | | SRX11504319 | GSE180456 | 28544601 | Single end-50bp |
| 8 | Small RNA-seq | *nrpd1-kd* | Rep2 | | SRX11504320 | GSE180456 | 27619824 | Single end-50bp |
| 9 | Small RNA-seq | WT | Rep1 | Endosperm | SRX11504321 | GSE180456 | 30944241 | Single end-50bp |
| 10 | Small RNA-seq | WT | Rep2 | | SRX11504322 | GSE180456 | 33043801 | Single end-50bp |
| 11 | Small RNA-seq | *nrpd1-kd* | Rep1 | | SRX11504323 | GSE180456 | 34105359 | Single end-50bp |
| 12 | Small RNA-seq | *nrpd1-kd* | Rep2 | | SRX11504324 | GSE180456 | 33446754 | Single end-50bp |
| 13 | Degradome-seq | WT | Rep1 | Panicle | SRX11490570 | GSE180312 | 79500184 | Single end-50bp |
| 14 | Degradome-seq | *nrpd1-kd* | Rep1 | | SRX11490571 | GSE180312 | 38231604 | Single end-50bp |
| 15 | Methylome-seq | WT | Rep1 | Panicle | SRX11492995 | GSE180333 | 32454221 pairs | Paired-end 100bp |
| 16 | Methylome-seq | *nrpd1-kd* | Rep1 | | SRX11492996 | GSE180333 | 46519465 pairs | Paired-end |

| | | | | | | | | |
|---|---|---|---|---|---|---|---|---|
| | | | | | | | | 100bp |
| 17 | Methylome-seq | *nrpd1-kd* | Rep2 | | SRX11492997 | GSE180333 | 46850796 pairs | Paired-end 100bp |
| 18 | H3K9me2_ChIP | WT | Rep1 | Panicle | SRX11490555 | GSE180311 | 49694617 | Single end-50bp |
| 19 | H3K9me2_ChIP | WT | Rep2 | | SRX11490556 | GSE180311 | 45286414 | Single end-50bp |
| 20 | H3K9me2_ChIP | *nrpd1-kd* | Rep1 | | SRX11490557 | GSE180311 | 39879848 | Single end-50bp |
| 21 | H3K9me2_ChIP | *nrpd1-kd* | Rep2 | | SRX11490558 | GSE180311 | 42994128 | Single end-50bp |
| 22 | H3K4me3_ChIP | WT | Rep1 | Panicle | SRX11490551 | GSE180311 | 50209378 | Single end-50bp |
| 23 | H3K4me3_ChIP | WT | Rep2 | | SRX11490552 | GSE180311 | 44933544 | Single end-50bp |
| 24 | H3K4me3_ChIP | *nrpd1-kd* | Rep1 | | SRX11490553 | GSE180311 | 36516201 | Single end-50bp |
| 25 | H3K4me3_ChIP | *nrpd1-kd* | Rep2 | | SRX11490554 | GSE180311 | 35517865 | Single end-50bp |
| 26 | H3K27me3_ChIP | WT | Rep1 | Panicle | SRX11490559 | GSE180311 | 47747901 | Single end-50bp |
| 27 | H3K27me3_ChIP | WT | Rep2 | | SRX11490560 | GSE180311 | 51446557 | Single end-50bp |
| 28 | H3K27me3_ChIP | *nrpd1-kd* | Rep1 | | SRX11490561 | GSE180311 | 52409180 | Single end-50bp |
| 29 | H3K27me3_ChIP | *nrpd1-kd* | Rep2 | | SRX11490562 | GSE180311 | 48057875 | Single end-50bp |
| 30 | H3_ChIP | WT | Rep1 | Panicle | SRX11490563 | GSE180311 | 33864148 | Single end-50bp |
| 31 | H3_ChIP | WT | Rep2 | | SRX11490564 | GSE180311 | 36276296 | Single end-50bp |
| 32 | H3_ChIP | *nrpd1-kd* | Rep1 | | SRX11490565 | GSE180311 | 33684346 | Single end-50bp |
| 33 | H3_ChIP | *nrpd1-kd* | Rep2 | | SRX11490566 | GSE180311 | 30225249 | Single end-50bp |
| 34 | PolII_ChIP | WT | Rep1 | Panicle | SRX11490567 | GSE180311 | 28985529 | Single end- |

| | | | | | | | | |
|---|---|---|---|---|---|---|---|---|
| | | | | | | | | 50bp |
| 35 | PolII_ChIP | *nrpd1-kd* | Rep1 | | SRX114 90568 | GSE180 311 | 357494 75 | Single end-50bp |
| 36 | INPUT_ch romatin | WT | Rep1 | | SRX114 90569 | GSE180 311 | 370389 43 | Single end-50bp |
| 37 | mRNA-seq | WT | Rep1 | Panicle | SRX114 93038 | GSE180 335 | 173213 26 pairs | Paired-end 100bp |
| 38 | mRNA-seq | WT | Rep2 | | SRX114 93039 | GSE180 335 | 173512 26 pairs | Paired-end 100bp |
| 39 | mRNA-seq | *nrpd1-kd* | Rep1 | | SRX114 93040 | GSE180 335 | 149268 95 pairs | Paired-end 100bp |
| 40 | mRNA-seq | *nrpd1-kd* | Rep2 | | SRX114 93041 | GSE180 335 | 160766 26 pairs | Paired-end 100bp |
| 41 | mRNA-seq | WT | Rep1 | Anther | SRX115 04317 | GSE180 335 | 386956 08 pairs | Paired-end 100bp |
| 42 | mRNA-seq | WT | Rep2 | | SRX115 04318 | GSE180 335 | 324552 82 pairs | Paired-end 100bp |
| 43 | mRNA-seq | *nrpd1-kd* | Rep1 | | SRX114 93044 | GSE180 335 | 292318 06 pairs | Paired-end 100bp |
| 44 | mRNA-seq | *nrpd1-kd* | Rep2 | | SRX114 93037 | GSE180 335 | 336894 53 pairs | Paired-end 100bp |
| 45 | H3K9me2 _ChIP | *nrpd1-kd* | Rep3 | Panicle | SRX135 86330 | GSE180 311 | | Single end-50bp |
| 46 | H3K9me2 _ChIP | *nrpd1-kd* | Rep4 | Panicle | SRX135 86331 | GSE180 311 | | Single end-50bp |
| 47 | PolII_ChIP | WT | Rep2 | Panicle | SRX135 86332 | GSE180 311 | | Single end-50bp |
| 48 | PolII_ChIP | *nrpd1-kd* | Rep2 | Panicle | SRX135 86333 | GSE180 311 | | Single end-50bp |
| 49 | Methylom e-seq | WT | Rep1 | Leaf | SRX135 86334 | GSE180 333 | | Paired-end 100bp |
| 50 | Methylom e-seq | *nrpd1-kd* | Rep1 | Leaf | SRX135 86335 | GSE180 333 | | Paired-end 100bp |

**Supplemental Table S2: Details of high-throughput genomics data obtained from publicly available datasets**

| Sl. No. | Species | Dataset type | Genotype | Source tissue | SRA number | GSE number | Reference |
|---|---|---|---|---|---|---|---|
| 1 | *Arabidopsis thaliana* | Small RNA-seq | Col-0 | 14d old seedlings | SRX2766992 | GSE98285 | (Yang et al. 2017) |
| 2 | *Arabidopsis thaliana* | Small RNA-seq | *nrpd1* | 14d old seedlings | SRX2766989 | GSE98285 | (Yang et al. 2017) |
| 3 | *Arabidopsis thaliana* | Small RNA-seq | *nrpe1* | 14d old seedlings | SRX2766990 | GSE98285 | (Yang et al. 2017) |
| 4 | *Arabidopsis thaliana* | Small RNA-seq | *nrpd1nrpe1* | 14d old seedlings | SRX1240828 | GSE72993 | (Zhang et al. 2016) |
| 5 | *Arabidopsis thaliana* | Small RNA-seq | Col-0 | Inflorescence | SRX1012515 | GSE61439 | (Zhai et al. 2015) |
| 6 | *Arabidopsis thaliana* | Small RNA-seq | *nrpd1* | Inflorescence | SRX1012518 | GSE61439 | (Zhai et al. 2015) |
| 7 | *Arabidopsis thaliana* | Small RNA-seq | Col-0 | Inflorescence | SRX1142679 | GSE61439 | (Zhai et al. 2015) |
| 8 | *Arabidopsis thaliana* | Small RNA-seq | Col-0 | Inflorescence | SRX1142680 | GSE61439 | (Zhai et al. 2015) |
| 9 | *Arabidopsis thaliana* | Small RNA-seq | *nrpb2* | Inflorescence | SRX1142681 | GSE61439 | (Zhai et al. 2015) |
| 10 | *Arabidopsis thalia* | Small RNA-seq | *nrpb2* | Inflorescence | SRX1142682 | GSE61439 | (Zhai et al. 2015) |

| | | | | | | | |
|---|---|---|---|---|---|---|---|
| | *na* | | | | | | |
| 11 | *Arabidopsis thaliana* | Small RNA-seq | *nrpd1dcl3* | Inflorescence | SRX1012521 | GSE61439 | (Zhai et al. 2015) |
| 12 | *Arabidopsis thaliana* | Small RNA-seq | *rdr2* | Inflorescence | SRX1012520 | GSE61439 | (Zhai et al. 2015) |
| 13 | *Arabidopsis thaliana* | Small RNA-seq | *rdr2dcl3* | Inflorescence | SRX1012522 | GSE61439 | (Zhai et al. 2015) |
| 14 | *Arabidopsis thaliana* | Small RNA-seq | *nrp(d/e)2* | Inflorescence | SRX1012519 | GSE61439 | (Zhai et al. 2015) |
| 15 | *Arabidopsis thaliana* | Small RNA-seq | Col-0 | Inflorescence | SRX7175631 | GSE140566 | (Li et al. 2020) |
| 16 | *Arabidopsis thaliana* | Small RNA-seq | *nrpd1* | Inflorescence | SRX7175632 | GSE140566 | (Li et al. 2020) |
| 17 | *Arabidopsis thaliana* | Small RNA-seq | 3xMyc_NRPD1_Complementation | Inflorescence | SRX7175633 | GSE140566 | (Li et al. 2020) |
| 18 | *Arabidopsis thaliana* | Small RNA-seq | 3xMyc_NRPD1_Complementation | Inflorescence | SRX7175634 | GSE140566 | (Li et al. 2020) |
| 19 | *Arabidopsis thaliana* | AGO1-IP | Col-0 | Inflorescence | SRX6385963 | GSE133618 | (Panda et al. 2020) |
| 20 | *Arabidopsis thaliana* | AGO1-IP | Col-0 | Inflorescence | SRX6385964 | GSE133618 | (Panda et al. 2020) |
| 21 | *Arabidopsis* | Mock-IP | Col-0 | Inflorescence | SRX6385965 | GSE133618 | (Panda et al. 2020) |

| | | | | | | | |
|---|---|---|---|---|---|---|---|
| | *thalia na* | | | | | | |
| 22 | *Arabi dopsi s thalia na* | Mock-IP | Col-0 | Infloresc ence | SRX638 5966 | GSE13 3618 | (Panda et al. 2020) |
| 23 | *Arabi dopsi s thalia na* | AGO1-IP | *nrpd1* | Infloresc ence | SRX638 5967 | GSE13 3618 | (Panda et al. 2020) |
| 24 | *Arabi dopsi s thalia na* | AGO1-IP | *nrpd1* | Infloresc ence | SRX638 5968 | GSE13 3618 | (Panda et al. 2020) |
| 25 | *Arabi dopsi s thalia na* | Mock-IP | *nrpd1* | Infloresc ence | SRX638 5969 | GSE13 3618 | (Panda et al. 2020) |
| 26 | *Arabi dopsi s thalia na* | Mock-IP | *nrpd1* | Infloresc ence | SRX638 5970 | GSE13 3618 | (Panda et al. 2020) |
| 27 | *Arabi dopsi s thalia na* | Mock-IP (for AGO4) | Col-0 | Infloresc ence | SRX114 2683 | GSE61 439 | (Zhai et al. 2015) |
| 28 | *Arabi dopsi s thalia na* | AGO4-IP | Col-0 | Infloresc ence | SRX114 2684 | GSE61 439 | (Zhai et al. 2015) |
| 29 | *Arabi dopsi s thalia na* | PolII - ChIP | Col-0 | Infloresc ence | SRX114 2672 | GSE61 439 | (Zhai et al. 2015) |
| 30 | *Arabi dopsi s thalia na* | PolII - ChIP | *nrpd1* | Infloresc ence | SRX114 2673 | GSE61 439 | (Zhai et al. 2015) |
| 31 | *Arabi dopsi s thalia na* | PolII - ChIP | *dcl234* | Infloresc ence | SRX114 2674 | GSE61 439 | (Zhai et al. 2015) |
| 32 | *Arabi dopsi* | ChIP-INPUT | Col-0 | Infloresc ence | SRX288 4373 | GSE99 694 | (Zhou et al. |

| | | | | | | | |
|---|---|---|---|---|---|---|---|
| | *s thaliana* | | | | | | 2018) |
| 33 | *Arabidopsis thaliana* | PolV-ChIP | Col-0 | Inflorescence | SRX2916399 | GSE100010 | (Liu et al. 2018) |
| 34 | *Arabidopsis thaliana* | Mock-IP | Col-0 | Inflorescence | SRX2916398 | GSE100010 | (Liu et al. 2018) |
| 35 | *Arabidopsis thaliana* | PolV-ChIP | *nrpd1* | Inflorescence | SRX2916401 | GSE100010 | (Liu et al. 2018) |
| 36 | *Arabidopsis thaliana* | Mock-IP | *nrpd1* | Inflorescence | SRX2916400 | GSE100010 | (Liu et al. 2018) |
| 37 | *Oryza sativa* | Small RNA-seq | WT-Nipponbare | Seedlings | SRX5724235 | GSE130166 | (Wang et al. 2020) |
| 38 | *Oryza sativa* | Small RNA-seq | WT-Nipponbare | Seedlings | SRX5724236 | GSE130166 | (Wang et al. 2020) |
| 39 | *Oryza sativa* | Small RNA-seq | *rdr2-6* | Seedlings | SRX5724237 | GSE130166 | (Wang et al. 2020) |
| 40 | *Oryza sativa* | Small RNA-seq | WT-Nipponbare | Panicle | SRX5724233 | GSE130166 | (Wang et al. 2020) |
| 41 | *Oryza sativa* | Small RNA-seq | *rdr2-6* | Panicle | SRX5724234 | GSE130166 | (Wang et al. 2020) |
| 42 | *Oryza sativa* | Small RNA-seq | *nrpd1* | Seedlings | SRX9211921 | GSE158709 | (Zheng et al. 2021) |
| 43 | *Oryza sativa* | Small RNA-seq | *nrpe1* | Seedlings | SRX9211922 | GSE158709 | (Zheng et al. 2021) |
| 44 | *Oryza sativ* | Small RNA-seq | *nrpe1* | Seedlings | SRX9211920 | GSE158709 | (Zheng et al. 2021) |

| | | | | | | | |
|---|---|---|---|---|---|---|---|
| | *a* | | | | | | |
| 45 | *Oryza sativa* | AGO1a-IP | WT | Seedlings | SRX014804 | GSE18250 | (Wu et al. 2009) |
| 46 | *Oryza sativa* | AGO1b-IP | WT | Seedlings | SRX014805 | GSE18250 | (Wu et al. 2009) |
| 47 | *Oryza sativa* | AGO1c-IP | WT | Seedlings | SRX014806 | GSE18250 | (Wu et al. 2009) |
| 48 | *Oryza sativa* | AGO4a-IP | WT | Seedlings | SRX017394 | GSE20748 | (Wu et al. 2010) |
| 49 | *Oryza sativa* | AGO4b-IP | WT | Seedlings | SRX017395 | GSE20748 | (Wu et al. 2010) |
| 50 | *Oryza sativa* | AGO16-IP | WT | Seedlings | SRX017396 | GSE20748 | (Wu et al. 2010) |
| 51 | *Oryza sativa* | Small RNA-seq | WT | Seedlings | SRX017400 | GSE20748 | (Wu et al. 2010) |
| 52 | *Arabidopsis thaliana* | Methylome | WT | Inflorescence | SRX2884303 | GSE99689 | (Zhou et al. 2018) |
| 53 | *Arabidopsis thaliana* | Methylome | *nrpd1* | Inflorescence | SRX2884312 | GSE99689 | (Zhou et al. 2018) |
| 54 | *Oryza sativa* | Small RNA-seq | WT Rep1 | Leaf | SRX11930857 | PRJNA758109 | (Chakraborty et al. 2021) |
| 55 | *Oryza sativa* | Small RNA-seq | WT Rep2 | Leaf | SRX11930858 | PRJNA758109 | (Chakraborty et al. 2021) |
| 56 | *Oryza sativa* | Small RNA-seq | WT Rep3 | Leaf | SRX11930859 | PRJNA758109 | (Chakraborty et al. 2021) |
| 57 | *Oryza sativ* | Small RNA-seq | nrp(d/e)2 Rep1 | Leaf | SRX11930854 | PRJNA758109 | (Chakraborty et al. |

| | | | | | | | |
|---|---|---|---|---|---|---|---|
| | *a* | | | | | | 2021) |
| 58 | *Oryza sativa* | Small RNA-seq | nrp(d/e)2 Rep2 | Leaf | SRX11930855 | PRJNA758109 | (Chakraborty et al. 2021) |
| 59 | *Oryza sativa* | Small RNA-seq | nrp(d/e)2 Rep3 | Leaf | SRX11930856 | PRJNA758109 | (Chakraborty et al. 2021) |
| 60 | *Arabidopsis thaliana* | Small RNA-seq | ago4ago6ago9 | Inflorescence | SRX11482427 | GSE165574 | (Sigman et al. 2021) |
| 61 | *Arabidopsis thaliana* | Small RNA-seq | drm1drm2_Rep1 | Inflorescence | SRX11482428 | GSE165574 | (Sigman et al. 2021) |
| 62 | *Arabidopsis thaliana* | Small RNA-seq | suvh2suvh9_Rep1 | Inflorescence | SRX11482429 | GSE165574 | (Sigman et al. 2021) |
| 63 | *Arabidopsis thaliana* | Small RNA-seq | suvh2suvh9_Rep2 | Inflorescence | SRX11482430 | GSE165574 | (Sigman et al. 2021) |
| 64 | *Arabidopsis thaliana* | Small RNA-seq | nrpe1 | Inflorescence | SRX252961 | GSE45368 | (Law et al. 2013) |
| 65 | *Arabidopsis thaliana* | Small RNA-seq | drm2 | Inflorescence | SRX252963 | GSE45368 | (Law et al. 2013) |
| 66 | *Arabidopsis thaliana* | Small RNA-seq | WT_Rep1 | Inflorescence | SRX2884335 | GSE99694 | (Zhou et al. 2018) |
| 67 | *Arabidopsis thaliana* | Small RNA-seq | WT_Rep2 | Inflorescence | SRX2884336 | GSE99694 | (Zhou et al. 2018) |
| 68 | *Arabidopsis thaliana* | Small RNA-seq | WT_Rep3 | Inflorescence | SRX2884337 | GSE99694 | (Zhou et al. 2018) |

| | | | | | | | |
|---|---|---|---|---|---|---|---|
| 69 | *Arabidopsis thaliana* | Small RNA-seq | clsy1_R1 | Inflorescence | SRX2884338 | GSE99694 | (Zhou et al. 2018) |
| 70 | *Arabidopsis thaliana* | Small RNA-seq | clsy2_R1 | Inflorescence | SRX2884339 | GSE99694 | (Zhou et al. 2018) |
| 71 | *Arabidopsis thaliana* | Small RNA-seq | clsy3_R1 | Inflorescence | SRX2884340 | GSE99694 | (Zhou et al. 2018) |
| 72 | *Arabidopsis thaliana* | Small RNA-seq | clsy4_R1 | Inflorescence | SRX2884341 | GSE99694 | (Zhou et al. 2018) |
| 73 | *Arabidopsis thaliana* | Small RNA-seq | clsy1_2_R1 | Inflorescence | SRX2884342 | GSE99694 | (Zhou et al. 2018) |
| 74 | *Arabidopsis thaliana* | Small RNA-seq | clsy1_3_R1 | Inflorescence | SRX2884343 | GSE99694 | (Zhou et al. 2018) |
| 75 | *Arabidopsis thaliana* | Small RNA-seq | clsy1_4_R1 | Inflorescence | SRX2884344 | GSE99694 | (Zhou et al. 2018) |
| 76 | *Arabidopsis thaliana* | Small RNA-seq | clsy2_3_R1 | Inflorescence | SRX2884345 | GSE99694 | (Zhou et al. 2018) |
| 77 | *Arabidopsis thaliana* | Small RNA-seq | clsy2_4_R1 | Inflorescence | SRX2884346 | GSE99694 | (Zhou et al. 2018) |
| 78 | *Arabidopsis thaliana* | Small RNA-seq | clsy3_4_R1 | Inflorescence | SRX2884347 | GSE99694 | (Zhou et al. 2018) |
| 79 | *Arabidopsis thalia* | Small RNA-seq | clsy1_2_3_4_R1 | Inflorescence | SRX2884348 | GSE99694 | (Zhou et al. 2018) |

| | | | | | | | |
|---|---|---|---|---|---|---|---|
| | *na* | | | | | | |
| 80 | *Arabidopsis thaliana* | Small RNA-seq | clsy1_2_3_4_R2 | Inflorescence | SRX2884349 | GSE99694 | (Zhou et al. 2018) |
| 81 | *Arabidopsis thaliana* | Small RNA-seq | shh1_R2 | Inflorescence | SRX2884350 | GSE99694 | (Zhou et al. 2018) |
| 82 | *Arabidopsis thaliana* | Small RNA-seq | shh1clsy1_R1 | Inflorescence | SRX2884351 | GSE99694 | (Zhou et al. 2018) |
| 83 | *Arabidopsis thaliana* | Small RNA-seq | shh1clsy1_2_R1 | Inflorescence | SRX2884352 | GSE99694 | (Zhou et al. 2018) |
| 84 | *Arabidopsis thaliana* | Small RNA-seq | shh1clsy3_4_R1 | Inflorescence | SRX2884353 | GSE99694 | (Zhou et al. 2018) |
| 85 | *Arabidopsis thaliana* | Small RNA-seq | nrpd1 | Inflorescence | SRX2884354 | GSE99694 | (Zhou et al. 2018) |
| 86 | *Arabidopsis thaliana* | Small RNA-seq | drm1_2_R2 | Inflorescence | SRX2884355 | GSE99694 | (Zhou et al. 2018) |
| 87 | *Arabidopsis thaliana* | Small RNA-seq | cmt2_R1 | Inflorescence | SRX2884356 | GSE99694 | (Zhou et al. 2018) |
| 88 | *Arabidopsis thaliana* | Small RNA-seq | cmt3_R1 | Inflorescence | SRX2884357 | GSE99694 | (Zhou et al. 2018) |
| 89 | *Arabidopsis thaliana* | Small RNA-seq | met1-3_rep1 | Inflorescence | SRX2884360 | GSE99694 | (Zhou et al. 2018) |
| 90 | *Arabidopsis* | Small RNA-seq | met1-3_rep2 | Inflorescence | SRX2884361 | GSE99694 | (Zhou et al. 2018) |

| | | | | | | | |
|---|---|---|---|---|---|---|---|
| | *thaliana* | | | | | | |
| 91 | *Arabidopsis thaliana* | Small RNA-seq | ddm1_R1 | Inflorescence | SRX2884362 | GSE99694 | (Zhou et al. 2018) |
| 92 | *Arabidopsis thaliana* | Small RNA-seq | atxr56_R1 | Inflorescence | SRX2884363 | GSE99694 | (Zhou et al. 2018) |
| 93 | *Arabidopsis thaliana* | Small RNA-seq | suvh4_5_6_R1 | Inflorescence | SRX2884364 | GSE99694 | (Zhou et al. 2018) |

### Supplemental Table S3: List of oligos and probes used in this study

| Oligo Name | Oligo ID | Oligo sequence (5' – 3') | Purpose | Reference |
|---|---|---|---|---|
| PolIV_miR-s I | 330 | agcaggagattcagtttga | amiR:NRPD1 construct | This study |
| PolIV_miR-a II | 331 | tgatgtccaagagtaacactatactgctgctgctacagcc | amiR:NRPD1 construct | This study |
| PolIV_miR*s III | 332 | ctatgtcgaagtgtaacactatattcctgctgctaggctg | amiR:NRPD1 construct | This study |
| PolIV_miR*a IV | 333 | aatatagtgttacacttcgacatagagaggcaaaagtgaa | amiR:NRPD1 construct | This study |
| OsNRPD1b_CDS_Fwd | 1859 | atggaggagccaagtcttgaggtgaaaatgcctgaagc | OsNRPD1b complementation construct | This study |
| OsNRPD1b_CDS_Rev | 1860 | ctacaactgatgtggccgaattgctctattagatagtatcttc | OsNRPD1b complementation construct | This study |
| OsNRPD1b_miRRES_F | 2665 | cgggttactttgtccaggagtaaacttcaccgtaac | OsNRPD1b complementation construct | This study |
| OsNRPD1b_miRRES_R | 2666 | gttacggtgaagtttactcctggacaaagtaacccg | OsNRPD1b complementation construct | This study |
| OsNRPD1b_Prom_F | 2191 | gagagtaaccaacacagaaacaggtgggccttg | OsNRPD1b complementation construct | This study |
| OsNRPD1_Prom_R | 2185 | cctgaacaataaacacgactagaaaacattaacatacatac aagtacc | OsNRPD1b complementation construct | This study |
| qPCR_NRPD1a&b _F | 1769 | ttgcaagtattgctcaaaggatgg | RT-qPCR and RT-PCR | This study |
| qPCR_NRPD1a&b _R | 1770 | ccaccagcaacttctgcgataattg | RT-PCR | This study |
| qPCR_NRPD1b_R | 3623 | ccccaccagcaacttctgcg | RT-qPCR | This study |
| qPCR_NRPD1a_R | 3622 | cttggacttatgtgccaccccg | RT-qPCR | This study |
| Tos17_TP_RT_F | 2061 | caccaggtgtggaagctccac | RT-qPCR and RT-PCR | (Nosaka et al. 2012) |
| Tos17_TP_RT_F | 2062 | taccactgagctgaagcgtgc | RT-qPCR and RT-PCR | (Nosaka et al. 2012) |
| HygR_F | 415 | aaagcctgaactcaccgc | RT-PCR and Southern probe | This study |
| HygR_R | 416 | ggtttccactatcggcga | RT-PCR and Southern probe | This study |
| BlpR_F | 340 | tcaaatctcggtgacgggcag | RT-PCR | This study |
| BlpR_R | 341 | atgagcccagaacgacgc | RT-PCR | This study |

| | | | | |
|---|---|---|---|---|
| OsActin1_F | 1786 | gctatgtacgtcgccatccagg | RT-qPCR and RT-PCR | This study |
| OsActin1_R | 1787 | tgagatcacgcccagcaagg | RT-qPCR and RT-PCR | This study |
| rRNA S7A and S7B (Region marked as 1) | 3478 and 3479 | tgttttggtcagggtcacgacaatgatcct and gcggtctgttttggtcagggtcacg | mRNA northern | (Hang et al. 2018) |
| rRNA p23 (Region marked as 2) | 3476 | gctgctcatacgctacgcagccac | mRNA northern | (Hang et al. 2018) |
| rRNA p42 (Region marked as 3) | 3480 | gcctcgcgcgcgagcgctcggcgggcaggggtga | mRNA northern | (Hang et al. 2018) |
| rRNA p22 (Region marked as 4) | 3477 | ctgtcgtcttccgagagcatct | mRNA northern | (Hang et al. 2018) |
| rRNA p4 (Region marked as 5) | 3481 | cgtgtggatttaactcgtggtatc | mRNA northern | (Hang et al. 2018) |
| SB_probe_Ubi_P_F | 2282 | ctgcagtgcagcgtgacccggtcg | Southern probe | This study |
| SB_probe_Ubi_P_R | 2283 | ctgcagaagtaacaccaaacaacag | Southern probe | This study |
| SB_probe_LINE1_F | 2344 | tctctggacgagcctgttccaa | Southern probe | (Cui et al. 2013) |
| SB_probe_LINE1_R | 2345 | ggctaagtcgtcagttgaatgc | Southern probe | (Cui et al. 2013) |
| Linear_UBCE2_F | 2943 | ccttcggagacaccttttgaagg | ECC DNA analyses | This study |
| Linear_UBCE2_R | 2944 | gtacgcagagaaggcaatgcag | ECC DNA analyses | This study |
| Linear_Tos17_F | 2229 | gctacccgttcttggactat | ECC DNA analyses | This study |
| Linear_Tos17_R | 2230 | ctgaaatcggagcactgaca | ECC DNA analyses | This study |
| Circular_Tos17_F | 2955 | aactcgagagcatcatcggttaca | ECC DNA analyses | (Lanciano et al. 2017) |
| Circular_Tos17_R | 2966 | cattagctgtatgaacggtggcac | ECC DNA analyses | (Lanciano et al. 2017) |
| Circular_PopRice_F | 2953 | acaaactgctgtcctaactgtcct | ECC DNA analyses | (Lanciano et al. 2017) |
| Circular_PopRice_R | 2954 | gcagctataaatatgtatccaatcct | ECC DNA analyses | (Lanciano et al. 2017) |
| Linear_PopRice_F | 2951 | gttatttcctgctgctcgtcgac | ECC DNA analyses | (Lanciano et al. 2017) |
| Linear_PopRice_R | 2952 | gtcgccgagaagatcctccatc | ECC DNA analyses | (Lanciano et al. 2017) |
| miRNA168 | 32 | gtcgccgagaagatcctccatc | sRNA northern | This study |
| U6_probes | 13 and 14 | ggccatgctaatcttctctgtatcgtt and ccaattttatcggatgtccccgaagggac | sRNA northern | This study |

| | | | | |
|---|---|---|---|---|
| amiR_PolIV_probe | 1697 | atgtccaagagtaacactata | sRNA northern | This study |
| miRNA444 | 245 | aagcttgaggcagcaactgca | sRNA northern | This study |
| miRNA156 | 3094 | gtgctcactctcttctgtcaa | sRNA northern | This study |
| MITE siRNA | 3430 | ggtcccacctgtcatacacacact | sRNA northern | This study |
| Tos17 siRNA | 2091 | ggcctctacatccaaccggatat | sRNA northern | This study |
| miRNA815 | 3345 | cccaatctcctcaatcccctt | sRNA northern | (Zhang et al. 2016) |
| miRNA397 | 511 | ggttcatcaacgctgcactcaa | sRNA northern | (Swetha et al. 2018) |
| miRNA820 | 3343 | cctggtccatccacgaggccga | sRNA northern | (Nosaka et al. 2012) |
| CACTA siRNA | 2086 | acgttcaatacaatctcggcgcgt | sRNA northern | This study |
| OsPolIV suppressed locus1 | 3318 | gctgacagaaagagaagtgagc | sRNA northern | This study |
| OsPolIV suppressed locus2 | 3319 | tatagtgttactcttggactt | sRNA northern | This study |
| miRNA845 | 3344 | ccgacaattggtatcagagca | sRNA northern | This study |
| Simplehat | 3676 | tgggttacccattttgacacccccta | sRNA northern | (Mosher et al. 2009) |
| AtRep2 | 3677 | gcgggacgggtttggcaggacgttacttaat | sRNA northern | (Mosher et al. 2009) |
| miR158 | 3620 | tgctttgtctacatttggga | sRNA northern | This study |
| AtPolIV suppressed locus 1 | 3736 | atttgtctaccgcatcattcatt | sRNA northern | This study |
| AtPolIV suppressed locus 2 | 3673 | attctaccactagactactggtgc | sRNA northern | This study |

Cui X, Jin P, Cui X, Gu L, Lu Z, Xue Y, Wei L, Qi J, Song X, Luo M, et al. 2013. Control of transposon activity by a histone H3K4 demethylase in rice. *Proc Natl Acad Sci U S A* **110**: 1953–1958.

Hang R, Wang Z, Deng X, Liu C, Yan B, Yang C, Song X, Mo B, Cao X. 2018. Ribosomal RNA biogenesis and its response to chilling stress in Oryza sativa. *Plant Physiol* **177**: 381–397.

Lanciano S, Carpentier M-C, Llauro C, Jobet E, Robakowska-Hyzorek D, Lasserre E, Ghesquière A, Panaud O, Mirouze M. 2017. Sequencing the extrachromosomal circular mobilome reveals retrotransposon activity in plants. *PLoS Genet* **13**: e1006630.

Mosher RA, Melnyk CW, Kelly KA, Dunn RM, Studholme DJ, Baulcombe DC. 2009. Uniparental expression of PolIV-dependent siRNAs in developing endosperm of Arabidopsis. *Nature* **460**: 283–286.

Nosaka M, Itoh J-I, Nagato Y, Ono A, Ishiwata A, Sato Y. 2012. Role of transposon-derived small RNAs in the interplay between genomes and parasitic DNA in rice. *PLoS Genet* **8**: e1002953.

Swetha C, Basu D, Pachamuthu K, Tirumalai V, Nair A, Prasad M, Shivaprasad PV. 2018. Major domestication-related phenotypes in Indica rice are due to loss of miRNA-mediated laccase silencing. *Plant Cell* **30**: 2649–2662.

Zhang H, Tao Z, Hong H, Chen Z, Wu C, Li X, Xiao J, Wang S. 2016. Transposon-derived small RNA is responsible for modified function of WRKY45 locus. *Nat Plants* **2**: 16016.

Supplemental_Table_S4_List_of_Arabidopsis_mutants_used_in_this_study

| **Sl. No.** | **Genotype** | **Genotype symbol used** | **Reference (or) source** |
|---|---|---|---|
| 1 | Col-0 | Col-0 | N/A |
| 2 | *nrpd1a-2* | *nrpd1* | (Herr et al. 2005) |
| 3 | *nrpd1a-1;rdr2-3* | *nrpd1 rdr2* | (Prof. David Baulcombe lab) |
| 4 | *dcl3-1* | *dcl3* | (SALK_005512) |
| 5 | *dcl2-1;dcl3-1;dcl4-2* | *dcl234* | (Henderson et al. 2006) |
| 6 | *ago4-4* | *ago4* | INRA FLAG_216G02 |
| 7 | *ddm1-2* | *ddm1* | (Jeddeloh et al. 1999) |


Henderson IR, Zhang X, Lu C, Johnson L, Meyers BC, Green PJ, Jacobsen SE. 2006. Dissecting Arabidopsis thaliana DICER function in small RNA processing, gene silencing and DNA methylation patterning. *Nat Genet* **38**: 721–725.

Herr AJ, Jensen MB, Dalmay T, Baulcombe DC. 2005. RNA polymerase IV directs silencing of endogenous DNA. *Science* **308**: 118–120.

Jeddeloh JA, Stokes TL, Richards EJ. 1999. Maintenance of genomic methylation requires a SWI2/SNF2-like protein. *Nat Genet* **22**: 94–97.
